# Biomolecular tracking by FIRESCAPE reveals distinct modes of clearance, damage induction and cellular uptake for extracellular histone H3

**DOI:** 10.1101/2025.08.13.667490

**Authors:** Brian Josephson, Adeline W. J. Poh, Yana Demyanenko, Abi. G. Yates, Joseph Ford, Andrew M. Giltrap, Tatsiana Auchynnikava, Kel Vin Tan, Isobel K. Dunstan, Weibing Liu, Adam P. Cribbs, Chen-Yi Wang, Martin Philpott, Sarah Able, Jeroen B. I. Sap, Alice Bedwell, Richard Aarnio, Alex M. Dickens, Jatta S. Helin, Luciana Kovacs, Noora A. Rajala, Maxwell W. G. Miner, Udo Oppermann, Katherine A. Vallis, Daniel R. McGowan, Anu J. Airaksinen, Veronique Gouverneur, Shabaz Mohammed, Daniel C. Anthony, Benjamin G. Davis

**Author notes:** These authors contributed equally: Brian Josephson, Adeline W. J. Poh (arranged alphabetically).

## Abstract

In attempting to observe the behaviour of a protein of interest *in vivo*, akin to other observer effects, current techniques require modifications or interventions that inherently alter the protein or its host, leaving one uncertain as to whether natural behaviour remains unperturbed. The study of chromatin has mostly been restricted to defining its function in the nucleus, where histone proteins fulfil vital roles in packaging genomic DNA and regulating transcription. However, chromatin components can be released into the extracellular space, either intentionally via cellular secretion or during disease-induced cell death. These extracellular chromatin components, depending on the context, can consist of: free histones, free DNA, intact nucleosomes (histone octamers wrapped by DNA), or heterogeneous, higher order structures such as neutrophil extracellular traps (NETs). They have been associated with diverse pathologies such as inflammation, cancer, and sepsis, and distinct toxic effects. However, there is a widely acknowledged lack of methods that distinguish between them and between their unique functions. Here, we now address protein observer effects to explore the fate and function of extracellular free histones by utilizing FIRESCAPE ([^18^F]-**<u>F</u>**luorine **<u>I</u>**sotopic **<u>R</u>**adiolabeling **<u>E</u>**nabling **<u>S</u>**canning of **<u>C</u>**learance **<u>A</u>**fter **<u>P</u>**roteolytic **<u>E</u>**vents), a novel radiolabeling concept that leverages the unique, high-sensitivity properties of the radioisotope fluorine-18, ^18^F, and residue-specific protein editing chemistry. By installing close and then ‘true’ ^18^F-containing protein sidechain mimics site-specifically, FIRESCAPE enables the hierarchical *in vivo* scanning of the half-lives, proteolytic susceptibility and clearance of single residues in a protein of interest, and at microdoses far below toxic levels (low nanomole). These ‘radioequivalent’ proteins bearing near-zero-size, zero-background labels now precisely reveal the strikingly distinct distribution, half-lives, damage-inducing abilities and accumulation of free extracellular histones *in cellulo* and *in vivo* compared to intact nucleosomes. Free extracellular histone H3 is rapidly cleared from circulation, mediated first by proteolysis of the histone tail. By contrast, direct injection of free histones vs nucleosomes into tissue that is unprotected by such proteolysis (brain), provokes a starkly different response; free histones exhibited limited diffusion and swiftly promoted damage both in cell culture and *in vivo*, whilst intact nucleosomes were essentially passive and benign. Remarkably, synthetic extracellular histone H3 was observed to enter cells and integrate into chromatin, indistinguishable from native H3 in both localization and post-translational modification (PTM) accumulation, yet paired with cellular and tissue damage. The exploratory studies described here now provide much needed clarity to the distinct fates and effects of extracellular histones vs nucleosomes, in particular the strongly damaging effects of free histones, their rapid uptake into cells, and an associated histone-specific proteolysis pathway via the removal of the histone tail.

## Introduction

Over the past half century, powerful studies have revealed the critical roles of histones in binding DNA and regulating transcription in the nucleus, where they exist as DNA-wrapped octamers within the smallest subunit of chromatin, the nucleosome^1,2^. When histones are expressed in the cytosol, they are immediately bound by chaperones and transported into the nucleus^3^, where their role in controlling gene expression through regulating chromatin accessibility and accumulating PTMs is a field of intense activity^4,5^. However, rare studies have also questioned possible functions of extracellular histones that are released after cell death due to injury or disease^6–8^ and during cellular secretion^9^. Some reports are conflicting about exactly which types of extracellular bodies contain histones, such as exosomes^10,11^, while other reports highlight the difficulty in distinguishing between non-vesicle bound “free” histones and nucleosomal histones in circulation^7^. Regardless, extracellular histones have intriguing and apparently surprising suggested activities for a protein whose function is canonically restricted; these include apparent activation of the immune system, leading even to inflammation and cell death^6–8^.

Extracellular free histones, cell free nucleosomes and associated DNA also have potential as surrogate biomarkers for diseases, such as cancer (**Extended Data** Figure 1). Nucleosome footprinting of cell free DNA^12^ and the detection of residual histone PTMs on cell free nucleosomes^13^ have been leveraged to infer the cells, tissues or tumours of origin that release these various chromatin components upon cell death (**Extended Data** Figure 1). However, the downstream (potentially damaging) effects of extracellular histones may be more directly relevant to health than the indirect information they carry about their origin prior to release. Cell free nucleosomes are suggested to be far less toxic than free histones^7,14^; while sepsis and lethality were observed in mice injected with a high dose of free histone (50-75 mg/kg)^15^, no immediate toxicity was detected with similar amounts of nucleosomes^16^. The reason for this discrepancy in toxicity remains unclear. More fundamentally, the distribution, stability, half-lives and even fates of extracellular histones and nucleosomes in circulation remain essentially undefined.

Here, we introduce and implement a novel protein radiolabeling concept, termed FIRESCAPE (**Figure 1**), which goes beyond conventional radiolabeling strategies by enabling not only tracking of near-native protein fate but simultaneous domain- and residue-specific processing *in vivo* via time-resolved positron emission tomography (PET) imaging. FIRESCAPE accomplishes this via subtle, near-zero-size H-to-^18^F editing within the sidechains of predetermined ‘scanned’ protein residues coupled with deconvolutive, characteristic pharmacokinetics.

**Figure 1:**
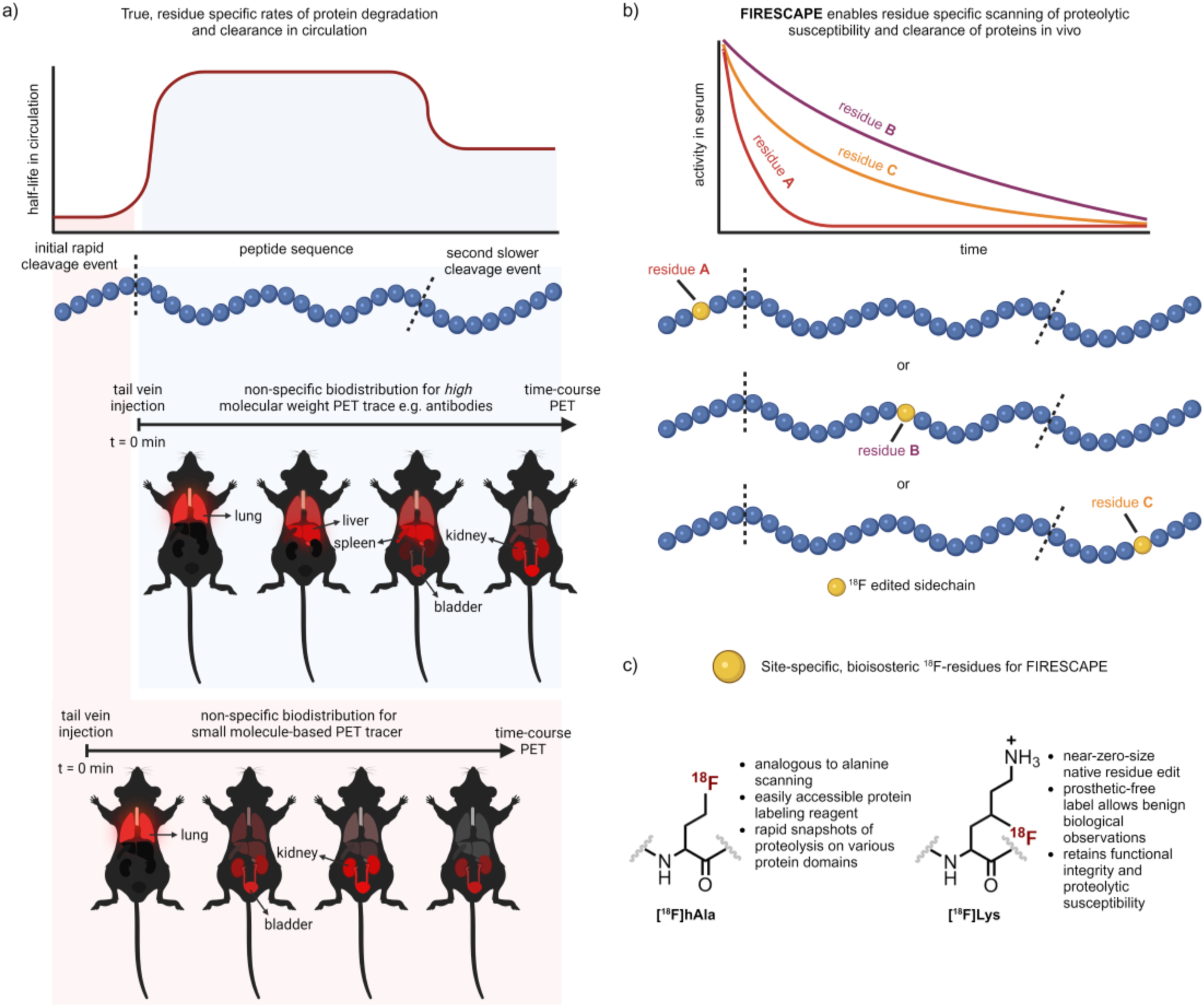
The concept of FIRESCAPE allows dynamic monitoring of protein distribution, residue-specific susceptibility to proteolysis, and clearance kinetics. a) Proteolytic processing *in vivo* can result in varying biodistribution profiles for a particular protein species, depending on the nature of the cleaved product, such as whether it behaves as a small molecule or as a larger biomolecule. b) By site-specifically introducing a minimally perturbative radioactive label, it should be possible to map the *in vivo* stability of residues across the entire protein length as a proxy for simultaneous determination of domain-dependent functions, including processing, clearance kinetics as well as cellular uptake. c) [^18^F]hAla and [^18^F]Lys were designed and edited into proteins of interest as minimally-altered and/or radio-equivalent scanning residues to facilitate step-wise FIRESCAPE (see also Figure 2a) and provide genuine insights into the behaviour of target proteins *in vivo*.

FIRESCAPE-mediated domain and site scanning now pinpoints an *N*-terminal-tail site in extracellular histone H3 that is rapidly cleaved *in vivo* in proteolytic connective tissue. Apparent protection afforded by this proteolysis is indicated by correspondingly robust cellular damage response to extracellular histones in non-proteolytic tissues and cells, which contrasts with minimal effects of extracellular, intact nucleosomes. Finally, FIRESCAPE-determined localization of ultimate cellular destination in various tissues revealed apparent uptake of extracellular histones into proximal cells both *in cellulo* and *in vivo*. Characterization of the penetration of synthetic extracellular histones into cells confirmed not only uptake but also their deposition into chromatin in a manner that is functionally indistinguishable from endogenous histone H3, both in genomic localization and PTM accumulation.

## Results

### Design of FIRESCAPE, a method for precise tracking of protein fate in circulation

PET imaging is a long-standing tool for tracking radiolabelled small molecules, revealing features of metabolic processes and non-invasively mapping diseased tissues in patients^17^. In recent years, efforts have increasingly focused on radiolabelling proteins, primarily antibodies, that can also bind epitopic sites of disease^18^. A potentially valuable but largely under-used^19^, distinguishing metric of biomolecules in physiology is their characteristic circulation half-life and the associated kinetics of biodistribution, including clearance. For proteins, this can be further modulated by modes of degradation (proteolysis) into smaller peptides and even amino acid components. Moreover, the process of glomerular filtration distinguishes between biomolecules in circulation based on their size; those with larger Stokes radius (and so higher molecular mass) are retained with greater efficiency.^20,21^ Taking these together, we reasoned that if smaller fragments of proteins were to be released as peptides and amino acids, accumulating in a differential manner in liver or kidney (and then bladder)^22^, these might be readily distinguishable via time-resolved PET imaging. We considered that in turn this might serve as a putative strategy for the ‘scanning’ or ‘mapping’ of the *in vivo* susceptibility of protein domains to proteolysis in a general manner based on their release as fragments with consequently drastically-altered pharmacokinetics (**Figure 1a**).

Nearly all current protein (radio)labelling strategies use bulky linkers and/or prosthetic groups to attach radioisotopes, typically in a non-site-specific manner, to uncontrolled protein sidechains^23–25^ (**Extended Data** Figure 3a). These can mask or block protease recognition sites and, at best, can reveal only an artificial or incomplete picture of the protein of interest (**Extended Data** Figure 3b). We reasoned that a true reporter of *in vivo* protein fate would allow for the entire length of the protein of interest to be assessed. In a concept partially analogous to so-called alanine scanning^26^, a minimal radiolabel edited into the side chain of native protein residues could be installed into various residue positions within a protein of interest, allowing for single amino acid specificity in assessing half-life and clearance along the peptide chain. The editing of H→^18^F in chosen sidechains (**Extended Data** Figure 2) would not only allow for minimal (‘near-zero’) size variation,^27^ fluorine-18 is also a highly efficient (97%) positron emitter^28^ with a half-life of ∼110 min that can be administered in doses down to picomolar levels whilst still allowing images of high resolution. This could enable superior sensitivity of detection potentially critical in the *in vivo* study of proteins of interest at low abundance or concentration or that are potentially toxic at higher concentrations (such as extracellular histone).

This [^18^F]**<u>F</u>**luorine **<u>I</u>**sotopic **<u>R</u>**adiolabeling **<u>E</u>**nabling **<u>S</u>**canning of **<u>C</u>**learance **<u>A</u>**fter **<u>P</u>**roteolytic **<u>E</u>**vents (**FIRESCAPE**) strategy (**Figure 1**) deploys post-translational, near-zero-size H-to-^18^F substitution within protein residue sidechains to generate two types (**Figure 1c**) of FIRESCAPE reporter amino acid sites. First, a minimal radiolabeled alanine residue, [^18^F]hAla,^27^ is installed at various sites along a protein to “scan” proteolytic susceptibility by domain (**Figure 1b**) using clearance kinetics of specific protein domains via *in vivo* time-resolved PET imaging. Second, through the precise editing of an ^18^F-isotope into the sidechain of one of the primary, specific residues (so-called P_1_ or P_1_′, using the nomenclature of Schechter and Berger^29^) that are targeted^30^ for proteolytic cleavage, lysine (Lys), we generate [^18^F]Lys to access true ‘radio-equivalent’ protein mimics. In these ‘radio-equivalent’ proteins the only perturbation from the native connectivity is the substitution of one hydrogen atom in the sidechain of a pre-determined residue for fluorine-18 (**Figure 1c, right**). This in principle could enable essentially native proteolytic processing (and so highly precise residue-by-residue mapping) even when the radiolabel is found at these pivotal, active site-engaged P_1_ or P_1_′ residues.

### FIRESCAPE in connective tissue reveals protein domain-associated proteolysis in vivo

Implementing FIRESCAPE, we began “scanning” the length of the histone protein by first installing the minimal residue [^18^F]hAla into each of the two primary domains of the canonical human histone H3 (see **Supplementary Note 1** for construct details): the disordered *N*-terminal tail (at site Lys4), or within the folded core (at site Lys56) (**Figure 2a,b**). Synthetic, precisely-labelled histone H3 variants, H3-[^18^F]hAla4 and H3-[^18^F]hAla56 were readily created through post-translational editing (see **Extended Data** Figure 2 for further methodoligical details),^31^ exploiting the rapid generation of appropriate ^18^F-containing carbon-centred (C•) H_2_FC• radicals from [^18^F]btSOOCH_2_F (see **Extended Data** Figure 3c)^27^ and their selectivity and demonstrated compatibility^32^ for C–C bond formation on previously installed dehydroalanine (Dha) residues in proteins (**Supplementary Information).** This was critically accomplished efficiently (radiochemical yield (RCY) = 67 ± 5% (n = 2) and 49%, respectively, with 16%-24% isolated activity yields (non-decay-corrected, n.d.c) starting from purified [^18^F]btSOOCH_2_F, in a >99% radiochemical purity) and in a timescale (<180 min via manual radiosynthesis starting from azeotropically dried [^18^F]fluoride) compatible with the half-life of ^18^F (**Supplementary Information** and **Supplementary Note 5**).

**Figure 2:**
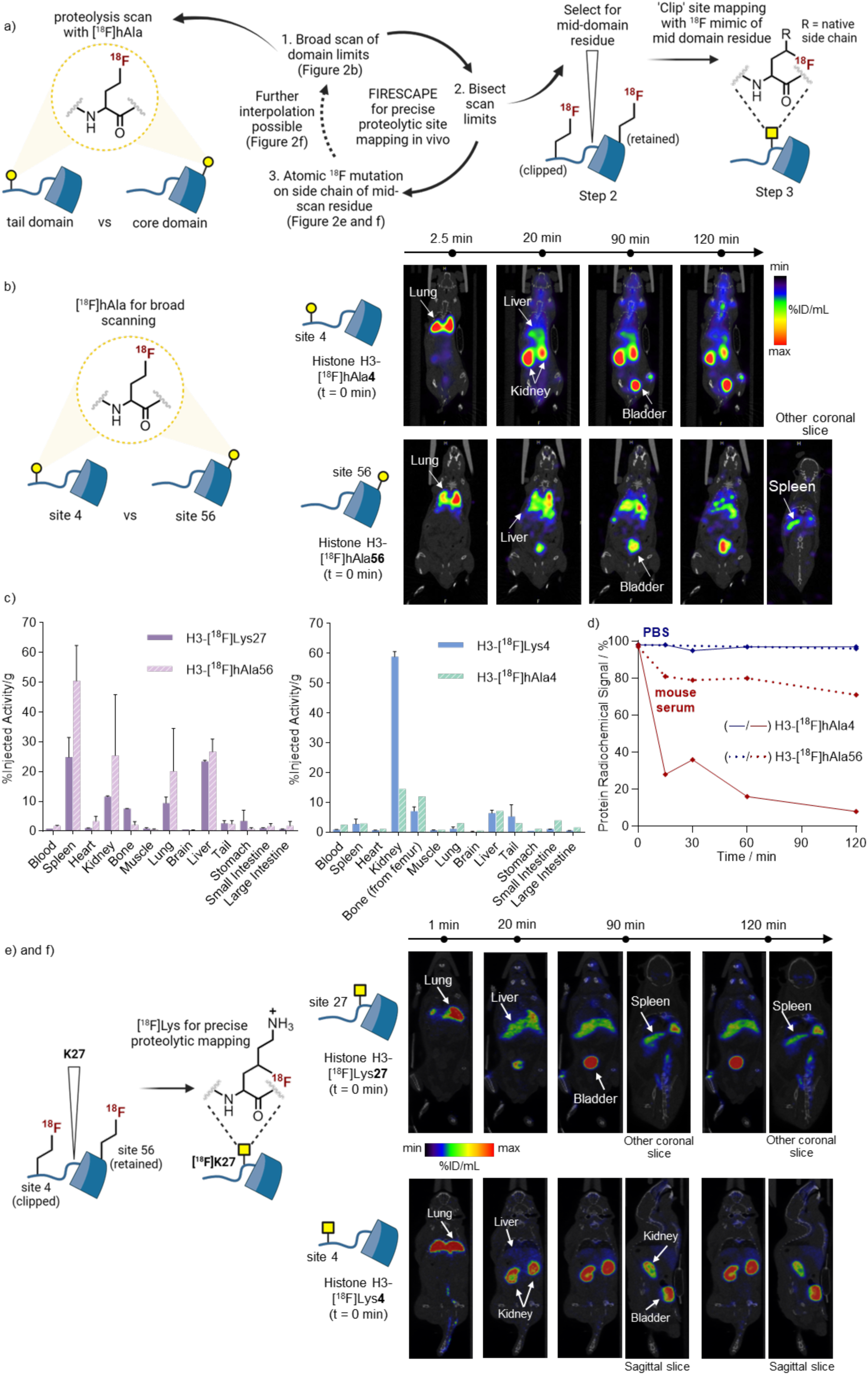
FIRESCAPE in connective tissue enables *in vivo* histone tracking via PET and reveals distinct clearance kinetics and histone specific proteolysis. Selected time-resolved PET images have been overlayed on CT scans. **a)** Multistage FIRESCAPE implemented on histone H3. FIRESCAPE begins with the installation of [^18^F]hAla as a broad proteolysis ‘scanner’ across protein domains (Step 1). Through differences in biodistribution and kinetic profiles *in vivo* (observed by PET) this Stage 1 [^18^F]hAla domain-scanning then informs Stage 2 [^18^F]Lys site-mapping via subsequent iterative Steps 2,3 etc. Selection of a bifurcating, mid-domain residue (Step 2) and then installation of a ^18^F side-chain radio-equivalent and subsequent PET analysis (Step 3) can be accompanied by further iterations to narrow down potential proteolytic site(s) *in vivo*. **b)** [^18^F]hAla-containing histone H3 showed distinct differences in clearance pathway between tail-labelled (at K4, top row) vs core-labelled (at site 56, bottom row) domain of circulating histone H3. c) *Ex vivo* organ accumulation analysis 2 h post-i.v. injection of ^18^F-labelled histone H3 proteins. Histones H3-[^18^F]hAla56 and H3-[^18^F]Lys27 display organ-uptake profiles characteristic of larger biomolecules (left), whilst histones H3-[^18^F]hAla4 and H3-[^18^F]Lys4 show highly predominant localisation in the kidneys, characteristic of smaller molecules. n=2 for H3-[^18^F]Lys27, [^18^F]hAla56, [^18^F]Lys4; n = 1 for H3-[^18^F]hAla4. **d)** Radio-HPLC analysis of [^18^F]hAla-labelled histone H3 revealed rapid decrease in the radiochemical signal of tail-labelled (site 4) vs core-labelled (site 56) histone in serum (n = 1, see also **Extended Data** Figure 4). **e) and f)** Radical precursor reagent [^18^F]btSOOLys (**Extended Data** Figure 5) enabled the installation of [^18^F]Lys in the mid-domain region to create H3-[^18^F]Lys27; Creation of this ‘radio-equivalent’ protein, in which the γ-H atom on the side chain of WT K27 is substituted for a γ-^18^F, enabled precise mapping of site susceptibility to proteolysis in Stage 2 FIRESCAPE. **e)** Retention of the radioisotope in the core of histone H3-[^18^F]Lys27 resulted in predominant localization in the liver and spleen (in addition to the bladder), while **f)** its removal as part of a smaller peptidic fragment released from histone H3-[^18^F]Lys4 led to fast signal concentration in the kidney and bladder.

Pharmacokinetics (**Figure 2b**) following parenteral administration (lateral tail vein in PBS, 2-10 MBq) of tail-labelled H3-[^18^F]hAla4) or core-labelled H3-[^18^F]hAla56) were analyzed via combined and co-registered PET-CT (**Extended Data** Figure 4b, images every 5 min). This dynamic analysis showed, on one hand, clearance (t_1/2_ ∼ 6.6 min, n = 1, see **Supplementary Note 4**) of the ^18^F-associated signal from H3-[^18^F]hAla4 to the kidney and then bladder. By contrast, core-labelled histone H3-[^18^F]hAla56 displayed longer half-life in circulation (t_1/2_ ∼ 15.2 min, n = 1) and a starkly different ^18^F-associated signal biodistribution pattern associated with high ^18^F-uptake into the spleen and liver, followed by later and lower transit to kidney and bladder (**Extended Data** Figure 4c). These differences were further confirmed by organ-specific *ex vivo* measurement of radioactivity (% injected activity per g, %IA.g^-1^) (**Figure 2c**). Consistent with the design of FIRESCAPE, these distinct biodistribution kinetics revealed very different glomerular filtration profiles, indicative of highly altered ^18^F-associated biomolecule masses^20,21^ precisely probed here through the administration of H3 histone variants with near-identical sequences (differing only in the position of [^18^F]hAla residue in tail or core domains). Time-course *ex vivo* radio-liquid chromatographic analyses (**Figure 2e**) further confirmed these altered profiles (H3-[^18^F]hAla56, 81% compared to H3-[^18^F]hAla4, 28%, after 15 min, **Figure 2d**).

Together, these strongly contrasting dynamic biodistribution data revealed *N*-terminal-tail domain loss as a previously unrealized processing of histone H3 in mammalian circulation. They also vitally confirmed the core scanning methodology of FIRESCAPE. Notably, 1) highly efficient H→^18^F editing could be achieved with sufficient molar activity^27^ to allow FIRESCAPE to be implemented via serial imaging at doses as low as 10 nmol of histone; 2) this in turn enabled time-resolved profiling through the high sensitivity of ^18^F-PET; and, 3) this used protein doses low enough to avoid triggering of undesired toxicity or immune responses^15^.

### FIRESCAPE precisely maps an in vivo ‘clip’ site for histone H3 in proteolytic tissue

Having discovered the apparent *domains* of *in vivo* processing for histone H3 we next used FIRESCAPE to map the precise *site* of processing. To validate the site-specific susceptibility of free extracellular histone H3 to proteolysis in circulation following our broad [^18^F]hAla scan, the second step of the FIRESCAPE design strategy (**Figure 1, 2a, 2c**) called for the generation of a ‘radio-equivalent protein’ through the single atom transformation of one H atom within a given residue sidechain on wildtype (WT) histone to ^18^F. Given the prevalence of Lys as a target protease residue^30^ (see above) and also within the H3 *N*-terminal domain (Lys4, Lys9, Lys14, Lys18, Lys23, <u>Lys27</u>, Lys36, Lys37, Lys56) scanning with [^18^F]Lys residues was selected – the bifurcating, mid-domain Lys27 site was chosen first (**Figure 2a**).

This construction of a radio-equivalent histone histone H3-[^18^F]Lys27, in turn, necessitated development of an expedient radiosynthesis of another corresponding C• radical precursor reagent: [^18^F]btSOOLys (**Extended Data** Figure 5a). This stage-two mode of FIRESCAPE therefore utilized a residue-specific (here [^18^F]Lys) variant of the generic, domain-scanning [^18^F]hAla reagent used in stage-one FIRESCAPE activated under blue light irradiation. Positron-emitting lysine sidechain [^18^F]Lys27 was installed instead of Lys27 into WT histone H3 using a pre-positioned Dha site. Thus, using a combined genetic (Lys27(WT) → Cys27) and chemical (Cys27 → Dha27 → [^18^F]Lys27) mutagenesis sequence, we were able to efficiently recapitulate the wild-type sequence in histone H3 whilst simultaneously introducing ^18^F by installing a radioactive analogue sidechain (16% isolated activity yield (n.d.c.) starting from purified [^18^F]btSOOLys, 170 min overall radiosynthesis time starting from cyclotron-produced [^18^F]fluoride, >98% radiochemical purity of ^18^F-labelled H3). This histone H3-Lys27→[^18^F]Lys27 alteration is to our knowledge the first example of a ‘radioequivalent protein’ (**Extended Data** Figure 5b**, 5c and Supplementary Information**).

Indicatively, intravenous (iv) administration of histone H3-[^18^F]Lys27 (∼4 MBq, ∼190-200 μg) and subsequent dynamic PET showed predominant ^18^F-accumulation primarily in the spleen and liver 2 h post-injection (**Figure 2e and Extended Data** Figure 5c) essentially similar to that previously observed in stage-one FIRESCAPE analysis with H3-[^18^F]hAla56 (see above). These observed similarities in biodistribution (**Figure 2e**) and organ uptake profiles (**Figure 2c**) suggested that rather than the bifurcating [^18^F]Lys27 residue being removed it instead remained after ‘clipping’ as part of the histone H3 core protein; this implicated a clipping site towards the *N*-terminus.

Next, therefore, we constructed a second ‘radio-equivalent’ protein of WT histone as the next step of an iterative FIRESCAPE multi-stage residue-specific scan (**Figure 2a**). By editing Lys4 to create histone H3-[^18^F]Lys4, the radioactive scan site was now positioned at the Lys site (Lys4) that is closest to the *N*-terminus of the histone H3 tail (6% isolated yield (n.d.c.) starting from purified [^18^F]btSOOLys, 175 min overall radiosynthesis time starting from cyclotron- produced [^18^F]fluoride, >99% radiochemical purity of ^18^F-labelled-H3). In striking contrast to the signal observed for H3-[^18^F]Lys27, the ^18^F-signal from H3-[^18^F]Lys4 (injected dose ∼ 150 μg) rapidly localized in the kidney and bladder (**Figure 2f and Extended Data** Figure 6), reflecting a clearance pathway that is distinct from the histone core (**Extended Data** Figure 6) and thereby unambiguously locating the *in vivo* clip site towards the *N*-terminus of Lys27 in residues 5-26.

### Histone H3 in non-proteolytic tissue and cell culture is unprotected from damage

Following our observation of rapid and tail specific proteolysis and clearance of histone H3 in circulation, we next considered the fate of extracellular histones in tissues that are largely shielded from proteases. The blood-brain barrier provides a robust buffer from proteases in circulation^33^ and excludes neutrophils from the brain under normal conditions^34,35^. The brain is therefore a tissue free from proteases and also free from contamination from the histones found in NETs (see below).

Injection of microdoses of WT histone H3 into brain parenchyma at levels intended to mimic low-level release in damaged tissues^8,15,36^ and far below lethal^15^ (**Figure 3a**, 1 µL of 55 µM histone H3 formulated with H4 1:1 for solubility, **see Supplementary Note 2**, in PBS, around 1000-fold below toxic levels) using a finely drawn capillary^37^ via an atraumatic procedure^38^ accordingly did not elicit any overt clinical signs of distress or disability in animals (and no abnormal histology was observed in any of the other tissues of animals treated via either route of parenteral administration). However, histology following euthanasia surprisingly identified evidence of focal neuronal cell death and inflammatory response proximal to the site of the injection (identified in fixed brain tissue slices, **Figure 3a**). Detailed immunohistochemistry revealed limited histone diffusion within tissue, with focal activation of microglia and astrocytes in the tissue immediately adjacent to the area of cell death (evident as early as 30 minutes after the injection of the histone H3). Very few neutrophils were recruited in response to the cell death, consistent with the anticipated protection of the brain parenchyma by the blood-brain barrier^35^. Notably, by contrast, comparator samples, where synthetic histone H3 was assembled into intact nucleosome core particles prior to administration (see **Supplementary Methods**), exhibited minimal damage (**Figure 3a**). Corresponding protein was found widely diffused throughout the brain slice (**Figure 3a**) and there was an absence of any appreciable neuronal cell death.

**Figure 3:**
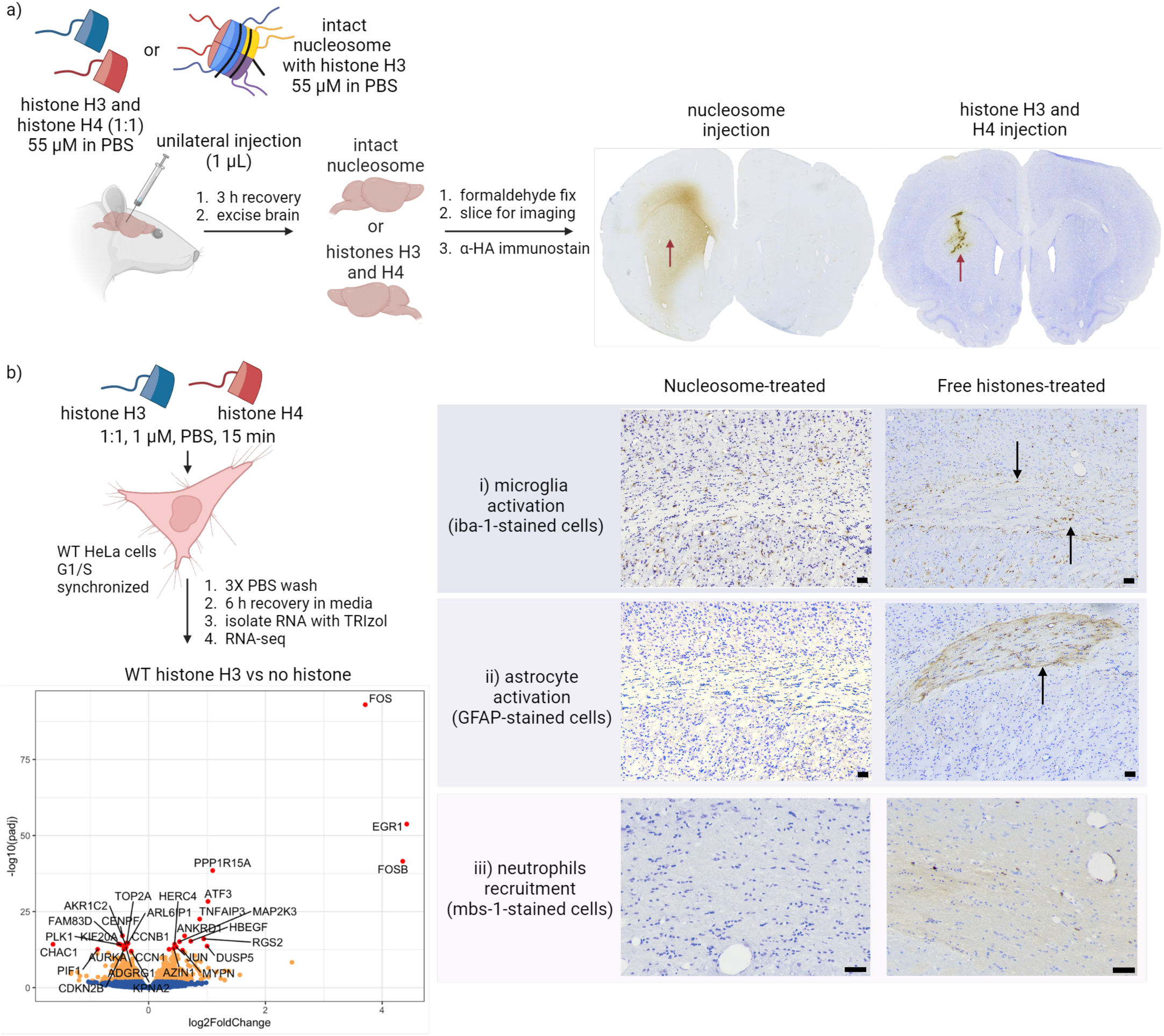
Synthetic histones and chromatin components in non-proteolytic tissue or cells. **a)** Immunohistochemistry (IHC) in rodent brain revealed striking differences in the diffusion and damage profiles between histone (right column) and nucleosome (left column). 3,3’- diaminobenzidine (DAB)-stained coronal brain sections immunolocalising the HA tag on a 10 μm- thick brain section from rats intracranially injected with nucleosome (left) and free histone (right) after 3 h. Red arrows indicate injection site. There is a loss of neurons adjacent to the histone injection site. Black arrows on histone-treated brain tissue revealed Iba-1-stained (brown) activated microglia surrounding a region of activated GFAP-stained astrocytes (brown region) and a few Mbs-1-stained neutrophils in vessels and within the brain parenchyma. IHC on nucleosome-treated brain tissue revealed the absence of microglial activation, astrocyte activation or the recruitment of neutrophils. Sections are counter-stained with cresyl violet. n = 3 per group. Scale bars = 50 μm. **b)** RNA-seq in synchronised HeLa cells exposed to extracellular histones shows transcripts profiles indicative of immediate early genes that respond to cellular stressors.

Next, and following these unexpected indications of putative histone-induced damage, all further investigations of the corresponding cellular damage response specific to extracellular histone H3 were restricted to cell culture. Initial studies tested the cytotoxicity of histone H3 on WT HeLa cells, revealing a concentration and time dependent cell viability response that became significant around 1 µM at 30 min (**Extended Data** Figure 8a). To assess the concomitant transcriptional response, cells were treated with WT histone H3 (again formulated with H4 1:1, 1 µM each in PBS) for 15 minutes under G1/S synchronized conditions^39^ (using double thymidine block^40^) and characterized via RNA-seq (**Figure 3b**). Notably, although there were no obvious signs of cellular toxicity, the transcriptional response to WT extracellular free histone H3 (*vs* no histone control, **Figure 3b**) clearly revealed representative so-called ‘immediate early genes’ that are commonly expressed in species as a first response to cellular stressors.^41^ These notably included FOS family proteins (regulators of cell proliferation, differentiation, and transformation through their roles in the activator protein 1 complex)^42,43^, and EGR1 (early growth response protein 1), a transcriptional regulator that modulates the immune response^44,45^. Moreover, a multitude of less, but nonetheless highly-significantly (-log10(padj) >10), altered transcripts were also identified that were consistent with emergent damage, including PPP1R15A and ATF3 that are known to promote growth arrest and DNA damage repair^46^ and/or are expressed upon physiological protein-induced stress^47^, respectively.

Together these data revealed seemingly contrasting physiological responses to extracellular histone H3 found in connective tissue where proteolysis is present (blood) and in tissue (brain) or cell culture where it is absent.

### FIRESCAPE suggests an intracellular fate for histone H3

These experiments also provided initial evidence of cellular uptake and internalization of extracellular histone H3 in both cell culture and tissue *in vivo*. First, there was apparent overlap of the synthetic histone H3 immunomarker (α-HA stain via a C-terminal HA epitope tag) and cresyl violet (used to visualise Nissl bodies and nuclei in neurons) in injected brain tissue (**Extended Data** Figure 8b). Second, the same immunomarker was also observed in intracellular protein extracts from the HeLa cells treated with extracellular histone H3 during RNA-seq experiments (**Extended Data** Figure 8c). Third, the observed transcriptional markers, such as ATF3, are typically upregulated only in response to intracellular protein.

Additionally, at short of time points (1 h) for administration of synthetic H3-[^18^F]Lys27 and H3-[^18^F]Lys4 histones into circulation we observed some specific accumulation of histone H3-[^18^F]Lys27 radioactivity, particularly in the lung and spleen, as organs with high exposure to blood flow (**Figure 4, right**). This contrasted with kidney and bladder accumulation of histone H3-[^18^F]Lys4 (indicative of rapid histone tail clipping and clearance, see above). Whilst the accumulation of positively-charged proteins in lung prior to transit to liver and spleen is known,^48,49^ cellular fractionation of both lung and spleen samples from our experiments strikingly revealed radioactive signals only in the nucleus of these associated lung- or spleen- derived cells (**Figure 4**). Moreover, autoradiographic-SDS-PAGE analysis of corresponding samples indicated bands at an expected molecular weight region for tail-cleaved histones. These bands were not evident in the samples from animals injected with histone H3 labelled at [^18^F]Lys4.

**Figure 4:**
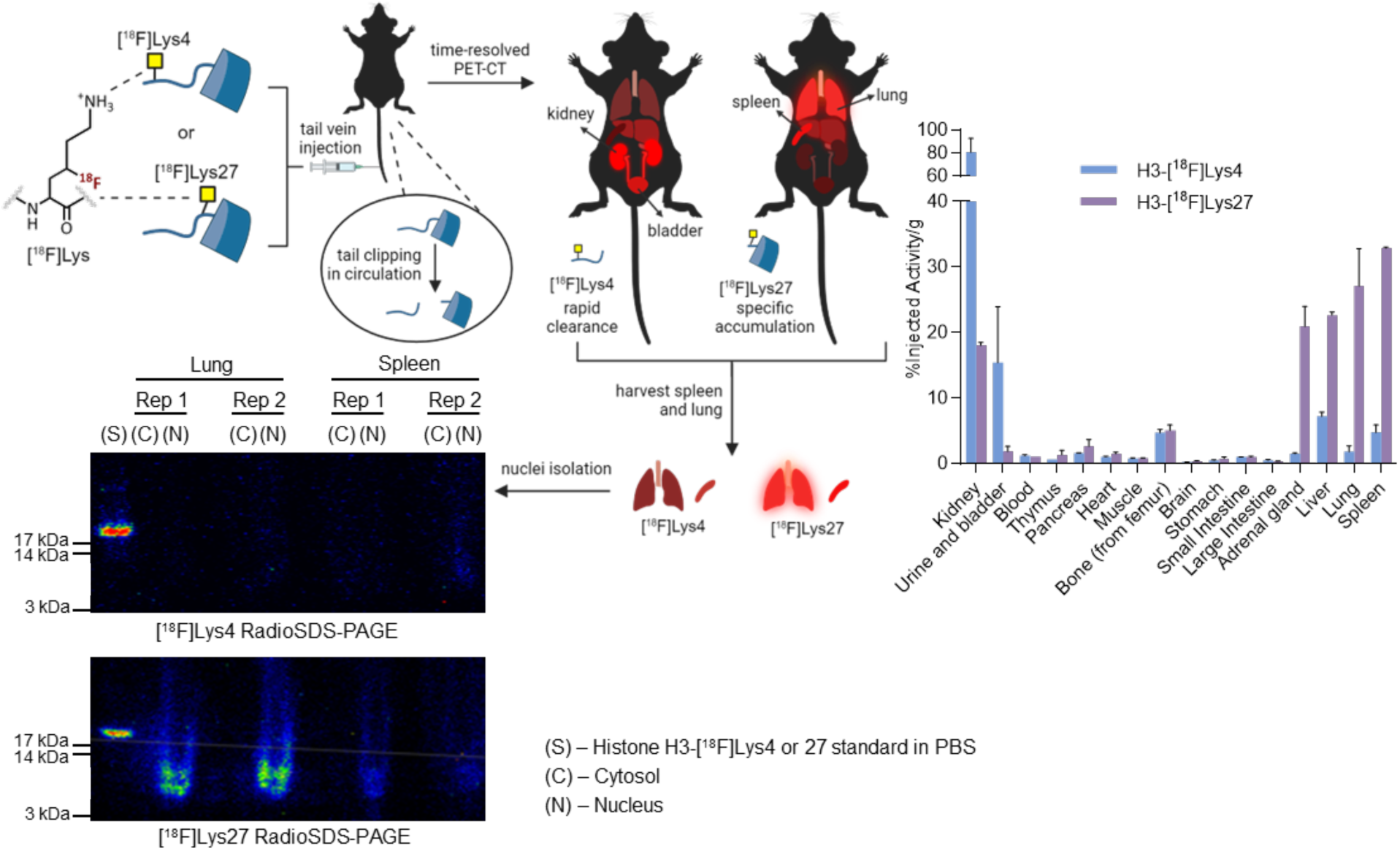
FIRESCAPE suggests an intracellular effect for histone H3. Radioactive signals corresponding to truncated histone H3 were detected in the nuclei (N) but not cytosol (C) of cells isolated from the lung and spleen (more weakly) 1 h after tail-vein administration of histone H3- [^18^F]Lys27 (n = 2). In contrast, cleavage of the N-terminus of H3-[^18^F]Lys4 containing [^18^F]Lys4 (n = 2) resulted in rapid clearance of radioactivity via the kidney and bladder (top right). Of note, the incubation time before sacrifice was reduced to 1 h. Radio-SDS-PAGE (bottom left) showed protein of lower molecular weight, consistent with truncated histone H3, reflecting the likely removal of the N- and / or C-terminal tails (as also observed by MS analysis, see **Extended Data** Figure 10).

These additional FIRESCAPE data not only indicated cellular uptake accumulating in the nuclei of specific organs reached through the circulatory system *in vivo* but also suggested that the core of histone H3, retained an ability to penetrate cells even after tail clipping.

### Extracellular histone H3 incorporates into chromatin in cell culture and in animals

We next characterized this apparent uptake of extracellular histones into cells via two further independent cellular extraction methods. First, we incubated WT HeLa cells with histone H3 (formulated as before with equimolar histone H4, PBS, 1 µM) for 15 min, followed by several PBS washes and then either 1 h or 24 h recovery in fresh media (**Figure 5a**) before collection and separation into cytosol, nuclear lysate, and chromatin fractions. Histone H3 was clearly detected in the chromatin fractions by blotting against its FLAG or HA C-terminal epitope tag, with lesser amounts only found at earlier time points (1 h) initially in the cytosol and nuclear lysate (**Figure 5a**). Histone H3 was solely located in chromatin after 24 h.

**Figure 5:**
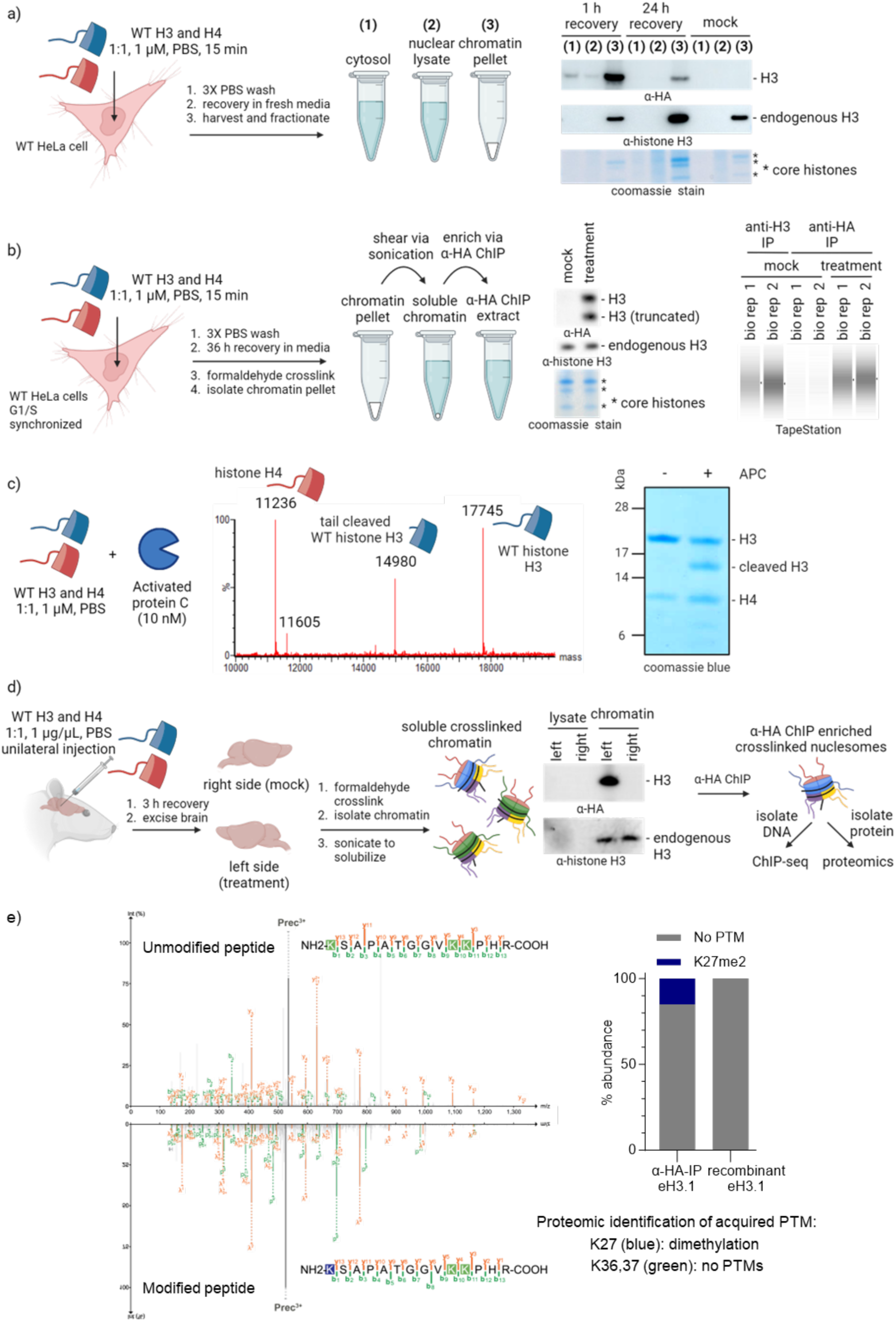
Extracellular histones can integrate into chromatin and become indistinguishable from native histones. a), **b)** Western blots of chromatin extracts from HeLa cells revealed localization of histone H3 in chromatin. Electrophoretic analysis (tapestation) on HA-ChIP extracts from histone-treated HeLa cells showed bound DNA enrichment compared to mock conditions. Size distribution of the sheared DNA matched shearing patterns of endogenous H3 IP. **c)** *In vitro* incubation of histone H3 (1:1 with histone H4) with Activated Protein C (APC) revealed a distinct mass corresponding to histone clipping predominantly between Arg26 and Lys27 (observed mass: 14980 Da, calculated mass: 14981 Da). **d)** In rodent brain tissue, exogenous histone H3 also incorporates into chromatin. Immunoprecipitation enabled ChIP-Seq and proteomic studies on the fate of these incorporated histones. **e)** Proteomic analyses confirmed the accumulation of abundant histone mark K27me2 on synthetic histone H3 at 3 h and 6 h timepoints following its introduction into rodent brain.

Next, we utilized a second, alternative chromatin extraction method to independently confirm the presence of histone H3 in chromatin, including a chromatin solubilization (via sonication) step to further investigate if incorporated extracellular histone H3 was bound to DNA, consistent with deposition into nucleosomes. G1/S synchronized WT HeLa cells were again incubated with synthetic, extracellular histone H3 (with equimolar H4, PBS, 1 µM each) for 15 min, washed with PBS and then allowed to recover for 36 h. Subsequent release into S phase then allowed cells to divide once and hence pass through S phase twice overall, thereby exposing cells to synthetic histone in a coordinated manner during processes (S phase DNA replication) where histones are most required.^50^ Through this orthogonal method, histone H3 was again identified (**Figure 5b**) in the soluble chromatin (Western blotting against HA epitope). We then performed an anti-HA immunoprecipitation (IP) from our soluble chromatin samples (ChIP), followed by decrosslinking. Notably, automated electrophoretic analysis (tapestation, **Figure 5b**) showed that bound DNA was enriched ∼15-fold more than under mock conditions and displayed a size distribution matching that of the shearing patterns of endogenous histone H3 IP. This indicated that incorporated synthetic histone H3 was associated with DNA in this soluble chromatin fraction (**Figure 5b**). Furthermore, in parallel experiments, this prevalence of histone localisation in chromatin was also observed after 3 h without requiring G1/S synchronization (**Extended Data** Figure 8d).

Short-timescale FIRESCAPE had suggested intracellular uptake even of clipped histone H3 (see above). Of the known proteases in connective tissue that are absent in brain, activated protein C (APC)^15^ is one of the most prevalent. Although we cannot discount the possibility that the proteolysis observed by FIRESCAPE in circulation may be caused by other proteases, when we incubated histone H3 with APC, both electrophoretic (truncated band ∼14 kDa, **Figure 5c, right**) and mass spectrometric analyses (**Figure 5c, middle**) revealed a precise primary proteolytic product. Notably, these corresponded to a clean cleavage event between residues Arg26–Lys27^51^ (calculated mass: 14981 Da, observed mass: 14980 Da), exactly mirroring the apparent histone tail targeting proteolysis revealed both *in vitro* and *in vivo* by our FIRESCAPE methodology. Moreover, the resulting histone H3, post-APC cleavage at Arg26/Lys27 (1 μM formulated with H4, 10 nM APC, 2 h, 37°C), retained the ability to localise in the soluble chromatin fraction of HeLa cells (**Extended Data** Figure 10a).

Finally, we also probed incorporation of synthetic histone H3 into chromatin in tissue. Utilizing brain tissue from the earliest timepoints (3 h post-injection) of our prior experiments (see above) allowed access to samples that exhibited the least amount of cell death. Chromatin extraction via essentially identical methods revealed the incorporated presence of injected exogenous histone H3 (**Figure 5d**). Moreover, analysis of DNA enriched from ChIP (α-HA) revealed expected fragment sizes with excellent enrichment over the mock condition (over 100- fold, tapestation, **Extended Data** Figure 9a). Sequencing of this DNA (obtained via α-HA ChIP-seq) gave results that revealed strikingly broad histone incorporation, highly similar to both control endogenous α-histone H3 ChIP-seq, and also to input (**Extended Data** Figure 9a). Moreover, α- histone H3 ChIP-seq, thereby sampling all endogenous H3, indicated no differences between treated and untreated conditions. Together these data, remarkably, not only further confirmed that extracellular synthetic histone H3 could penetrate cells *in vivo* but also revealed that upon uptake it is incorporated into chromatin in a global manner throughout the genome, with its chromatin deposition indistinguishable from that of endogenous histone H3.

### Extracellular histone H3 is a functional epigenetic substrate after penetrating cells and depositing into chromatin

These results indicated that extracellular histone H3 appears to mimic endogenous histone H3 at least in *location* once it has penetrated cells by depositing throughout chromatin both *in vitro* and *in vivo*. To also test the functional competence of this exogenous H3, once incorporated, we attempted to directly assess its putative conversion via *in vivo* epigenetic ‘writing’. To our knowledge, proteomic survey of such *in vivo*-acquired PTMs has not yet proven possible.

First, proteomic analyses of α-HA ChIP protein elutions were used to both independently probe exogenous histone H3 incorporation and to also assess the associated dynamic range of mass spectrometric (MS) sensitivity (**Extended Data** Figure 9b). These analyses further confirmed enrichment of the HA tag only into chromatin from treated tissue (left brain, 3 h and 6 h post injection, **Extended Data** Figure 9c), thereby validating both the α-HA ChIP and the presence of histone H3 in chromatin, as well as indicating the feasibility of MS analyses of such samples.

Next, electrophoretic analysis (visualized via silver stain, **Extended Data** Figure 9d) identified exogenous histone H3 in these samples, distinguishable from endogenous H3 (not observed) via its larger molecular weight (due to the added epitope tags, **Extended Data** Figure 9c**, 9d**).

Finally, gel excision followed by in-gel treatment with propionic anhydride allowed isolation and subsequent proteomic analyses. Strikingly, these unambiguously identified multiple histone PTMs: histone H3 K27me2, K18ac, and K23ac (**Figure 5e** and **Extended Data** Figure 9e). Notably, K27me2 is one of the most common PTMs^52^ distributed throughout large chromatin domains – this was observed in ∼15% of the corresponding total peptide in two independent biological samples recovered at two different timepoints (3 h and 6 h post-histone H3 injection, **Figure 5e**). Interestingly, whilst at 3 h K27me2 was the only PTM observed, further lysine modifications accumulated over the subsequent 3 h, including acetylation on K18 and K23 (**Extended Data** Figure 9e). Although, precise quantification of these additional PTMs was prevented by inconsistent frequencies across biological replicates at the limits (low ng) of MS detection, in all cases these detected, emergent, *in vivo* modifications of synthetic, exogenously-introduced histone H3 were absent in negative controls (anti-histone H3 IP on histone-treated sample; anti-HA IP on non-histone-treated sample; and unmodified recombinant histone H3 before administration). This first detection of *in vivo* epigenetic modifications of a synthetic protein employed *in vivo* – leading to not only acquired highly-abundant native PTM K27me2 but also others – suggested that this exogenous histone H3 is treated essentially indistinguishably from endogenous histone, once inside cells.

## Discussion

Through precise editing of ^18^F-containing amino acid residues, here [^18^F]hAla or [^18^F]Lys, into chosen protein domains FIRESCAPE exploits the ‘near-zero-size, zero-background’ ^18^F radioisotope as a tracer of extremely low doses, while still enabling precise and time-resolved PET tracking. This simultaneously reveals striking differences in proteolytic susceptibility; clearance kinetics; as well as tissue, cellular and even cell-fraction associated uptake – all enabled by sensitivities that are beyond current methods (see **Supplementary Note 3**). With FIRESCAPE, we can now precisely scan the entire length of a protein sequence for proteolytic susceptibility and clearance *in vivo*, even including the specific residues that are themselves proteolytically engaged (P_1_, P_1_′ residues) via the mimicry of corresponding ‘radioequivalent’ proteins (**Extended Data** Figure 5c). Our data therefore now shows unequivocally that label site not only matters but can provide a previously unrealised insight. For example, previous assessments of nucleosome clearance has utilised non-selectively labelled “histone components^16^”, almost certainly on the histone tails that we now show have privileged proteolysis and clearance pathways that are independent of the rest of the core. We therefore caution that such prior work, while honestly using the tools available at the time, must now be critically reassessed in the context of precise editing that we reveal here.

Notably, mid-twentieth century observations^53,54^ using isolated calf thymus histones^55,56^ led to ongoing suggestions that plasmid DNA^57–59^, RNA^59^ and proteins^56,60^ could be delivered into a variety of cell lines when in complex with histones, in a method coined “histonefection”^53^. Although the physiological relevance was not explored nor sought in this suggested technological application, all core histones (H2A, H2B, H3 and H4) as well as linker histone H1 seemingly exhibited some ability to penetrate cells in multiple mammalian cell lines, attributed in some part to charge,^53,60,61^ yet the associated utility and mechanism of membrane penetration has remained controversial. Several charged carrier•DNA complexes enter cells via endocytosis,^62,63^ and using histones as ‘carriers’ for DNA and proteins may function similarly;^64^ reduced uptake under energy depletion is consistent with such a mechanism.^60^

Overall, therefore, the fates, effects and utilities of extracellular chromatin components have remained poorly resolved. There is a clear disconnect, for example, in the suggested toxicity of extracellular histones^15^ yet also their use as “non-toxic” delivery agents for gene therapy.^53,54,61^ Our studies now bring some clarity to these discrepancies, by characterizing the circulatory distribution, clearance, damage response, cellular uptake, genomic localization and PTM accumulation of extracellular histone H3 in cells and animals. Cumulatively, our data suggest that extracellular histone H3 is rapidly proteolyzed in circulation but demonstrates a robust ability to penetrate the plasma membrane and enter cells in mammalian cell-cultures, rodent brain tissues, and seemingly other organs, via the circulatory system. Once in cells, extracellular histone H3 is apparently treated indistinguishably from endogenous histone H3 in deposition and as a functional epigenetic substrate.

This discovered molecular interloping may provide a possible explanation for the robust damage response of cells to histone uptake that we also observe. Cells apparently therefore cannot differentiate between endogenous histones and extracellular histones that have entered the cell^65^, leading to deposition of extracellular histones throughout the genome. Extracellular histones may therefore present a danger, in that they may carry PTMs (‘marks’) arising from and / or representative of damaged or diseased cells. These histone marks may therefore have epigenetic consequences once incorporated stochastically into the chromatin of healthy cells. Furthermore, while truncated forms of histone H3 retaining the histone fold domain can still form nucleosomes, the *N*-terminal region has been shown to regulate higher-order chromatin structure and gene regulation *per se*.^66^ Whilst we observe that histone marks are likely removed via the clipping of their tails when in circulation, as we also show here there could be specific tissue contexts and/or pathologies (such as in stroke^6,8,36^ or tumours) where released extracellular histones are protected from proteases, favouring intact uptake with marks into proximal cells. Moreover, chromatin incorporation of extracellular histones (recycling) may ensure their long-term retention, given that some endogenous histones are known to exhibit exceptionally long physiological half-lives^67^. We also cannot discount the possibility that mutations at proteolytic sites, some of which are known to be relevant to disease (e.g. Lys27Met ^68–70^), may also alter susceptibility to such proteolysis (**Extended Data** Figure 10b) and so also modulate subsequent survival and uptake, even in proteolytic connective tissue. It may also be that proteolytic processing of histone tails to erase marks in serum is an evolutionarily consequence of the ability of extracellular histone to incorporate into chromatin with potential epigenetic implications^71–73^ or, more prosaically, simply because the tail is more disordered and so accessible to enzymatic activity.

Finally, we should note that whilst we focused here on use of FIRESCAPE to resolve some mysteries of potentially toxic (histone) proteins *in vivo* by exploiting its sensitivity in ultra-low doses, it has clear broader potential. These include the *in vivo* monitoring of specific proteins targeted for degradation by emergent technologies,^74^ ^75,76^ as well as the application to scanning beyond the initial ‘radioequivalent’ residues ([^18^F]hAla or [^18^F]Lys) that we suggest here. Work to this effect is ongoing in our laboratories.

## Author Contributions

B.J., A.W.J.P., V.G., D.C.A and B.G.D. conceived the project and designed experiments.

B.J. designed and generated histone variants.

B.J. and A.W.J.P. performed chromatin extraction on HeLa cells and mice/rat brain.

A.W.J.P. designed and performed all ^18^F-protein experiments.

A.W.J.P. and A.M.G. developed ^18^F-Lysine radical reagent.

A.W.J.P., J.F. and J.B.I.S optimised radiosynthesis and A.W.J.P., T.A., N.A.R. and A.J.A. automated synthesis of ^18^F-Lysine radical reagent on Trasis AllinOne.

A.G.Y, K.T., S.A., M.W.G.M, L.K., J.S.H., R.A., A.M.D, K.A.V. and D.C.A. performed *in vivo*

mice and associated biodistribution experiments.

A.W.J.P. investigated histone cleavage.

A.G.Y, I.K.D. and D.C.A performed IHC on brain slices.

B.J. and M.P. performed transcriptomics and data was analysed by A.P.C.

B.J. and A.W.J.P. performed ChIP-seq and data was analysed by C.Y.W.

Y.D. carried out MS and data analysis.

W.L. performed cell viability assay.

A.B. generated nucleosomes.

B.J., A.W.J.P. and B.G.D. wrote the paper and all authors commented.

## Acknowledgements

We thank Johan Rajander (Åbo Akademi) for production of fluorine-18 and Mira Eisala, Aake Honkaniemi and Marko Vehmanen for technical assistance during *in vivo* mouse experiments and PET imaging at Turku PET Centre, Finland. We thank the following for funding: University of Brunei Darussalam Chancellor Scholarship (to A.W.J.P), UKRI-BBSRC (BB/V010999/1 to B.G.D., V.G. and K.A.V.). The Chemistry theme at the Rosalind Franklin Institute has been supported by grants from UKRI-EPSRC (EP/V011359/1, EP/T012021/1, EP/X527245/1).

## Potential Conflicts of Interest

A.P.C. is listed as an inventor on several patents filed by Oxford University Innovations concerning sequencing technologies. A.W.J.P., B.J., V.G. and B.G.D. are listed on patents filed by Oxford University Innovations concerning protein editing. A.P.C. is also a founder and employee of Caeruleus Genomics.

## Data Availability

Sequencing data have been deposited in the GEO under accession number GSE166493 (for RNA-seq analysis, sample names eH3WT and NoeH3) and GSE276733 (for ChIP-seq analysis).

## SUPPLEMENTARY NOTES

Supplementary Note 1

For our histone H3 construct, we chose to use an epitope tagged construct of the canonical core human histone H3, often called histone eH3.1 in literature (“e” denotes a C-terminal FLAG-HA dual-epitope tag) which has been successfully used in many *in cellulo* experiments with established protocols for immunoprecipitation;^82,83^ its epitope tags do not interfere with native function^84^. These epitope tags further differentiated our exogenous histones from any endogenous histone H3 present in the experiments and allowed for later extraction and analysis (see Figure 3). Both native Cys residues, C96 and C110, were mutated to Ala, as is commonly done in biophysical studies, to enable selective Cys based reactions at sites of interest.

Supplementary Note 2

Histone H3 exhibited poor solubility in PBS alone, displaying visible aggregation within seconds to minutes at the concentrations needed for the studies throughout this work. Initial screens found that by first incubating histone H3 with equimolar amounts of histone H4 in water, the mixture could then be exchanged to PBS while retaining full solubility of both proteins. We note that histones H3 and H4 are natural partners in cells, either during their occupancy in nucleosomes (also with H2A and H2B), but also while being chaperoned into the nucleus as a H3-H4 tetramer^3^. Therefore, we believe their stability when paired is expected and appropriate and was required to carry out these studies.

Supplementary Note 3

Historically, proteolytic clipping of nuclear histone H3 was identified through immuno-based assays^85^ or protein sequencing techniques (e.g. Edman degradation)^86^, applicable only to high concentrations found in nuclear extracts. Mass spectrometric mapping of cleavage sites^71,87^ curtailed when investigating low abundance extracellular histones in the blood (ranging up to only 0.06 ng/mL under healthy conditions^8^) due to the wide dynamic range of the blood proteome (e.g. serum albumin is present in the 35 – 50 mg/mL range).^88^ Analysing products of blood proteolysis is further complicated by ongoing clipping during sample extraction preparation. Furthermore, the *N*-termini of histones are rich in lysine and arginine residues and thus, produce peptides^89^ that are incompatible with classical MS detection parameters. In rare examples where histone H3 was observed in human body fluids of older or critically ill patients, e.g. urine^90^ and serum^91^, it has only been identified from core peptides.

Supplementary Note 4

Tail-labelled H3-[^18^F]hAla4 exhibited a biphasic clearance pattern (n = 1), characterised by a fast and slow half-life of 1.7 min (59%; 95% CI, 0.7 - 3.0 min) and 13.7 min (41%; 95% CI, 8.9 - 25.3 min), respectively, resulting in a weighted blood half-life of 6.6 min. Notably, unlike other [^18^F]hAla proteins^27^, radioactivity was retained in the heart tissue, contributing to the slower clearance phase. On the other hand, core-labelled H3-[^18^F]hAla56 showed significantly prolonged retention in circulating blood (n = 1), displaying a half-life that resembles a single- phase clearance rate of 15.2 min (95%, 7.6 – 34.0 min).

Supplementary Note 5

We note the ongoing emergence of a consensus on nomenclature.^92,93^ Here, radiochemical yields (RCYs) were determined by radio-HPLC analysis of the crude reaction mixture. Activity yields (AYs) of isolated and purified ^18^F-compounds are reported as % with respect to the activity of the starting ^18^F-fluorinating reagent, unless stated otherwise. RCY and AY values were not corrected for decay. Chemical purity of isolated ^18^F-compounds was based on radio-HPLC traces and UV detection.

**Extended Data Figure 1:**
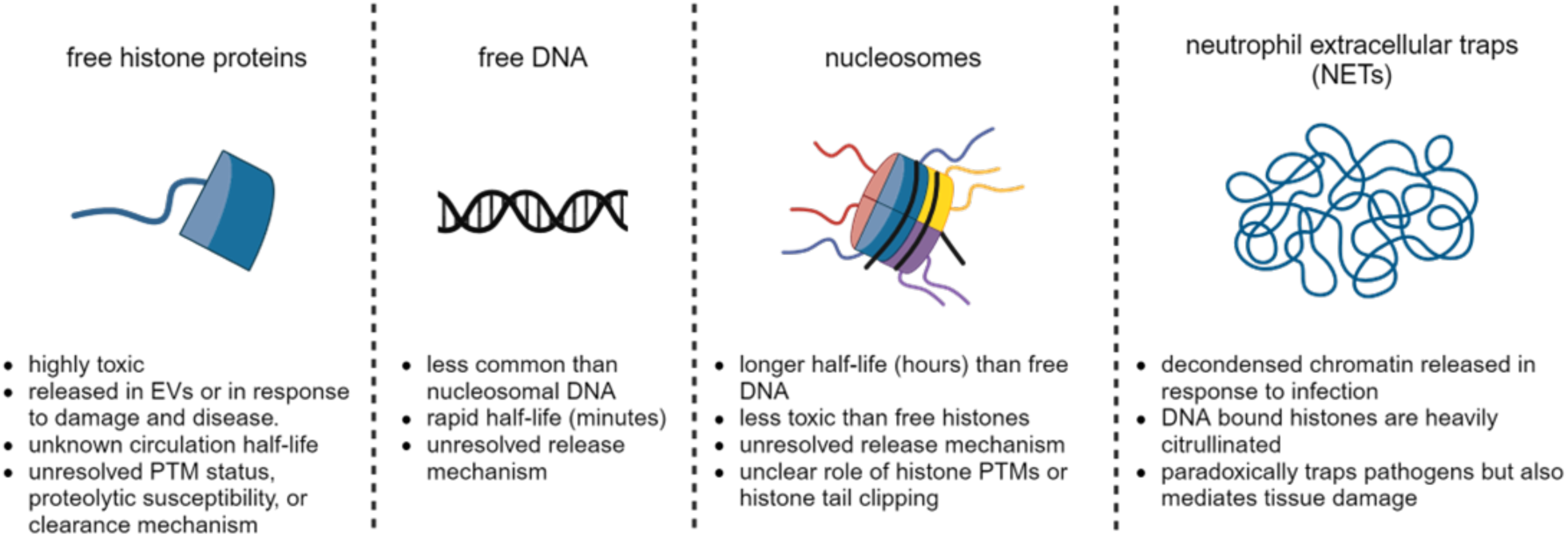
Chromatin components released into circulation, and associated roles and effects.

**Extended Data Figure 2:**
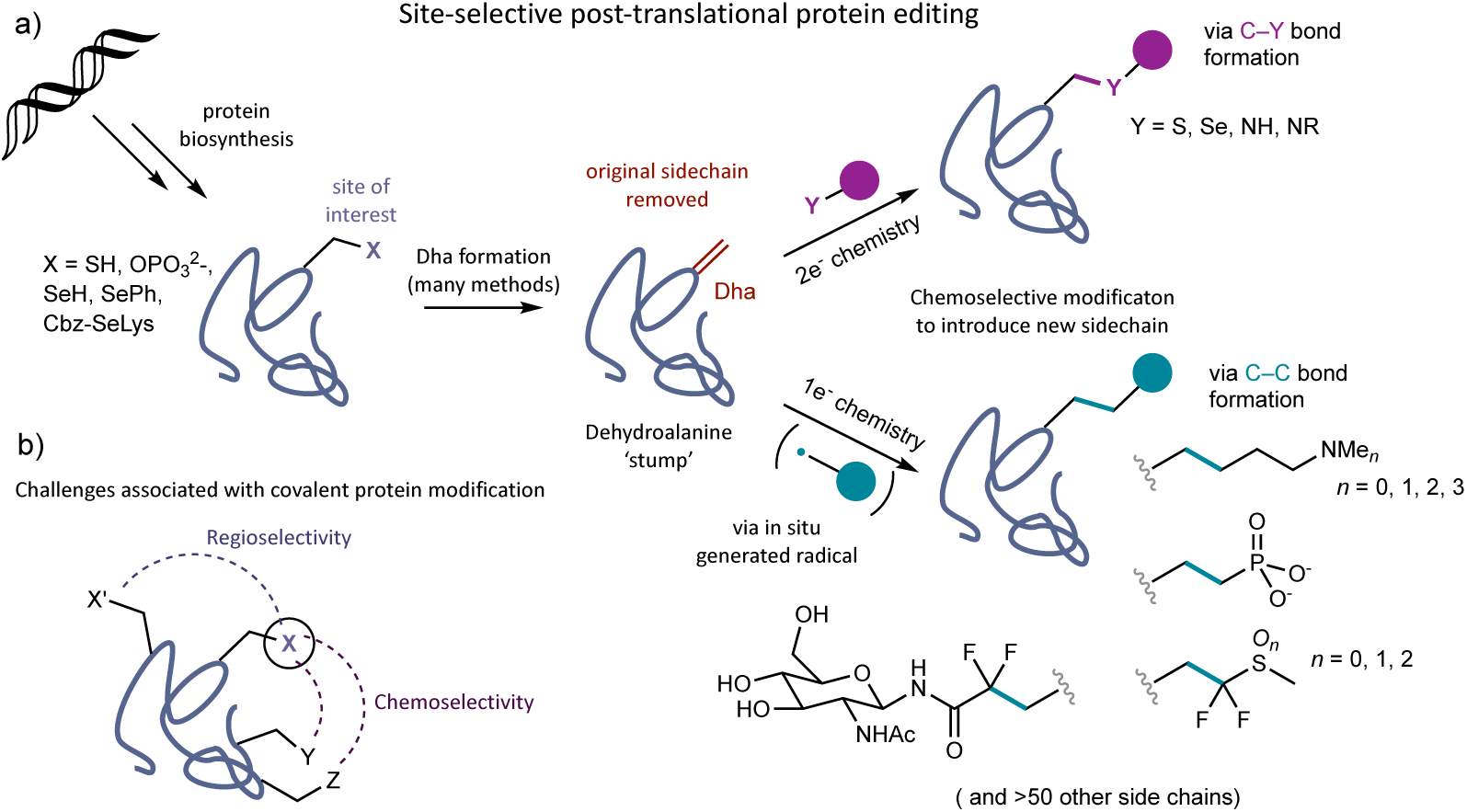
Recent work has made possible the chemical, post-translational editing of specific sidechains in purified full-length proteins of interest, including histones. Following combined genetic (mutate residue at site of interest to precursor such as Cys) and chemical removal of the side-chain (convert Cys to unnatural ‘stump’-residue amino acid dehydroalanine (Dha)), a new sidechain can be grafted onto the Dha sidechain ‘stump’ to reprogramme the primary sequence of the protein at that site, as well as allowing in principle any new sidechain of interest to be installed. Recent work has allowed for these new edited sidechains to consist of natural or PTM containing sidechains, or even unnatural sidechains containing unique functionalities rare in biology, such as the incorporation of the element fluorine. We reasoned that an extension of this methodology would allow for both ^18^F-containing bioisosteres and even ‘radioequivalent’ sidechains to be installed at any position within a protein of interest. The latter consists of a native residue bearing a ‘near-zero-size’ C–H-to C–^18^F bond substitution within the sidechain.

**Extended Data Figure 3:**
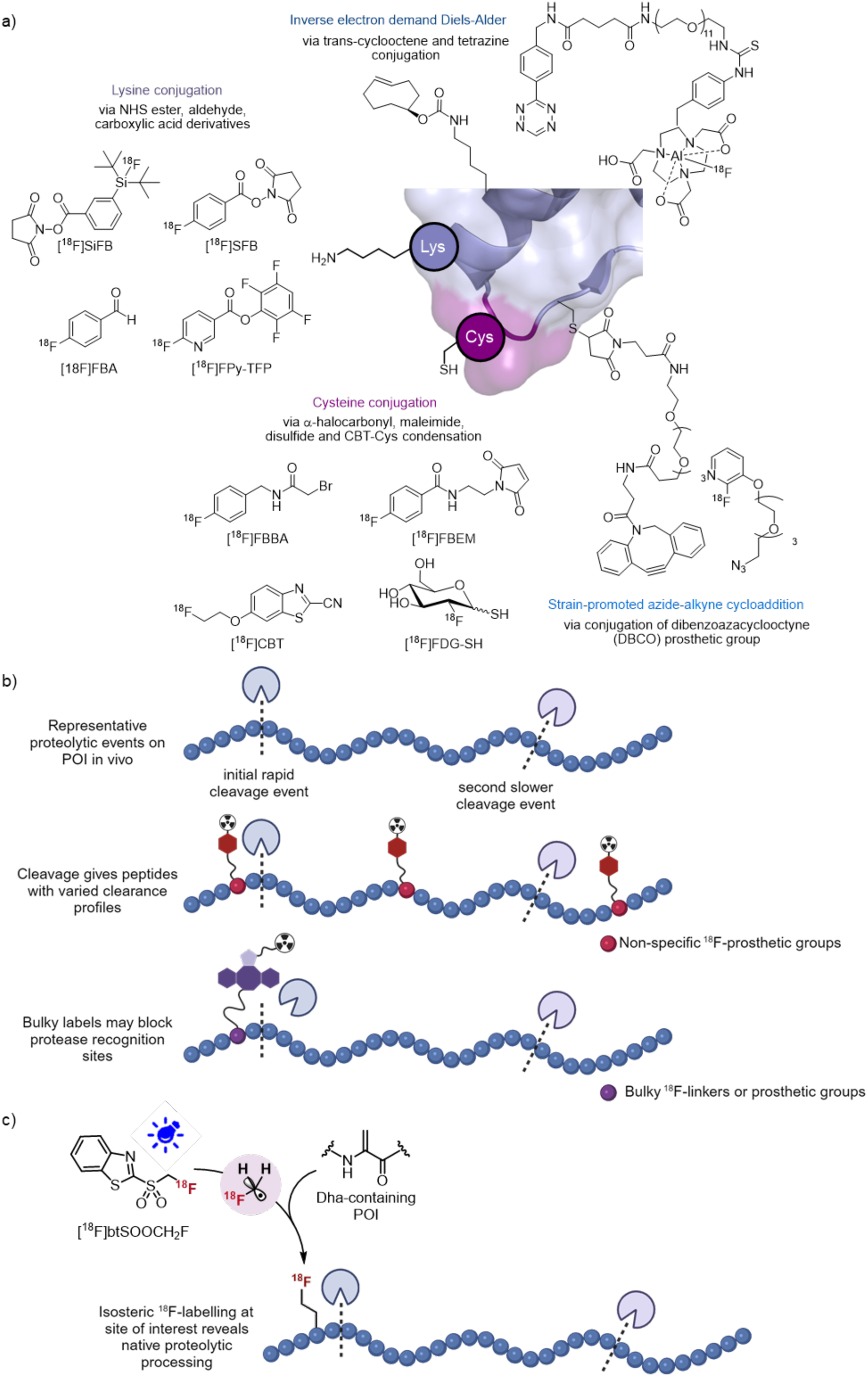
Conventional protein radiolabelling strategies to profile *in vivo* proteolysis is unsuitable. a) Radiolabelling often occurs non-site-specifically on lysine or cysteine side chains. Rapid bioorthogonal reactions (e.g. strain-promoted azide-alkyne cycloaddition^77,78^, inverse electron demand Diels-Alder^79^) have also been employed to install ^18^F-labelled, usually bulky, prosthetic groups. b) Random conjugation or the use of bulky linkers and/or prosthetic groups may give an incomplete or even inaccurate analysis of protease activity in the blood. c) To better characterise proteolytic processing *in vivo* and in real-time FIRESCAPE utilizes precise protein editing (see Figure 2 for the hierarchical FIRESCAPE strategy). Shown here, the first ‘broad’ scanning round of FIRESCAPE exploits the radical precursor^27^ [^18^F]btSOOCH_2_F for site-specific protein radiolabelling through the efficient generation of isosteric sidechains using ^18^F-containing carbon-centred radicals.

**Extended Data Figure 4:**
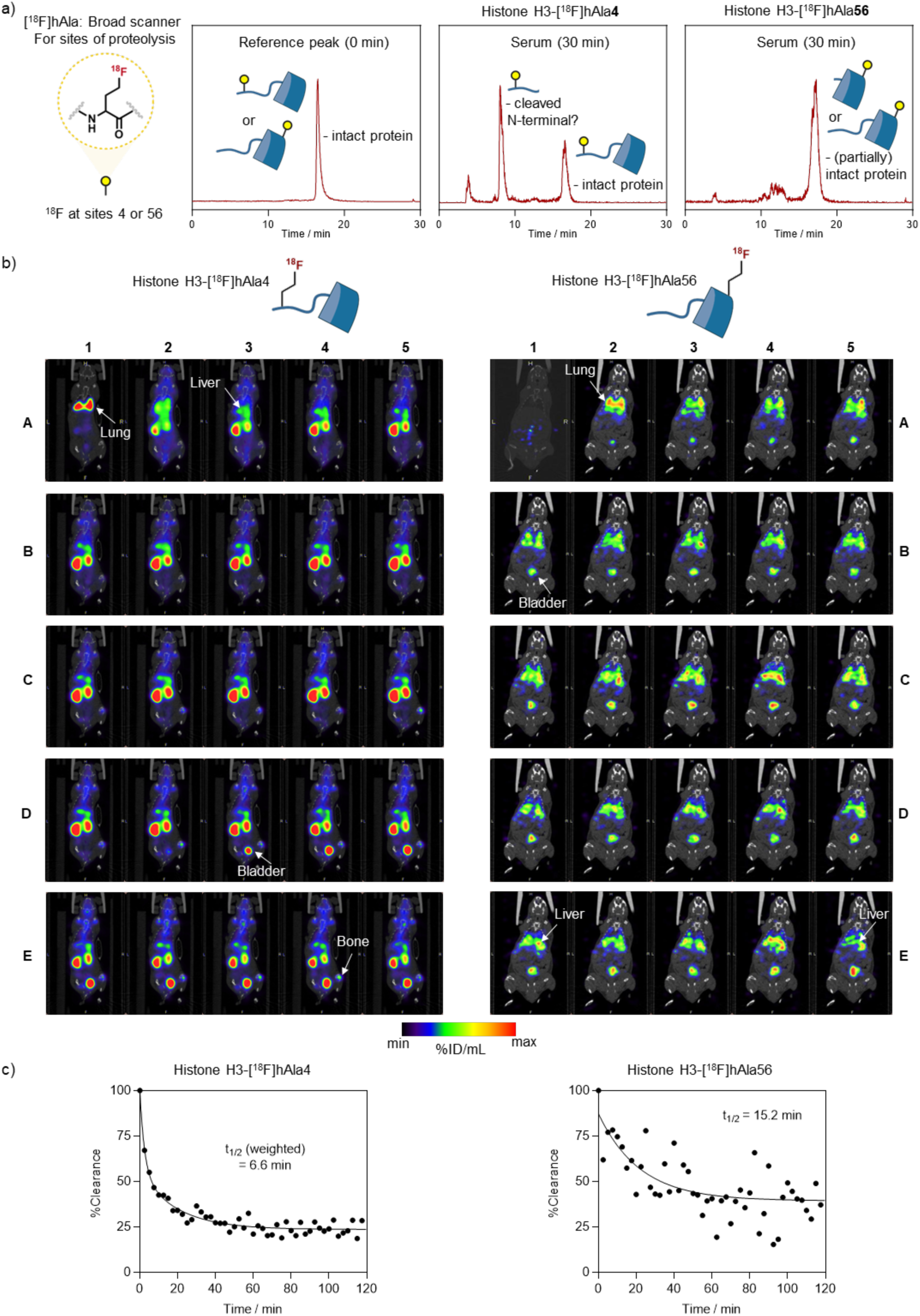
Mapping proteolysis of Histone H3**-**[^18^F]hAla4 vs [^18^F]hAla56 in a) serum following *ex vivo* incubation and analysis by radio-HPLC and b), c) in the circulatory system of living mice using dynamic PET, with the associated blood clearance weighted half-life shown in the bottom (left-[^18^F]hAla4, right-[^18^F]hAla56; (n = 1 each)). Images were taken every 5 min and arranged from left to right i.e., A1 is the earliest time point immediately after injection, A2 is taken 5 min post-injection and E5 is the last time point taken before organ harvest for *ex vivo* radioactivity counting (2 h post-injection).

**Extended Data Figure 5:**
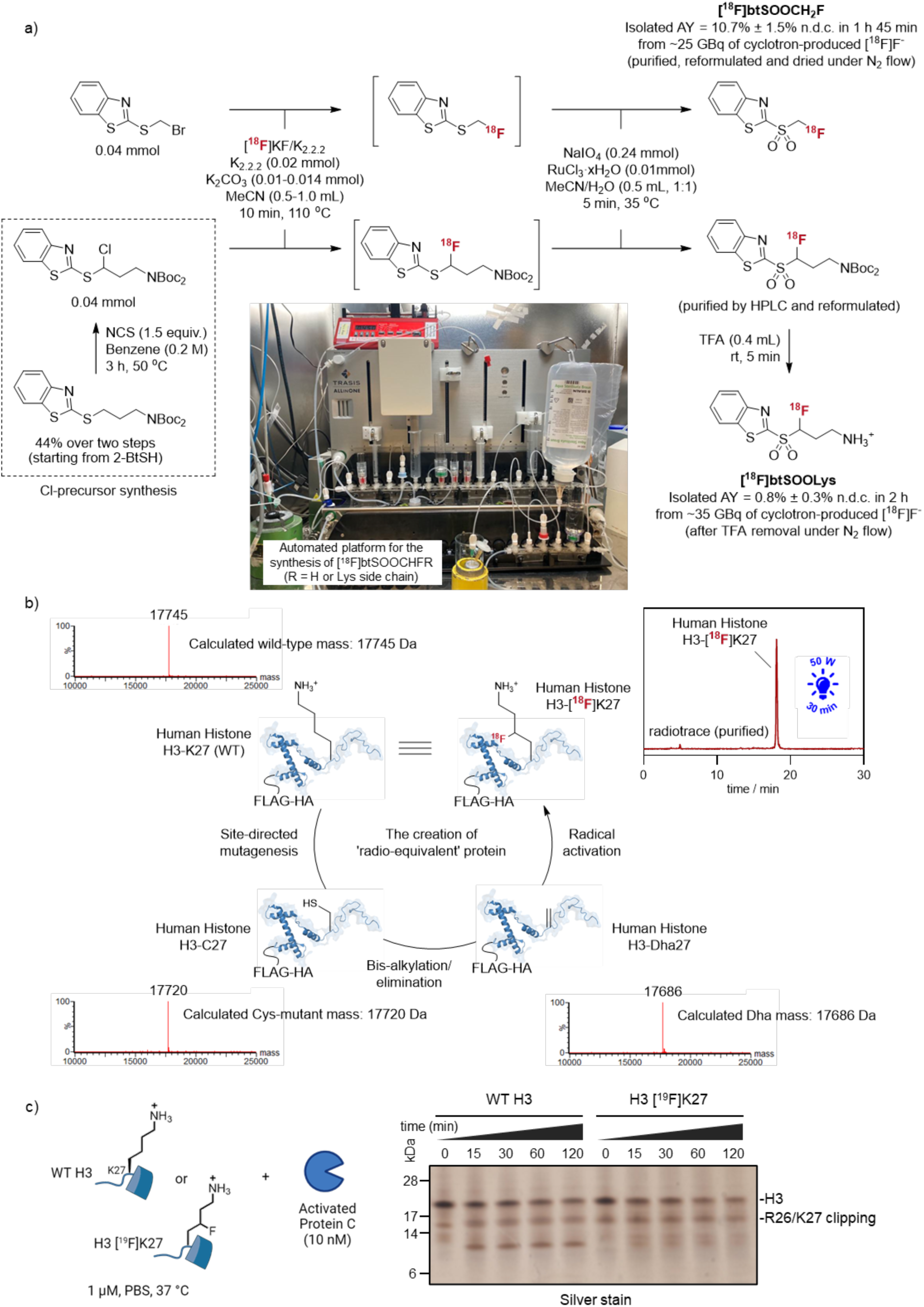
Creation of a ‘radio-equivalent’ protein. a) TrasisAllinOne platform for the automated two-step, one-pot synthesis of [^18^F]btSOOCHFR (where R is H or a Lys side chain). b) Systematic editing enables the radiolabelling of proteins at an atomic level. A residue of interest can be mutated to cysteine by site-directed mutagenesis. Following subsequent chemical generation to Dha, visible light mediated SET and [^18^F]btSOOLys as the radical precursor enabled recapitulation of the native side chain residue containing theoretically, the smallest perturbation to incorporate a radioactive isotope, that is, an isosteric H → [^18^F]F substitution. Intermediate species can be conveniently tracked by intact LC-MS analysis. c) Time course *in vitro* incubation with Activated Protein C on histone H3 fluorinated on Lys27 (H3- [^19^F]K27) showed retained cleavage between Arg26 and Lys27.

**Extended Data Figure 6:**
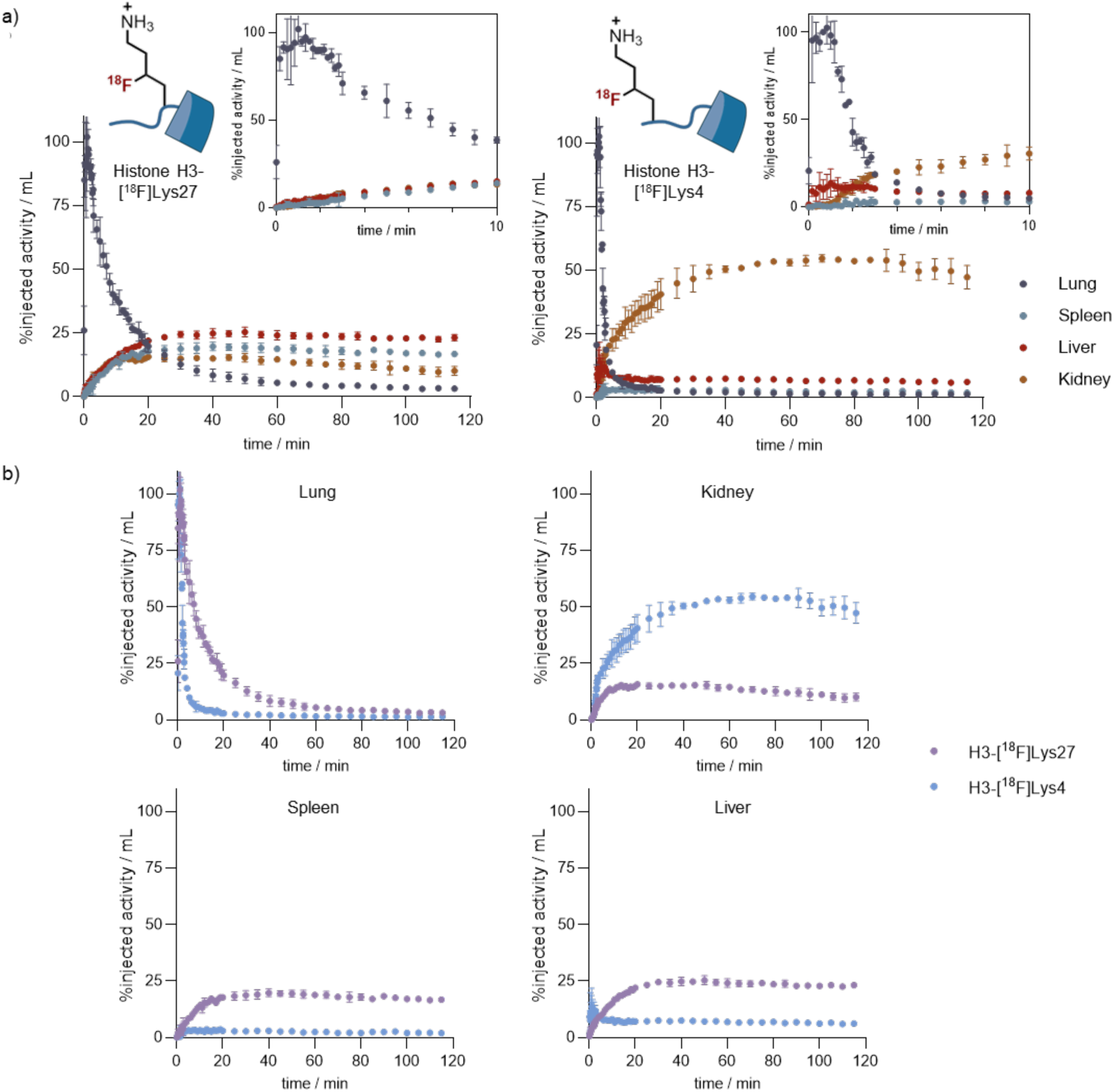
a) Time-activity curves across various tissues, including those involved in the clearance pathway, derived from VOI analysis of dynamic PET images following tail vein administration of histone H3-[^18^F]Lys27 (left) and H3-[^18^F]Lys4 (right). b) Comparison of radioactivity accumulation or clearance in proteolytic tissue between H3-[^18^F]Lys27 (purple) and H3-[^18^F]Lys4 (blue).

**Extended Data Figure 7:**
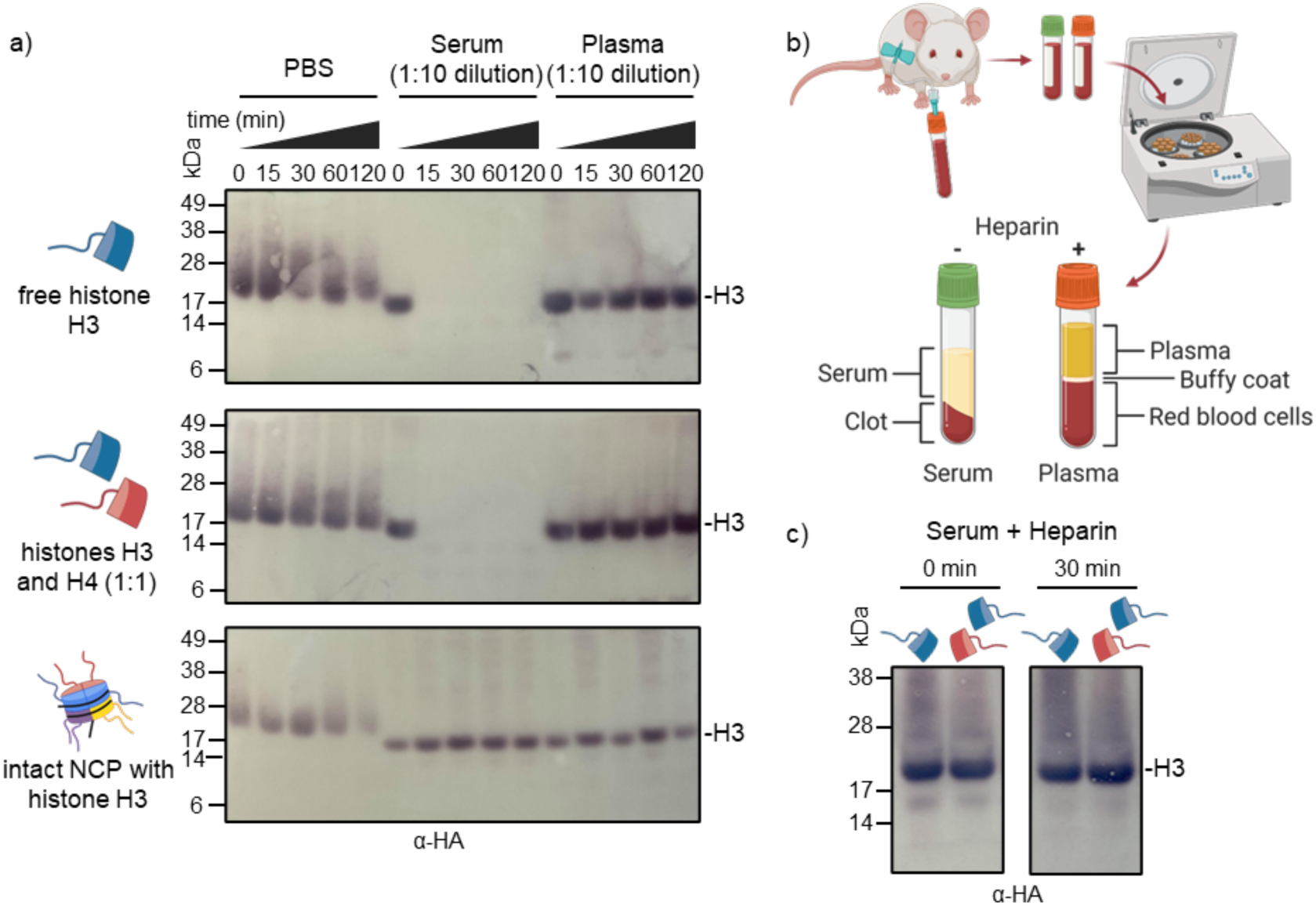
Susceptibility of free histones and intact nucleosomes to proteolysis in serum. a) *In vitro* incubation with blood serum followed by electrophoretic Western time-course analysis showed that both free histone H3 and an equimolar mix of histones H3 and H4 underwent rapid proteolysis while nucleosome core particles, where the backbone amide linkage between Arg26/Lys27 is seemingly well protected^80^, retention of the epitope-tagged histone band indicated that NCPs remained intact even in serum. b) Schematic illustrating the extraction of plasma and serum from mice. Heparin was used as the anticoagulant in plasma. c) The addition of heparin, a known histone interactor in circulation^81^, to serum negates proteolysis of free histones, as revealed by western blotting against the HA epitope tag on histone H3. Note: Mixing heparin directly with free histones resulted in rapid precipitation, likely due to the negatively charged heparin binding the positively charged histone tails, blocking proteolysis and promoting precipitation.

**Extended Data Figure 8:**
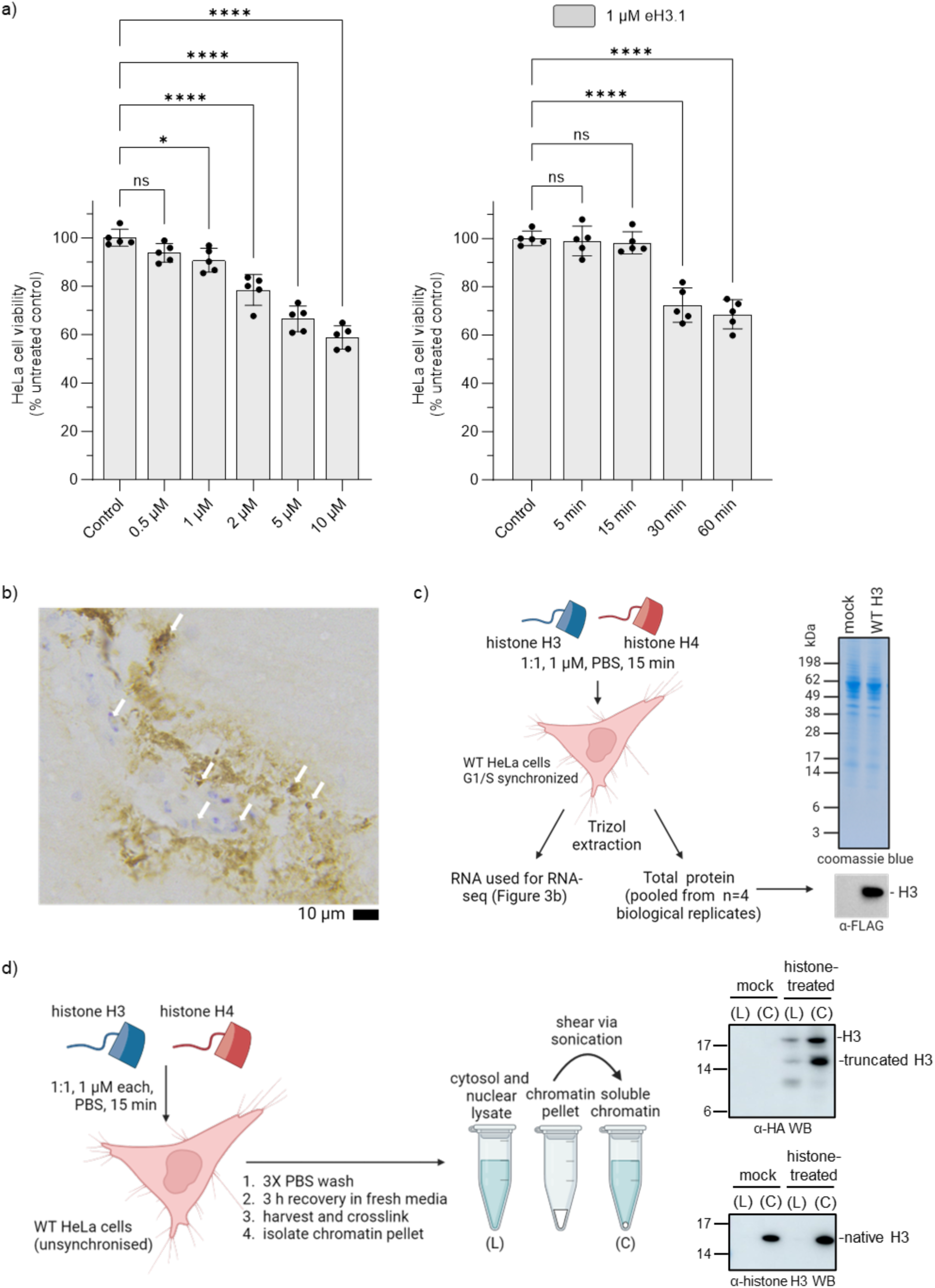
Evidence for exogenous histone penetration into cells and localisation in chromatin. a) Cell viability assay to determine histone concentration (left) and exposure period (right) suitable for HeLa cell studies. Statistical analysis for cell viability assay used one-way ANOVA. Error bars represent standard deviation from 5 independent biological replicates. ****p<0.0001, ***p<0.001, **p<0.01, *p<0.05, ns – not significant. b) Anti-HA immunostaining on fixed brain slice post-histone H3 administration (1:1 with Histone H4, 55 μM in PBS, 1 μL) suggested co-localisation of exogenous histone H3 with a nuclear (cresyl violet) stain. c) Western analysis (anti-FLAG) on TRIzol protein extracts of histone-treated HeLa cells showed the presence of exogenous histone H3. d) Localisation of exogenous histone H3 into chromatin was also observed for unsynchronised HeLa cells.

**Extended Data Figure 9:**
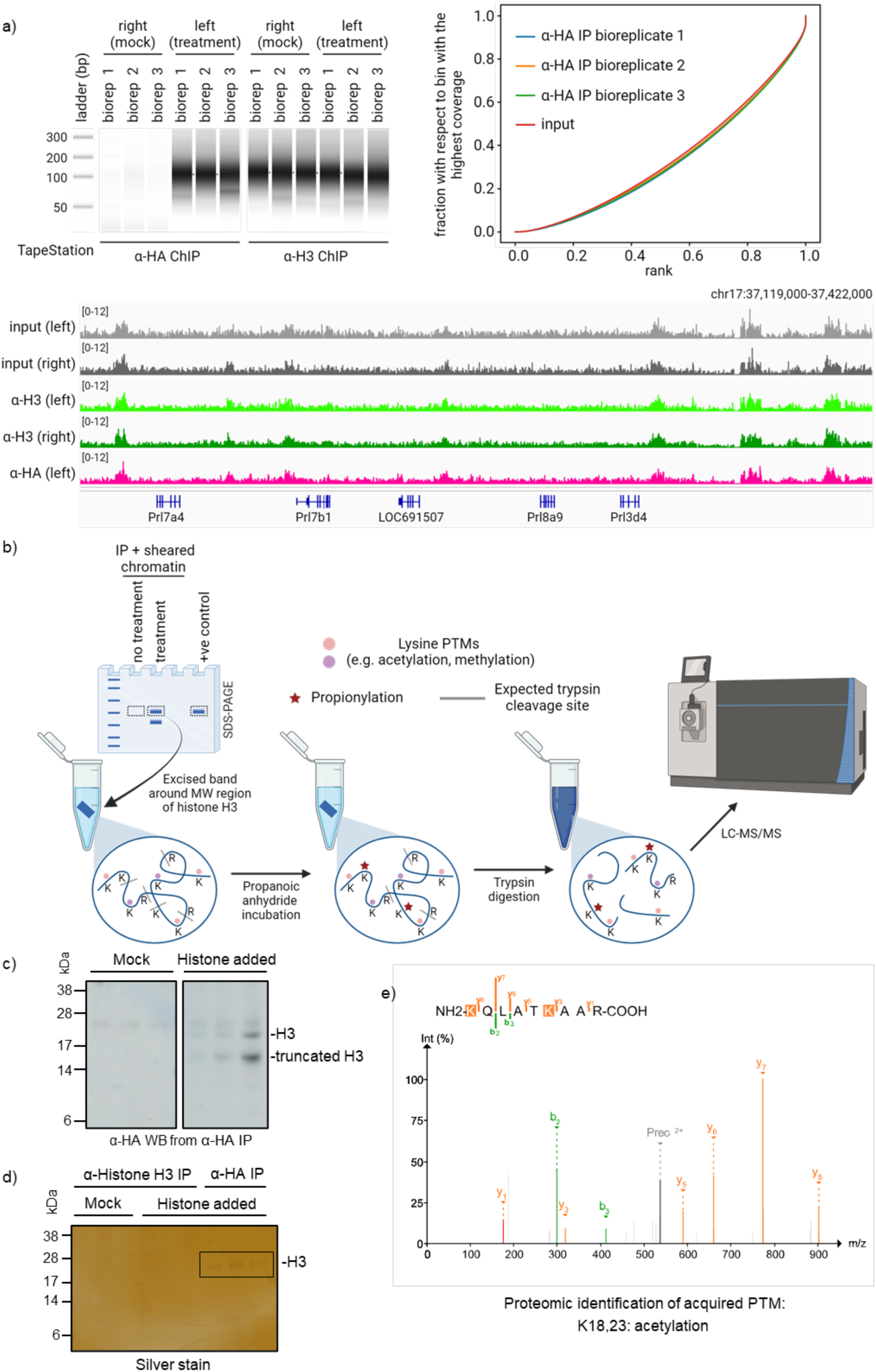
ChIP-seq and proteomic analysis of extracellular histones that incorporated into chromatin in rat brain. a) Tapestation analysis (left) reveals ∼100-fold enrichment of DNA from the HA-ChIP compared to mock treatment, with shearing patterns matching that of an endogenous histone H3 ChIP. ChIP-seq (right) reveals broad localization of HA peaks, suggesting stochastic deposition of extracellular histone H3 once it penetrated cells and incorporated into chromatin, indistinguishable from endogenous histone H3 or inpiut. Tracks (bottom) show that HA peaks from the treatment condition are indistinguishable from either endogenous H3 or input samples. b) The workflow for histone and histone PTM analyses by MS involved in-gel histone derivatization with propanoic anhydride prior to trypsin digestion. c) and d) Following immunoprecipitation after a 3 h (d) and 6 h (c) recovery post-injection, synthetic histone H3 was detected only in histone-treated tissue. e) In addition to gaining the histone PTM K27me2 (as with the 3 h treated tissue), proteomic analyses also revealed acetylation on K18 and K23 after prolonging the recovery period to 6 h (see also Figure 5e).

**Extended Data Figure 10:**
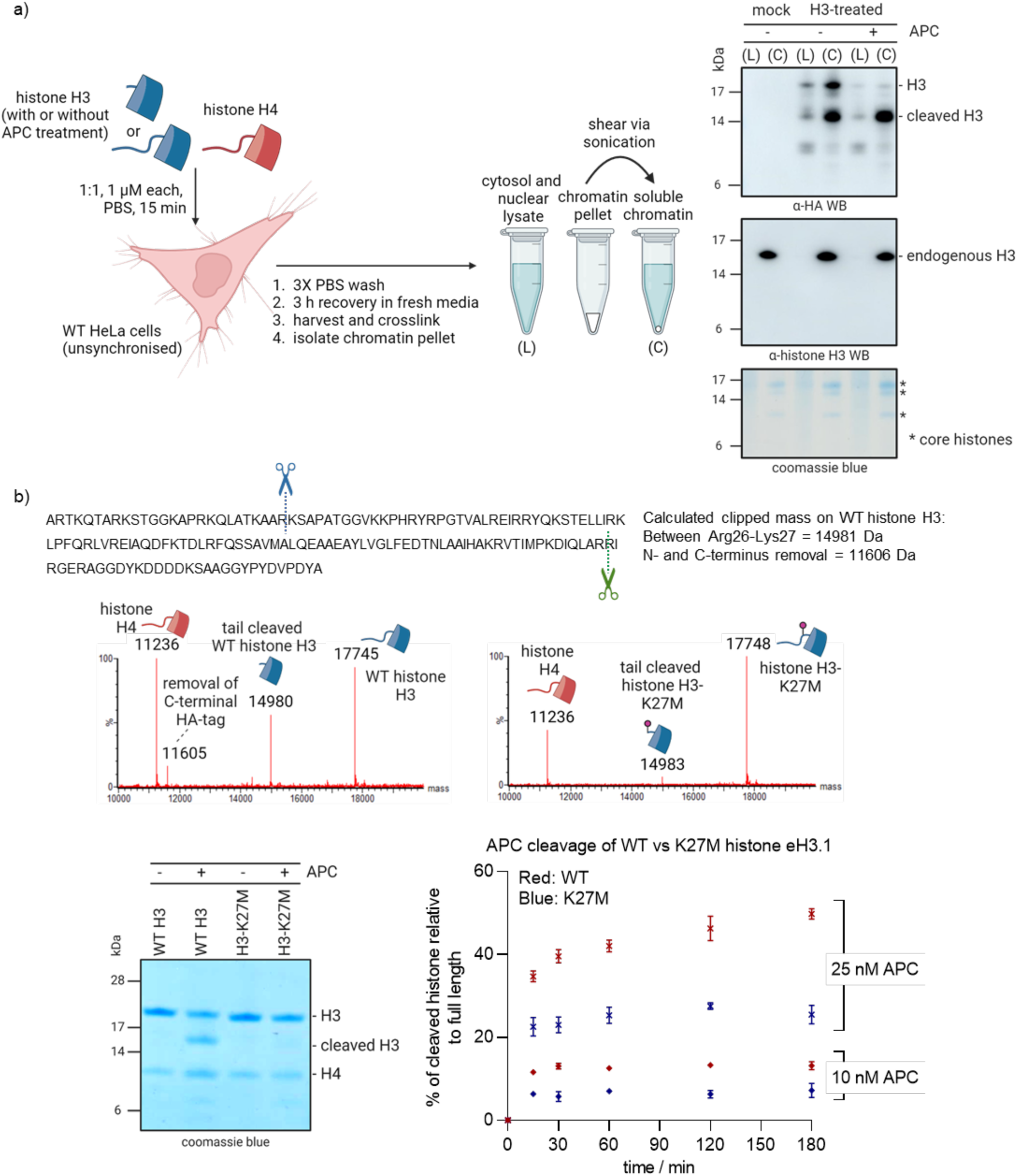
a) Tail-cleaved exogenous histone H3 (between Arg26 and Lys27) can still incorporate into chromatin. To generate tail-cleaved histone H3, 1:1 mixture of histones H3 and H4 (1 μM each) was incubated with Activated Protein C for 2 h at 37°C before incubation with HeLa cells. b) MS analysis of APC-treated histone H3 samples revealed a primary clipped mass from cleavage between Arg27-Lys27 (in blue) and, to a lesser extent, a secondary mass corresponding to likely removal of both the N- and C-terminal tails (in blue and green, respectively). Activated Protein C appeared to demonstrate lower cleavage activity on the K27M histone mutant (as a tetramer with histone H4) based on MS analysis and SDS-PAGE.

