## Supplementary Information for "Biomolecular tracking by FIRESCAPE reveals distinct modes of clearance, damage induction and cellular uptake for extracellular histone H3"

### *General Information*

#### Solvents and chemicals

Chemicals, solvents, media and *Escherichia coli* cell stocks were purchased from commercial suppliers in the United Kingdom (Acros, Alfa Aesar, Carbosynth, Fisher Scientific, Fluorochem, Sigma Aldrich, VWR) and used as received unless stated otherwise. Unless available as in-house supply, anhydrous solvents were purchased from Sigma Aldrich and stored under argon in a septum-capped bottle. All solvents were either analytical or HPLC grade. Reactions requiring anhydrous conditions were carried out with flame-dried reaction vessels under an inert atmosphere of nitrogen or argon.

#### Chromatography

Thin layer chromatography (TLC) was performed on Merck Millipore aluminium TLC plate, silica gel coated with fluorescent indicator F254 and visualised under UV irradiation at 254 nm, or by staining with aqueous potassium permanganate ($KMnO_4$ (3.00 g), $K_2CO_3$ (20.0 g), 5% NaOH (5 mL), $H_2O$ (300 mL)) with subsequent drying with a heat gun. Flash column chromatography was performed using Geduran® Silica Gel 60 (0.040-0.063 mm) or The Teledyne ISCO CombiFlash NextGen 100 Flash Chromatography System.

#### Nuclear Magnetic Resonance (NMR) spectrometry

All NMR spectra were recorded in commercially available deuterated solvents on a Bruker AVIIIHD 400 nanobay ($^{1}$H at 400, $^{13}$C at 101, $^{19}$F 376 MHz) or AVII HD 600 ($^{1}$H at 600, $^{13}$C at 151, $^{19}$F 565 MHz). All NMR data were processed using Mestrenova v.14.1.0. Chemical shifts are quoted in parts per million (ppm) relative to the centre of the solvent peak for $^{1}$H and $^{13}$C spectral data. All coupling constants are reported in Hz to the nearest 0.1 Hz. $^{13}$C spectra are $^{1}$H decoupled and $^{19}$F-$^{13}$C heteronuclear coupling on $^{13}$C spectra are reported. $^{19}$F NMR signals are referenced relative to $CFCl_3$.

### IR spectrometry

Infrared (IR) spectra of compounds were acquired on a Bruker Tensor 27 Fourier Transform spectrometer as crystals using a diamond attenuated total reflectance (ATR) attachment or as thin films of neat oils.

### Melting points

Melting points were measured using a Griffin apparatus on a Leica hotstage microscope and are uncorrected.

### Small-molecule mass spectrometry

Low resolution mass spectra were recorded on an Agilent 6120 Quadrupole spectrometer equipped with an electrospray ion source. High resolution mass spectra were obtained on a Thermo Exactive High-Resolution Orbitrap FTMS mass spectrometer with a lock-spray electrospray ion source. Methanol or acetonitrile were used as the carrier solvents. Some compounds were not stable under the MS ionisation methods and therefore, HRMS for these compounds were not obtained.

### Intact protein mass spectrometry general methods and data analysis

Protein reactions were monitored by LC-MS analysis of the crude reaction mixtures. Intact protein mass spectrometry was performed on Waters QTof mass spectrometers (Xevo G2-XS or Xevo G2-S) coupled to Acquity UHPLC systems. A Thermo Proswift column (250 mm x 4.6 mm x 5 μm) with a flow rate of 0.300 mL $min^{-1}$ and a solvent system of water + 0.1% formic acid (solvent A) and acetonitrile + 0.1% formic acid (solvent B) were employed for a total run time of 10 min with gradient elution as follows:

| Step | Time / min | %Solvent A | %Solvent B |
|---|---|---|---|
| 0 | 0 | 95 | 5 |
| 1 | 1.0 | 95 | 5 |
| 2 | 7.0 | 5 | 95 |
| 3 | 8.0 | 5 | 95 |
| 4 | 8.1 | 95 | 5 |
| 5 | 10.0 | 95 | 5 |

Nitrogen was used as the desolvation (650 L/h) and cone (30 L/h) gas with the following instrument parameters: capillary voltage 3 kV, cone voltage 20 V, source temperature 100 ºC, desolvation temperature 400 ºC and collision energy 6.0 eV.

The LC-MS chromatogram is based on total ion count. As the protein modification is miniscule in comparison to the entire protein, the retention times of different protein products will not vary. Hence, a protein peak on the chromatogram accounts for all protein species in the reaction i.e., starting material, desired product and unwanted by-products. Furthermore, appreciable changes in ionisation between different protein products are not expected. Hence, their relative intensities are used to calculate % conversion.

The raw spectra of multiple charged ion series were deconvoluted using Waters MassLynx software (v4.1) and its maximum entropy (MaxEnt1) function with the following set-up: resolution 1.00 Da/channel, width at half height (protein-dependent i.e., 0.40 Da for histones), 12 minimum intensity ratios 33% left and right and an iteration to convergence. The deconvolution mass range is protein dependent i.e. 10000 – 20000 Da for *X.l.* Histone H3, 10000 – 25000 Da for human histone eH3.1 and *X.l.* histone H3.NTEV. Reaction conversions were calculated as the relative intensity ratio of the peak of interest against the sum for all peaks. On histones, ~10% baseline methionine oxidation +16 Da adducts are often present during preparation and storage. These adducts were combined with their corresponding starting material or product.

### Biological reagents

Antibodies were used as per the manufacturer's recommendations. Primary antibodies: Anti-HA (C29F4) Rabbit mAb (Cell Signaling Technology, 3724) for HA-tag detection, Anti-Histone H3 (96C10) Mouse mAb (Cell signalling, 3680S) for histone detection. Secondary antibodies: Goat Anti-Rabbit IgG (H+L) HRP Conjugate (Promega, W4011), Goat Anti-Mouse IgG (H+L) HRP conjugate (Promega, W4021), Goat Anti-Mouse IgG H&L, Alkaline Phosphatase Conjugate (Abcam, ab97020), Goat Anti-Rabbit IgG (whole molecule), Alkaline Phosphatase Conjugate (Sigma, A3687). Human Activated Protein C (APC): ThermoFisher, RP-43095.

### Biological instruments

Sonication was performed using a Fisher Scientific Model 505 Sonic Dismembrator for protein purification, and a Bioruptor Pico (Dianogen) for chromatin shearing. Proteins were purified using an Äkta FPLC System UPC-900 (GE Healthcare, UK). Gel electrophoresis was performed using Invitrogen NuPAGE 10%, 12% or 4-12% Bis-Tris gels, Novex MiniCell tanks, and a BioRad PowerPac controller. Western blotting was performed using an iBlot gel transfer device from Thermo-Fisher. Colorimetric method (for alkaline phosphatase-conjugated antibody) was performed using Sigma BCIP/NBT Liquid Substrate System (catalogue number B1911). Chemiluminescent analysis (for horseradish peroxidase (HRP)-coupled antibody) was carried out using Thermo Scientific SuperSignal West Pico Plus Chemiluminescent substrate (catalogue number 34580). Nucleotide sequences were confirmed by the Source Bioscience DNA Sanger sequencing services based at Oxford University.

### Radiochemistry

#### General methods

For radiosynthesis carried out on the Advion Nanotek® radiosynthesizer, [$^{18}$F]fluoride was produced by Allliance Medical (UK), Invicro (UK) or PETIC (UK) via the $^{18}$O(p,n)$^{18}$F reaction and delivered as [$^{18}$F]fluoride in $^{18}$O-enriched-water. Radiosynthesis and azeotropic drying were performed on a NanoTek microfluidic device. HPLC analysis was performed with a Dionex Ultimate 3000 dual channel HPLC system equipped with shared autosampler, parallel UV-detectors and LabLogic NaI/PMT-radiodetectors with Flowram analog output. The radio signal is delayed by 0.1-0.3 min from the UV signal. Radio-TLC was performed on Merck Keiselgel 60 F254 plates with DCM/MeOH (9:1) elution system. Analysis was performed using a plastic scintillator/PMT detector. Radiochemical yield (RCY) for a protein reaction is the radio-fluorinated histone product expressed as a percentage against the remaining $^{18}$F-reagent. The values were determined by radio-HPLC of the crude protein reaction mixture. All isolated activity yields are non-decay corrected (n.d.c.).

[$^{18}$F]KF/$K_{2.2.2}$ elution on Advion Nanotek® radiosynthesizer: [$^{18}$F]Fluoride was separated from $^{18}$O-enriched-water using anion exchange cartridges (Waters Sep-Pak Accell Plus QMA Carbonate Plus Light Cartridge, pre-activated with $H_2O$ (10.0 mL, sorbent loading 46mg) and subsequently released with a solution of $K_{2.2.2}$/$K_2CO_3$ (Kryptofix® (7.5 mg in 600 μL of MeCN) and $K_2CO_3$ (1.5-2 mg in 150-200 μL of $H_2O$)) into the concentrator. The solution was further dried by azeotropic drying using anhydrous MeCN (3 x 500 μL) under a flow of $N_2$ at 105 °C.

For radiosynthesis carried out on the Trasis AllinOne radiosynthesizer, [$^{18}$F]fluoride was produced in-house (Turku PET Centre, Turku, Finland) by the $^{18}$O(p,n)$^{18}$F nuclear reaction using a CC-18/9 (Efremov Institut, Russia; 18 MeV) or a TR-19 (ACSI, Canada; 14-19 MeV) cyclotron and supplied as no-carrier added (n.c.a.) [$^{18}$F]fluoride in $^{18}$O-enriched-water.

**HPLC methods**

Method A: Synergi Hydro-RP C18 80 Å column was used (4 µm, 4.6 x 150 mm) with a flow rate of 1 mL min$^{-1}$, temperature of 25 °C and UV-Vis wavelength of 220 nm. A solvent system of $H_2O$/MeCN was used with the gradient elution as follows:

| Step | Time / min | % $H_2O$ | % MeCN |
|---|---|---|---|
| 0 | 0 | 95 | 5 |
| 1 | 1 | 95 | 5 |
| 2 | 10 | 5 | 95 |
| 3 | 14 | 5 | 95 |
| 4 | 16 | 95 | 5 |
| 5 | 18 | 95 | 5 |

Method B: Phenomenex Jupiter C4 300 Å column (5 µm, 4.6 x 250 mm) column was used with a flow rate of 1 mL min$^{-1}$, temperature of 25 °C and UV-Vis wavelength of 220 nm. A solvent system of $H_2O$ + 0.1% TFA (solvent A) and MeCN + 0.1% TFA (solvent B) was used with the gradient elution as follows:

| Step | Time / min | % Solvent A | % Solvent B |
|---|---|---|---|
| 0 | 0 | 95 | 5 |
| 1 | 2 | 95 | 5 |
| 2 | 22 | 5 | 95 |
| 3 | 24.5 | 5 | 95 |
| 4 | 25 | 95 | 5 |
| 5 | 30 | 95 | 5 |

### *Reagent synthesis*

#### 2-((bromomethyl)thio)benzothiazole

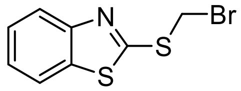

A solution of 2-benzothiazolethiol (168 mg, 1.00 mmol) in THF (1 mL) was added dropwise over 20 minutes to a mixture of $Cs_2CO_3$ (491 mg, 1.51 mmol) in dibromomethane (5 mL) and MeCN (1.5 mL) while stirring at rt. After complete addition, the reaction was monitored by TLC every 10 minutes for complete consumption of 2-mercaptobenzothiazole. The reaction mixture was then filtered and the filtrate was concentrated. Combiflash purification by silica gel column chromatography (0 to 50% DCM in pet. ether over 10 min) gave a white solid (75 mg, 29%).

$C_8H_6BrNS_2$ (258.9 g/mol):
**TLC:** $R_f$ = 0.8 (70% DCM/pet. ether)
**$^1$H NMR** (400 MHz, $CDCl_3$) δ 7.98 (ddd, *J* = 8.2, 1.2, 0.6 Hz, 1H), 7.82 (ddd, *J* = 8.0, 1.2, 0.6 Hz, 1H), 7.48 (ddd, *J* = 8.3, 7.2, 1.2 Hz, 1H), 7.37 (ddd, *J* = 8.0, 7.2, 1.2 Hz, 1H), 5.22 (s, 2H). **$^{13}$C NMR** (101 MHz, $CDCl_3$) δ 162.7, 153.0, 135.7, 126.7, 125.1, 122.5, 121.4, 31.3. **IR** (ATR): $\tilde{\upsilon}$/cm$^{-1}$ = 3027, 2921, 2851, 1555, 1454, 1422, 1365, 1309, 1275, 1234, 1192, 1128, 1069, 1018, 999, 941, 807, 758, 728, 178.

### Ethyl 2-(benzothiazole-2-ylthio)-2-fluoroacetate

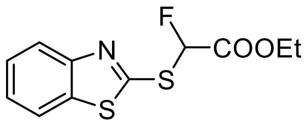

The following procedure was adapted from a known procedure.[1]

Triethylamine (2.08 mL, 15.0 mmol) was added dropwise to a mixture of 2-benzothiazolethiol (2.50 g, 15.0 mmol) in EtOH (35 mL) under nitrogen. The solution was allowed to stir at rt for 15 min before adding ethyl bromofluoroacetate (1.95 mL, 16.4 mmol) and leaving to stir for a further 16 h. The crude mixture was quenched with 1 M HCl (20 mL) and extracted with DCM (3 x 50 mL) and the combined organic layers were washed with $H_2O$ (50 mL) and brine (50 mL). The organic phase was then dried over $MgSO_4$, filtered and concentrated under reduced pressure. Purification by silica gel column chromatography (5% EtOAc in pet. ether) gave a colourless oil (3.85 g, 95%).

$C_{11}H_{10}FNO_2S_2$ (271.3 g/mol):
**TLC:** $R_f$ = 0.7 (20% EtOAc/pet. ether)
**$^1$H NMR** (400 MHz, $CDCl_3$) δ 7.97 (ddd, *J* = 8.1, 1.2, 0.6 Hz, 1H), 7.81 (ddd, *J* = 8.0, 1.3, 0.6 Hz, 1H), 7.47 (ddd, *J* = 8.3, 7.3, 1.3 Hz, 1H), 7.37 (ddd, *J* = 8.3, 7.2, 1.2 Hz, 1H), 6.94 (d, *J* = 51 Hz, 1H), 4.33 (q, *J* = 7.1 Hz, 2H), 1.32 (t, *J* = 7.2 Hz, 3H). **$^{19}$F NMR** (376 MHz, $CDCl_3$) δ -161.1 (d, *J* = 51 Hz).

Spectroscopic data was consistent with literature reports.[2]

### Ethyl 2-(benzothiazole-2-ylsulfonyl)-2-fluoroacetate

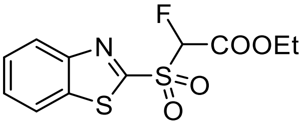

Following general procedure 3, ethyl 2-(benzothiazole-2-ylthio)-2-fluoroacetate was prepared from a mixture of the starting sulfide (400 mg, 1.47 mmol), $NaIO_4$ (1.58 g, 7.37 mmol) and ruthenium trichloride hydrate (3 mg). Purification by column chromatography (0 to 60% EtOAc in pet. ether over 10 min) gave the sulfone product as a yellow solid (272 mg, 61%)

$C_{11}H_{10}FNO_4S_2$ (303.0 g/mol):
**TLC:** $R_f$ = 0.3 (20% EtOAc/pet. ether)
**$^1$H NMR** (400 MHz, $CDCl_3$) δ 8.30 – 8.24 (m, 1H), 8.08 – 8.01 (m, 1H), 7.71 – 7.62 (m, 2H), 6.04 (d, *J* = 47.5 Hz, 1H), 4.47 – 4.34 (m, 2H), 1.34 (t, *J* = 7.1 Hz, 3H). **$^{19}$F NMR** (376 MHz, $CDCl_3$) δ -181.0 (d, *J* = 47.6 Hz).

Spectroscopic data was consistent with literature reports.[2]

### 2-((fluoromethyl)sulfonyl)benzothiazole

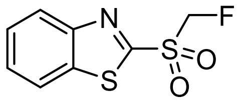

The following procedure was adapted from a known procedure.[3]

To a solution of $BzSO_2CFHCO_2Et$ (245 mg, 808 μmol) in EtOAc (15 mL) was added DBU (170 μL, 1.13 mmol) and $H_2O$ (50 μL). The resulting mixture was allowed to stir at rt for 16 h before quenching with sat. aq. $NH_4Cl$ (5 mL). The organic layer was separated and the aqueous layer was extracted with DCM (2 x 10 mL). The combined organic layers were washed with brine (20 mL) and then dried over $MgSO_4$, filtered and concentrated under reduced pressure. Purification was performed instead by trituration in a cold mixture of ether:DCM (3:1) to a white solid (95 mg, 51%).

$C_8H_6FNO_2S_2$ (231.0 g/mol):
**$^1$H NMR** (400 MHz, DMSO-$d_6$) δ 8.44 – 8.36 (m, 1H), 8.37 – 8.29 (m, 1H), 7.82 – 7.72 (m, 2H), 6.14 (d, $J$ = 45 Hz, 2H). **$^{19}$F NMR** (376 MHz, DMSO-$d_6$) δ -208.3 (t, $J$ = 45 Hz). **MS$^+$** 232.0 $[M + H]^+$

Spectroscopic data was consistent with literature reports.[4]

### tert-Butyl (3-(benzothiazol-2-ylthio)propyl)carbamate

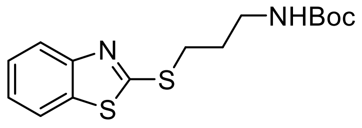

Under argon, 2-benzothiazolethiol (1.69 g, 10.1 mmol), 3-(boc-amino)propyl bromide (2.00 g, 8.40 mmol) and $K_2CO_3$ (2.32 g, 16.8 mmol) were dissolved in dry MeCN (50 mL). NaI (60 mg) was then added and the solution was stirred for 16 h at rt. The reaction mixture was concentrated and then diluted with $H_2O$ (50 mL) and $Et_2O$ (100 mL). The organic was separated and the aqueous phase was extracted with $Et_2O$ (2 x 50 mL). The combined organic layer was washed with sat. aq. $NH_4Cl$ (50 mL) and brine (50 mL) then dried over $MgSO_4$, filtered and concentrated under reduced pressure. The *N*-boc protected amine was used directly in the next step.

### tert-Butyl (3-(benzothiazol-2-ylthio)propyl)(tert-butoxycarbonyl)carbamate

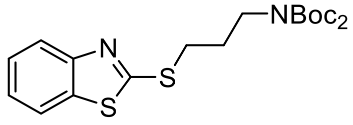

Under argon, the *N*-boc protected amine was dissolved in dry MeCN (17 mL). $Boc_2O$ (3.67 g, 16.8 mmol), DMAP (103 mg, 0.84 mmol) and $NEt_3$ (3.52 mL, 25.2 mmol) were then added and the solution was stirred for 16 h at rt. The reaction mixture was concentrated and then diluted with sat. aq. $NH_4Cl$ (50 mL) and EtOAc (100 mL). The organic phase was separated and the aqueous phase was extracted with EtOAc (2 x 50 mL). The combined organic layer was washed with sat. aq. $NH_4Cl$ (50 mL) and brine (50 mL) then dried over $MgSO_4$, filtered and concentrated under reduced pressure. Silica gel column chromatography (5% EtOAc in pet. ether) gave the *N,N*-diboc protected amine as a white solid (1.68 g, 44% over 2 steps). The yield was lower than expected as twice the amount of DMAP i.e. 1.68 mmol, should have been used.

$C_{20}H_{28}N_2O_4S_2$ (424.6 g/mol):

**TLC:** $R_f$ = 0.4 (10% EtOAc/pet. ether)

**$^1$H NMR** (400 MHz, $CDCl_3$) δ 7.85 (ddd, *J* = 8.1, 1.2, 0.6 Hz, 1H), 7.75 (ddd, *J* = 8.0, 1.3, 0.6 Hz, 1H), 7.40 (ddd, *J* = 8.3, 7.3, 1.3 Hz, 1H), 7.29 (ddd, *J* = 7.9, 7.0, 1.2 Hz, 1H), 3.76 (t, *J* = 7.0 Hz, 2H), 3.37 (t, *J* = 7.2 Hz, 2H), 2.12 (p, *J* = 7.1 Hz, 2H), 1.50 (s, 18H). **$^{13}$C NMR** (151 MHz, $CDCl_3$) δ 166.8, 153.4, 152.6 (2C), 135.3, 126.1, 124.3, 121.6, 121.1, 82.6 (2C), 45.4, 31.0, 29.0, 28.2 (6C). **IR** (ATR): $\tilde{\upsilon}$/cm$^{-1}$ = 3063, 2975, 2933, 1728, 1685, 1460, 1445, 1428, 1398, 1354, 1210, 1291, 1250, 1219, 1171, 1157, 1133, 1109, 1036, 1017, 1001, 973, 933, 892, 858, 830, 796, 788, 772, 753, 724, 704. **HRMS** (ESI (+), MeOH): (*m/z*) calc. for $C_{20}H_{28}N_2O_4S_2$: 447.1383 $[M + Na]^+$; found: 447.1390.

tert-Butyl (3-(benzothiazol-2-ylthio)-3-chloropropyl)(tert-butoxycarbonyl)carbamate

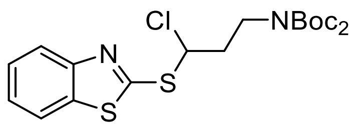

Under argon, the *N,N*-diboc protected amine (850 mg, 2.00 mmol) was dissolved in dry benzene (10 mL) in a sealed vial. The reaction was allowed to stir for 5 min at 50 °C. *N*-chlorosuccinimide (401 mg, 3.00 mmol) was then added in one portion. The solution was left to stir for a further 3 h at 50 °C before the solvent was removed under a flow of $N_2$. Silica gel column chromatography (3% EtOAc in pet. ether) gave the α-chlorosulfide product as a white solid (640 mg, 70%).

$C_{20}H_{27}ClN_2O_4S_2$ (459.0 g/mol):
**TLC:** $R_f$ = 0.4 (10% EtOAc/pet. ether)
**$^1$H NMR** (400 MHz, $CDCl_3$) δ 7.95 (ddd, *J* = 8.2, 1.2, 0.6 Hz, 1H), 7.81 (ddd, *J* = 8.0, 1.3, 0.6 Hz, 1H), 7.46 (ddd, *J* = 8.3, 7.2, 1.3 Hz, 1H), 7.36 (ddd, *J* = 8.3, 7.3, 1.2 Hz, 1H), 6.10 (dd, *J* = 7.6, 5.4 Hz, 1H), 3.90 (t, *J* = 7.2 Hz, 2H), 2.60 – 2.44 (m, 2H), 1.51 (s, 18H). **$^{13}$C NMR** (151 MHz, $CDCl_3$) δ 162.2, 153.1, 152.3 (2C), 135.8, 126.5, 125.1, 122.5, 121.3, 83.0 (2C), 64.2, 43.8, 38.4, 28.2 (6C). **IR** (ATR): $\tilde{\upsilon}$/cm$^{-1}$ = 3059, 2979, 2923, 1719, 1682, 1558, 1455, 1439, 1427, 1398, 1356, 1348, 1313, 1285, 1259, 1243, 1213, 1135, 1080, 1041, 1017, 1007, 945, 891, 860, 847, 809, 788, 773, 755, 730, 703. **MS$^+$** 481.0 [M + Na]$^+$ **HRMS** (ESI (+), MeOH): (*m/z*) calc. for $C_{20}H_{27}ClN_2O_4S_2$: 481.0993 [M + Na]$^+$; found: 481.1013

### tert-Butyl (3-(benzothiazol-2-ylsulfonyl)propyl)carbamate

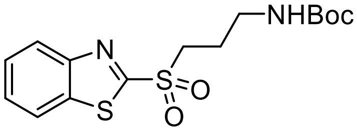

To a solution of 2-benzothiazolethiol (2.92 g, 17.4 mmol) in MeCN (100 mL) was added $K_2CO_3$ (4.80 g, 34.8 mmol), NaI (350 mg, 2.33 mmol) and 3-(boc-amino)propyl bromide (5.00 g, 20.9 mmol). The reaction mixture was stirred for 16 h before being diluted with EtOAc (250 mL) and washed with water (150 mL), saturated aqueous $NH_4Cl$ (150 mL), saturated aqueous NaCl (150 mL) before being dried ($Na_2SO_4$), filtered and concentrated *in vacuo* to yield a yellow oil.

To a solution crude thioether (17.4 mmol) in DCM (200 mL) at 0 °C was added *m*CPBA (11.7 g, 50.2 mmol, 77% by weight). The mixture was stirred at this temperature for 5 h before a further portion of *m*CPBA (3.0 g, 13.38 mmol, 77% by weight) was added and the reaction stirred for 16 h. At this point the reaction mixture was cooled to 0 °C and quenched with 10% aqueous $Na_2S_2O_3$ (200 mL) and diluted with DCM (200 mL). The organic phase was washed with saturated aqueous $NaHCO_3$ (5 x 400 mL), saturated aqueous NaCl (300 mL) before being dried ($Na_2SO_4$), filtered and concentrated *in vacuo*. The crude product was then purified by flash chromatography (4% to 5% EtOAc in DCM) to yield the desired sulfone as a white solid (4.85 g, 78% yield over two steps).

$C_{15}H_{20}N_2O_4S_2$ (356.5 g/mol):
**$^1$H NMR** (400 MHz, $CDCl_3$) δ 8.24 – 8.19 (m, 1H), 8.05 – 7.99 (m, 1H), 7.69 – 7.56 (m, 2H), 4.73 (br s, 1H), 3.61 – 3.54 (m, 2H), 3.30 (q, J = 6.5 Hz, 2H), 2.10 (dq, J = 9.3, 6.7 Hz, 2H), 1.42 (s, 9H). **$^{13}$C NMR** (101 MHz, $CDCl_3$) δ 165.8, 156.0, 152.8, 136.9, 128.3, 127.9, 125.7, 122.5, 79.9, 52.4, 38.9, 28.5 (3C), 23.5. **MS$^+$** 379.0 [M + Na]$^+$ **HRMS** (ESI (+), MeOH): (*m/z*) calc. for $C_{15}H_{20}N_2O_4S_2$: 379.0757 [M + Na]$^+$; found: 379.0758

### tert-Butyl (3-(benzothiazol-2-ylsulfonyl)-3-fluoropropyl)carbamate

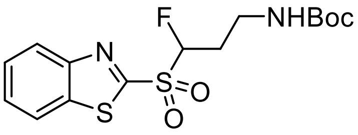

Under argon, *tert*-butyl (3-(benzothiazole-2-ylsulfonyl)propyl)carbamate (890 mg, 2.50 mmol) was dissolved in anhydrous THF (12.5 mL). The solution was cooled to -78 °C and left to stir for 10 min. 1 M LiHMDS in anhydrous THF (5 mL, 5.00 mmol) was added dropwise and the solution was stirred for a further 5 min). 0.4 M NFSI in anhydrous THF (7.50 mL, 3.00 mL) was then added and the resulting mixture was left to stir at -78 °C for 45 min. The reaction was quenched with sat. aq. $NH_4Cl$ (20 mL) and $Et_2O$ (20 mL) at -78 °C before allowing the mixture to reach rt. The organic layer phase was separated and the aqueous layer was extracted with $Et_2O$ (2 x 25 mL). The combined organic layers was washed with sat. aq. $NH_4Cl$ (25 mL) then dried over $MgSO_4$, filtered and concentrated under reduced pressure. Purification by silica gel column chromatography (10% $Et_2O$ in DCM) gave *tert*-butyl (3-(benzothiazole-2-ylsulfonyl)-3-fluoropropyl)carbamate as a white solid (433 mg, 46%).

$C_{15}H_{19}FN_2O_4S_2$ (374.5 g/mol):
**$^1$H NMR** (400 MHz, $CDCl_3$) δ 8.31 – 8.24 (m, 1H), 8.08 –8.01 (m, 1H), 7.70 – 7.61 (m, 2H), 5.79 (ddd, *J* = 48.2, 9.5, 3.5 Hz, 1H), 4.76 (br s, 1H), 3.51 – 3.41 (m, 2H), 2.62 – 2.44 (m, 1H), 2.44 – 2.28 (m, 1H) 1.45 (s, 9H). **$^{19}$F{$^1$H} NMR** (377 MHz, $CDCl_3$) δ -178.72. **$^{13}$C NMR** (101 MHz, $CDCl_3$) δ 162.2, 156.0, 152.9, 137.5, 128.5, 128.0, 125.8, 122.4, 100.6 (d, *J* = 222.0 Hz), 79.9, 35.9, 28.4 (3C), 28.1 (d, *J* = 19.0 Hz). **IR** (ATR): $\tilde{\upsilon}$/cm$^{-1}$ = 3391, 2971, 2361, 2341, 1696, 1520, 1472, 1438, 1417, 1391, 1366, 1337, 1318, 1292, 1271, 1250, 1229, 1170, 1153, 1085, 1025, 1001, 986, 876, 847, 784, 760, 727. **MS$^+$** 397.1 $[M + Na]^+$ **HRMS** (ESI (+), MeOH): (*m/z*) calc. for $C_{15}H_{19}FN_2O_4S_2$: 397.0663 $[M + Na]^+$; found: 397.0664

### tert-Butyl 3-(benzothiazol-2-ylsulfonyl)-3-fluoropropan-1-amine hydrochloride

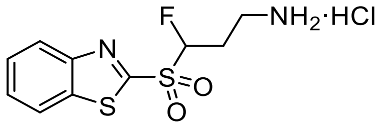

*Tert*-butyl (3-(benzothiazole-2-ylsulfonyl)-3-fluoropropyl)carbamate (187 mg, 0.499 mmol) was dissolved in DCM (5 mL). 4 M HCl in dioxane (1.25 mL, 4.99 mmol) was added and the mixture was left to stir at rt for 2 h (or until complete consumption of the starting material monitored by TLC). Solvents were removed under a flow of $N_2$ to give *tert*-butyl 3-(benzothiazole-2-ylsulfonyl)-3-fluoropropan-1-amine hydrochloride as a white solid (quant. yield).

$C_{10}H_{12}ClFN_2O_2S_2$ (310.8 g/mol):
**$^1$H NMR** (600 MHz, MeOD) δ 8.30 – 8.22 (m, 2H), 7.77 – 7.71 (m, 2H), 6.15 (ddd, *J* = 47.3, 9.1, 3.5 Hz, 1H), 3.34 – 3.23 (m, 2H), 2.75 – 2.63 (m, 1H), 2.59 – 2.49 (m, 1H). **$^{19}$F NMR** (565 MHz, MeOD) δ -181.27 (ddd, *J* = 47.3, 32.0, 16.7). **$^{13}$C NMR** (151 MHz, MeOD) δ 163.6, 154.2, 138.8, 130.0, 129.4, 126.5, 124.1, 101.3 (d, *J* = 219.5 Hz), 36.3 (d, *J* = 4.0 Hz), 26.9 (d, *J* = 19.6 Hz). **IR** (ATR): $\tilde{\upsilon}$/cm$^{-1}$ = 3370, 2922, 2360, 2342, 2089, 2030 ,2003, 1961, 1636, 1541, 1515, 1465, 1410, 1350, 1321, 1284, 1238, 1155, 1078, 1028, 913, 857, 764, 729, 717, 701. **MS$^+$** 274.7 [M]$^+$ **HRMS** (ESI (+), MeOH): (*m/z*) calc. for $C_{10}H_{12}FN_2O_2S_2$: 275.0319 [M]$^+$; found: 275.0325

tert-Butyl (3-(benzothiazol-2-ylsulfonyl)-3-fluoropropyl)(tert-butoxycarbonyl)carbamate

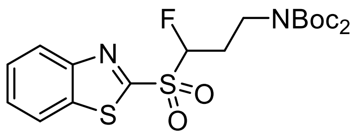

Under argon, *tert*-butyl (3-(benzothiazole-2-ylsulfonyl)-3-fluoropropyl)carbamate (112 mg, 299 µmol) was dissolved in dry MeCN (0.6 mL). $Boc_2O$ (131 mg, 598 µmol), DMAP (7.30 mg, 59.8 µmol) and $NEt_3$ (125 µL, 897 µmol) were then added and the solution was stirred for 16 h at rt. The reaction mixture was concentrated and then diluted with sat. aq. $NH_4Cl$ (3 mL) and EtOAc (5 mL). The organic phase was separated and the aqueous phase was extracted with EtOAc (2 x 5 mL). The combined organic layer was washed with sat. aq. $NH_4Cl$ (5 mL) and brine (5 mL) then dried over $MgSO_4$, filtered and concentrated under reduced pressure. Combiflash purification by silica gel column chromatography (0 to 40% EtOAc in pet. ether over 12 min) gave the *N,N*-diboc protected amine as a white solid (100 mg, 70%).

$C_{20}H_{27}FN_2O_6S_2$ (474.6 g/mol):
**TLC:** $R_f$ = 0.3 (20% EtOAc/pet. ether)
**$^1$H NMR** (400 MHz, $CDCl_3$) δ 8.29 – 8.22 (m, 1H), 8.07 –8.00 (m, 1H), 7.70 – 7.59 (m, 2H), 5.81 (ddd, *J* = 48.1, 9.9, 2.6 Hz, 1H), 3.98 – 3.83 (m, 2H), 2.60 (ddtd, *J* = 36.9, 14.5, 7.1, 2.7 Hz, 1H), 2.49 – 2.32 (m, 1H), 1.50 (18H, s). **$^{19}$F NMR** (376 MHz, $CDCl_3$) δ 178.0 (ddd, *J* = 48.3, 36.8, 16.4 Hz). **$^{13}$C NMR** (101 MHz, $CDCl_3$) 162.4, 153.0, 152.3 (2C), 137.6, 128.6, 128.0, 125.9, 122.5, 100.7 (d, *J* = 222.4 Hz), 83.3 (2C), 41.7 (d, *J* = 2.3 Hz), 28.2 (6C), 27.2 (d, *J* = 19.2 Hz). **IR** (ATR): $\tilde{\upsilon}$/cm$^{-1}$ = 2982, 2935, 1739, 1714, 1698, 1553, 1467, 1441, 1408, 1394, 1370, 1350, 1337, 1317, 1305, 1279, 1253, 1235, 1215, 1156, 1112, 1078, 1051, 1024, 998, 948, 924, 896, 852, 833, 804, 770, 760, 729. **MS$^+$** 497.11 $[M + Na]^+$ **HRMS** (ESI (+), MeOH): (*m/z*) calc. for $C_{20}H_{27}FN_2O_6S_2$: 497.1187 $[M + Na]^+$; found: 497.1194

#### *Protein production and characterisation*

All human canonical core histones (eH3.1, H4, H2A and H2B) were expressed and purified as previously described[5] ('e' on histone eH3.1 refers to the C-terminal FLAG-HA dual epitope tag). Briefly, histones (cloned previously into pET3 expression vectors[5]) were expressed in either BL21 (DE3) CodonPlus RIPL *E. coli* cells for H2B, or BL21 (DE3) pLysS *E.coli* cells for eH3.1 WT, H4 and H2A. Cells were transformed and expanded to 4 x 1 L in 2xYT media containing carbenicillin and chloramphenicol and grown at 37 °C until $OD_{600}$ = 0.4 - 0.6. IPTG was added to a final concentration of 0.5 mM and the flask was shaken at 37 °C for 2 h. Cells were harvested by centrifugation for 15 min at 8k rpm at 4 °C and the pellet was resuspended into a 5-fold v/w wash buffer (50 mM Tris pH 7.5, 100 mM NaCl with protease inhibitors). Suspensions were flash-frozen and stored at -80 °C until lysis. Frozen cells were thawed in a water bath. When the solution was viscous, DNase was added (1 mg), followed by sonication (5× at 40% amplitude in 30 s bursts) and then centrifuged at 13k rpm for 20 min at 4 °C. The supernatant was discarded, and the inclusion body pellet was resuspended in 40 mL wash buffer. Sonication was repeated twice at 40% amplitude for 30 s, and the suspension centrifuged at 13k rpm for 20 min after each wash. After addition of 1 mL DMSO, the pellet was mixed with a spatula and left at RT for 10 min. 10 mL unfolding buffer (7 M Gdn·HCl, 20 mM Tris pH 7.5, 10 mM DTT) was then added, and shaken for 1 h at RT, after which it was centrifuged for 20 min at 13k rpm at RT. The supernatant was concentrated to 1.5 mL and then filtered before loading onto an S200 size-exclusion column pre-equilibrated with 1 CV SAU-100 (7 M urea, 20 mM NaOAc, pH 5.2, 100 mM NaCl, 1 mM EDTA, 10 mM DTT). Histone proteins were eluted with SAU-100 buffer and analysed by SDS-PAGE. Fractions containing the desired histones were pooled. Further purification by cation exchange chromatography with HiTrap SP 5 mL was performed using a linear gradient of 0-100% SAU-1000 buffer (SAU-100 with 1000 mM NaCl final concentration). Pure fractions were pooled, dialyzed thrice against water containing 2 mM βME and lyophilized.

### Histone eH3.1 sequences

Human histone eH3.1-WT

Mutations: C96A, C110A, C-terminal FLAG-HA epitope tags (e)

```
        10         20         30         40         50         60
ARTKQTARKS TGGKAPRKQL ATKAARKSAP ATGGVKKPHR YRPGTVALRE IRRYQKSTEL

        70         80         90        100        110        120
LIRKLPFQRL VREIAQDFKT DLRFQSSAVM ALQEAAEAYL VGLFEDTNLA AIHAKRVTIM

       130        140        150
PKDIQLARRI RGERAGGDYK DDDDKSAAGG YPYDVPDYA
```

Calculated mass: 17745 g $mol^{-1}$

Observed mass: 17745 g $mol^{-1}$

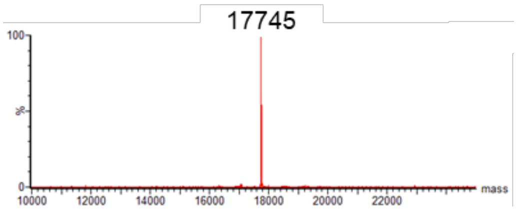

Human histone eH3.1-Cys4

Mutations: K4C, C96A, C110A, C-terminal FLAG-HA epitope tags (e)

```
        10         20         30         40         50         60
ARTCQTARKS TGGKAPRKQL ATKAARKSAP ATGGVKKPHR YRPGTVALRE IRRYQKSTEL

        70         80         90        100        110        120
LIRKLPFQRL VREIAQDFKT DLRFQSSAVM ALQEAAEAYL VGLFEDTNLA AIHAKRVTIM

       130        140        150
PKDIQLARRI RGERAGGDYK DDDDKSAAGG YPYDVPDYA
```

Calculated mass: 17720 g $mol^{-1}$

Observed mass: 17720 g $mol^{-1}$

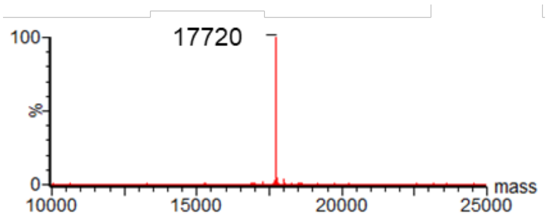

Human histone eH3.1-Cys27

Mutations: K27C, C96A, C110A, C-terminal FLAG-HA epitope tags (e)

```
        10         20         30         40         50         60
ARTKQTARKS TGGKAPRKQL ATKAARCSAP ATGGVKKPHR YRPGTVALRE IRRYQKSTEL

        70         80         90        100        110        120
LIRKLPFQRL VREIAQDFKT DLRFQSSAVM ALQEAAEAYL VGLFEDTNLA AIHAKRVTIM

       130        140        150
PKDIQLARRI RGERAGGDYK DDDDKSAAGG YPYDVPDYA
```

Calculated mass: 17720 g $mol^{-1}$

Observed mass: 17720 g $mol^{-1}$

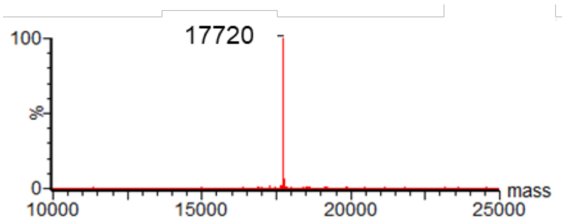

Human histone eH3.1-Cys56

Mutations: K56C, C96A, C110A, C-terminal FLAG-HA epitope tags (e)

```
        10         20         30         40         50         60
ARTKQTARKS TGGKAPRKQL ATKAARKSAP ATGGVKKPHR YRPGTVALRE IRRYQCSTEL

        70         80         90        100        110        120
LIRKLPFQRL VREIAQDFKT DLRFQSSAVM ALQEAAEAYL VGLFEDTNLA AIHAKRVTIM

       130        140        150
PKDIQLARRI RGERAGGDYK DDDDKSAAGG YPYDVPDYA
```

Calculated mass: 17720 g $mol^{-1}$

Observed mass: 17720 g $mol^{-1}$

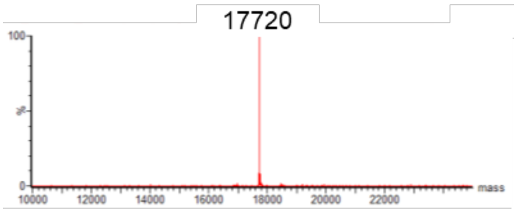

Human histone eH3.1-Met27

Mutations: K27M, C96A, C110A, C-terminal FLAG-HA epitope tags (e)

```
         10         20         30         40         50         60
ARTKQTARKS TGGKAPRKQL ATKAARMSAP ATGGVKKPHR YRPGTVALRE IRRYQKSTEL

         70         80         90        100        110        120
LIRKLPFQRL VREIAQDFKT DLRFQSSAVM ALQEAAEAYL VGLFEDTNLA AIHAKRVTIM

        130        140        150
PKDIQLARRI RGERAGGDYK DDDDKSAAGG YPYDVPDYA
```

Calculated mass: 17748 g $mol^{-1}$

Observed mass: 17748 g $mol^{-1}$

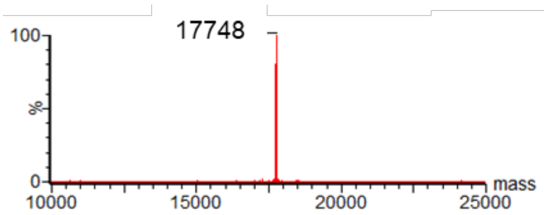

### Histone octamer assembly

Histones eH3.1 WT, H4, H2B and H2A were dissolved in unfolding buffer (7 M Gdn·HCl, 20 mM Tris pH 7.5, 10 mM DTT) and shaken at room temperature for at least 15 min (500 rpm). All four histones were mixed in an equimolar concentration of 38 µM, in a volume of approximately 1.5 mL. This mixture was dialysed against refolding buffer (2 M NaCl, 10 mM Tris, 1 mM EDTA, pH 7.4) three times at 4 °C and then concentrated to 1 mL and loaded onto a Superdex 16/60 pg75 size-exclusion column pre-equilibrated with refolding buffer. Fractions containing all four histones were combined and concentrated to 1.17 mg $mL^{-1}$ (10.3 µM) at 4 °C.

### Large scale 145bp Widom 601 DNA preparation

Based on literature protocol[6], the pUC57 8 × 145 bp Widom 601 plasmid was transformed into XL10-Gold cells, expanded to 8 x 1 L TB containing ampicillin and grown for 24 h at 37 °C. Cells were then harvested by centrifugation and resuspended in 80 mL of alkaline lysis buffer 1 per litre (50 mM glucose, 25 mM Tris, 10 mM EDTA, pH 8). To each tube, 120 mL of alkaline lysis buffer 2 was added (200 mM NaOH, 1% (w/v) SDS). The pellets were shaken vigorously until homogenous, then incubated for 20 minutes on ice. 210 mL of ice cold alkaline lysis buffer 3 was then added (4 M NaOAc, 2 M acetic acid) and the mixture was incubated on ice for 20 minutes. The mixture was centrifuged (1000 g, 20 min, 4 °C) and the cleared supernatant was then filtered through a miracloth before 0.52 v/v of isopropanol was added. The mixture was left to stand at room temperature for 15 min and then centrifuged again. The pellet was air-dried and resuspended in 25 mL TE 10/50 (10 mM Tris, 50 mM EDTA, pH 8). 120 µL RNAse A was added and incubated at 37 °C overnight. The mixture was centrifuged, and DNA was precipitated from the supernatant by adding one-fifth of 4 M NaCl and two-fifths 40% PEG 6000. The supernatant was shaken at 37 °C for 5 min and then incubated on ice for 30 min to allow full precipitation. DNA was harvested by centrifugation (3000 g, 20 min, 4 °C), and the pellet was dissolved in 15 mL TE 10/0.1 (10mM Tris, 0.1 mM EDTA, pH 8) by shaking overnight at 37 °C. 10,000 units of EcoRV were added and incubated at 37 °C for 20 h. 0.192 volume of 4 M NaCl and 0.346 volume of 40% PEG6000 were added and the mixture was incubated on ice for 1 h. The precipitated vector was harvested by centrifugation (27000 g, 20 min,

4 °C), and the pellet was dissolved in 2 mL TE 10/0.1. Both the vector fraction and the pellet fraction were analysed on a 6% TBE gel stained with SYBR safe. The supernatant was added to 125 mL of ice cold ethanol and incubated on ice for 20 min. The precipitated insert was harvested by centrifugation (27000 g, 20 min, 4 °C) and dissolved in 5 mL TE 10/0.1. 0.4 volume of phenol and 0.2 volume of chloroform were then added to the dissolved insert solution. The mixture was shaken vigorously and centrifuged until the layers were separated. The aqueous layer was extracted and mixed 1:1 with chloroform. After centrifugation, the aqueous layer was extracted a final time with chloroform. 0.1 volume of 3 M NaOAc (pH 5.2) and 3 volume of ethanol were added. The DNA was finally spun down (3000 g), and resuspended and stored in water at -20 °C.

#### Nucleosome core particle (NCP) reconstitution and characterisation

Reconstitution was performed following literature protocol[6]. Reconstituted products were analysed by non-denaturing PAGE followed by SYBR SAFE staining.

Large-Scale Reconstitution of NCP:

The concentration of Widom 601 145 bp DNA was adjusted with $H_2O$ and then added to the same volume of 4 M KCl to achieve a final salt concentration of 2 M and equimolar concentration to octamer. The DNA and octamer were mixed in the ratio 1.15:1.0 and dialysed at 4 °C against TCS reconstitution buffer (2 M KCl, 10 mM Tris, 1 mM EDTA, pH 7.4) for 1.5 h, and then against TCS buffer with 0.85 M KCl (1.5 h), 0.65 M KCl (3 h) and finally PBS (overnight). The sample was centrifuged at 4 °C for 15 min (10000 g) to separate soluble NCP precipitates. An aliquot was obtained for SDS-PAGE analysis to ensure that the core histones were still present in equimolar ratios. Non-denaturing PAGE analysis showed a homogenous population of NCP and very little free DNA.

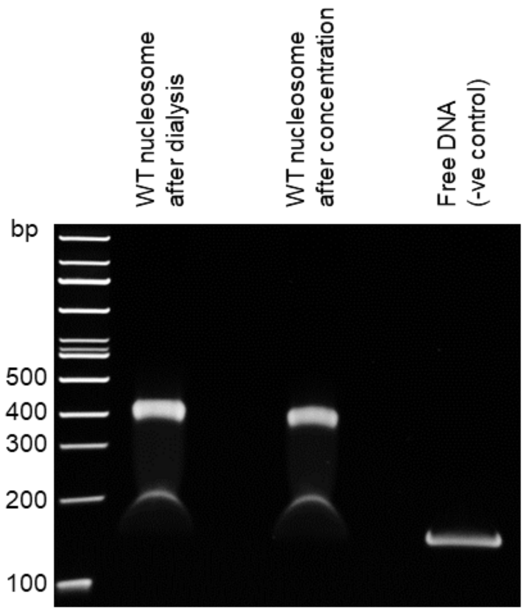

Non-denaturing PAGE

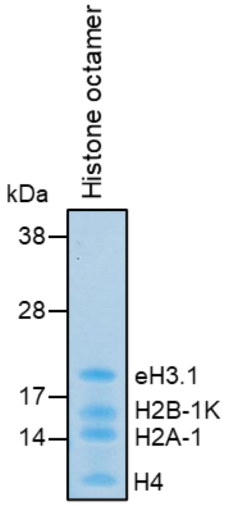

SDS-PAGE

To investigate effects of nucleosomes on rodent brain tissue, the NCP was concentrated using an Amicon centrifugal filter (10 kDa molecular weight cut-off, 4 ˚C,10000 g). Non-denaturing PAGE analysis showed that the NCP remained intact.

Non-denaturing (native) PAGE protocol: Non-denaturing PAGE analysis was performed using 6% Novex TBE Gel (Thermofisher) and a 100-bp DNA ladder (Thermofisher). The gel was prerun in 0.5x TBE at 150 V for 90 min at 4 °C. Then ~20 pmol of nucleosome in 4 µL was supplemented with 1 µL of 30% sucrose as the loading buffer before the sample was added into the well. The gel was run at 150 V for at least 60 min at 4 °C and then stained with SYBR Safe (Invitrogen, S-33102) for 15 min. The gel was rinsed with $H_2O$ for at least another 15 min and then visualised using a Bio-Rad Gel Doc XR+ instrument.

Coomassie stain on native PAGE: After imaging, the 6% TBE native gel was stained and fixed with formic acid and methanol overnight (5 g Coomassie G250 in 1 L of MeOH, $H_2O$ and AcOH (4.5:4.5:1)) before destaining several times with a solution of EtOH, AcOH and $H_2O$ (3:1:6).

### *Histone Dha formation*

Lyophilised histone (7 mg) was dissolved in denaturing phosphate buffer (100 mM $NaP_i$, pH 8, 3 M Gdn·HCl, 500 µL). DTT was added (30 mg) and the reaction mixture was shaken for 30 min at RT (500 rpm) to reduce disulfide bonds, before desalting into 1 mL of the same buffer (PD Minitrap G25, GE Healthcare). The resulting protein concentration was determined spectrophotometrically. This was accomplished here and for all proteins via absorption at 280 nm (noted as 'Nanodrop'; Implen NanoPhotometer® N60) based on protein-specific extinction coefficients calculated with the Expasy ProtParam tool for given amino acid sequences. This was then immediately followed by the addition of DBHDA (60 equiv from a freshly prepared 0.5 M DMSO stock) and subsequent shaking (500 rpm) at 25 °C for 1 h, then 37 °C for at least 2 h. The protein was desalted as before to remove the excess DBHDA and exchange into the desired buffer. Protein yield and concentration were determined by its UV absorbance using the Nanodrop and conversion was determined by LC-MS analysis.

Representative LC-MS analysis (ion series and magnification of the deconvoluted spectrum):

Human histone eH3.1-Dha4
Calculated mass: 17686 Da
Observed mass: 17686 Da

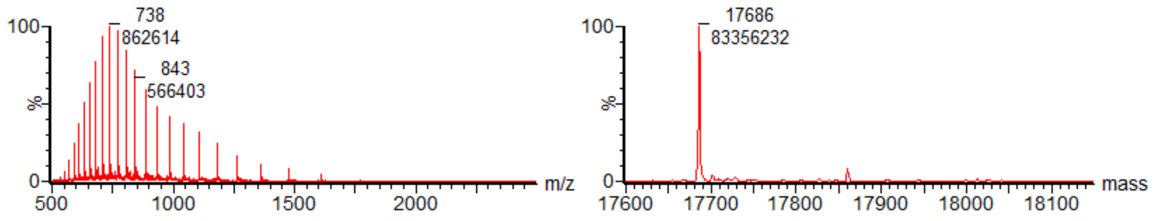

Human histone eH3.1-Dha27
Calculated mass: 17686 Da
Observed mass: 17687 Da

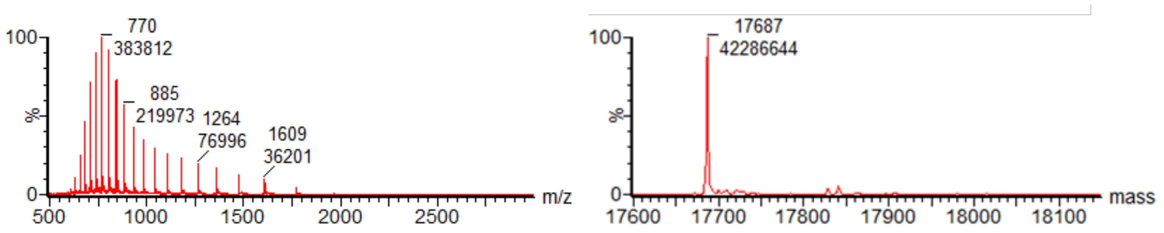

Human histone eH3.1-Dha56

Calculated mass: 17686 Da

Observed mass: 17687 Da

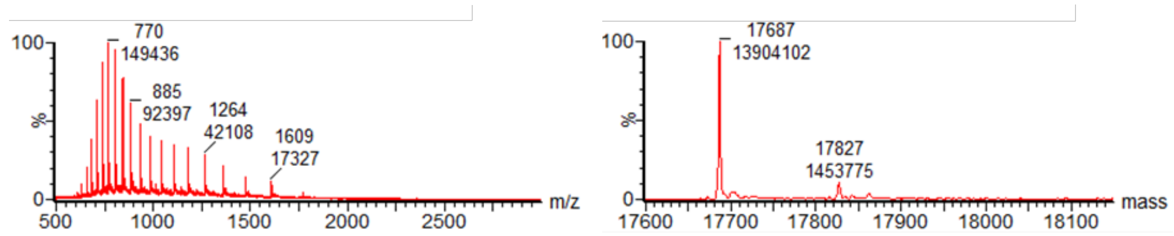

### *Light-driven modification of Dha-containing histone H3*

#### General protein reaction protocol for $^{19}F$-modification

All protein solutions and solvents were degassed for at least 8 h in a glovebox (<10 ppm $O_2$). All solid reagents were weighed and then transferred into the glovebox where stock solutions were prepared prior to protein reactions. Protein reactions were carried out in clear glass vials (ThermoFisher Scientific 9 mm clear glass screw thread vial with 300 µL fused insert, for reactions ≤50 µL, ThermoFisher Scientific 2.0 mL 9 mm clear glass screw thread vial, flat bottom, for reactions ≤100 µL) with gas-tight lids. Standard reactions were prepared by diluting the Dha-containing protein solution in the buffer of choice to reach the desired protein concentration, followed by sequential addition of photocatalyst, sulfone reagents and iron additives in the glovebox. The reactions were then mixed thoroughly with a pipette, capped and moved out of the glovebox for irradiation. A variable light intensity (10 – 50 W) photobox was used with blue LEDs arranged in series to allow up to 7 reactions at a time. Cooling fan is attached to the photobox for temperature control especially for reactions requiring >20 min. After irradiation, an aliquot of the crude reaction mixture was obtained and diluted into 15 mM β-mercaptoethanol in 0.1% formic acid for LC-MS analysis. Unless stated otherwise, conversions reported are to the desired single addition fluoroalkylated product. Low level of double addition arises from quenching of the α-carbon radical intermediate with a second $^{19}F$-containing radical. [7]

### Formation of human histone eH3.1-hAla4

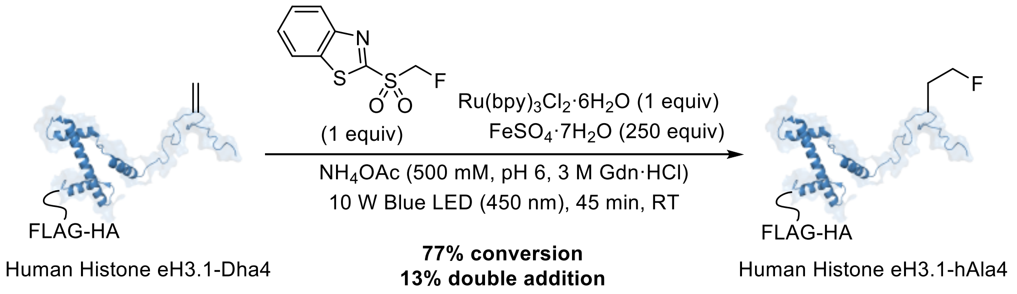

In the glovebox, degassed $NH_4OAc$ buffer (500 mM, pH 6, 3 M Gdn·HCl) was added to a glass HPLC vial containing human histone eH3.1-Dha4 (100 µg) such that the final concentration was 1 mg $mL^{-1}$ (57 µM) i.e. total reaction volume was 100 µL. $Ru(bpy)_3Cl_2·6H_2O$ (1 µL of a 5.7 mM stock prepared fresh in degassed $H_2O$, 1 equiv), mono-btSOOF radical precursor (1 µL of a 5.7 mM stock prepared fresh in degassed DMSO, 1 equiv) and $FeSO_4·7H_2O$ (5 µL of a 285 mM stock prepared fresh in degassed $H_2O$, 250 equiv) were added sequentially and the vial was sealed with a lid before transferring out of the glovebox and irradiating with blue LED light (10 W) for 45 min. A sample of the crude mixture was then analysed by LC-MS to determine reaction conversion.

ESI-MS analysis of human histone eH3.1:

Calculated hAla-modified mass: 17720 Da

Observed mass: 17719 Da (+16 Da is methionine oxidation, +32 Da is double addition)

Calculated Dha mass: 17686 Da

Observed mass: 17686 Da

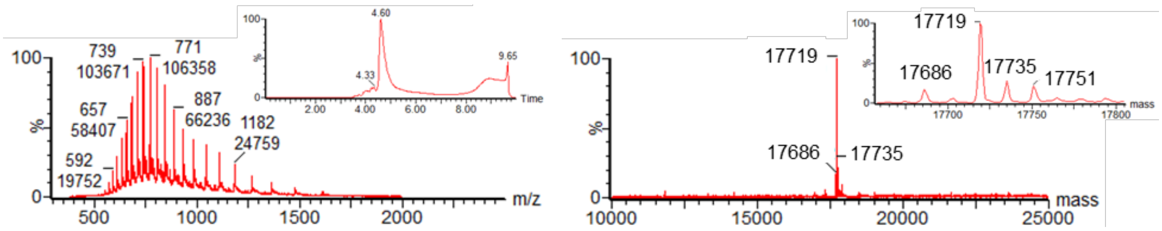

### Formation of human histone eH3.1-hAla56

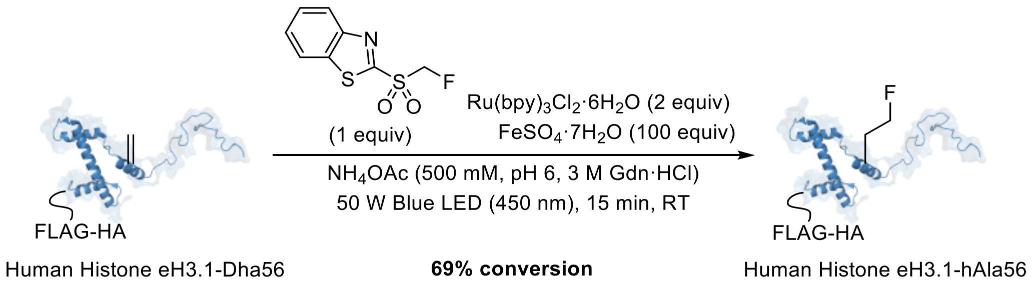

In the glovebox, degassed $NH_4OAc$ buffer (500 mM, pH 6, 3 M Gdn·HCl) was added to a glass HPLC vial containing human histone eH3.1-Dha56 (100 μg) such that the final concentration was 1 mg $mL^{-1}$ (57 μM) i.e. total reaction volume was 100 μL. $Ru(bpy)_3Cl_2{\cdot}6H_2O$ (1 μL of a 5.7 mM stock prepared fresh in degassed $H_2O$, 1 equiv), mono-btSOOF radical precursor (1 μL of a 5.7 mM stock prepared fresh in degassed DMSO, 1 equiv) and $FeSO_4{\cdot}7H_2O$ (2.5 μL of a 228 mM stock prepared fresh in degassed $H_2O$, 100 equiv) were added sequentially and the vial was sealed with a lid before transferring out of the glovebox and irradiating with blue LED light (50 W) for 15 min. A sample of the crude mixture was then analysed by LC-MS to determine reaction conversion.

ESI-MS analysis of human histone eH3.1:
Calculated hAla-modified mass: 17720 Da
Observed mass: 17720 Da (+16 Da is methionine oxidation, +32 Da is double addition)
Calculated Dha mass: 17686 Da
Observed mass: 17686 Da (+16 Da is methionine oxidation)

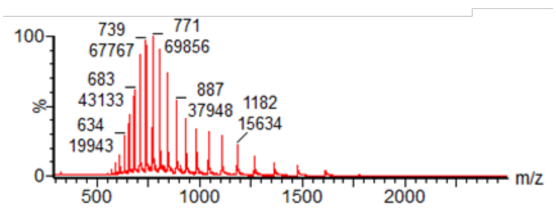

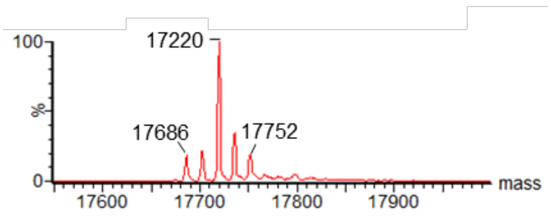

### Formation of human histone eH3.1-Lys(F)27

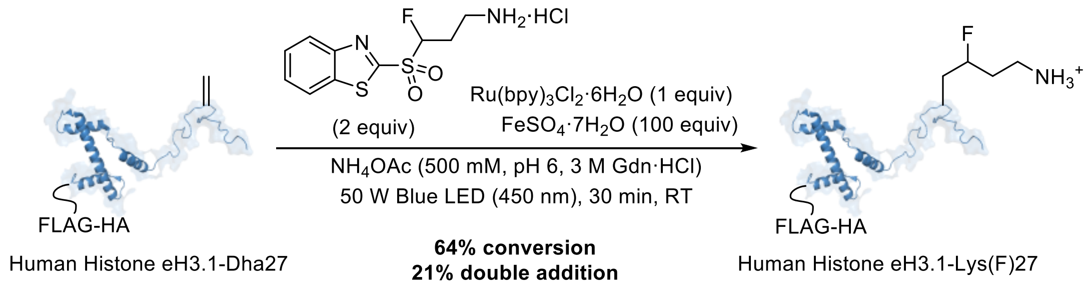

In the glovebox, degassed $NH_4OAc$ buffer (500 mM, pH 6, 3 M Gdn·HCl) was added to a glass HPLC vial containing human histone eH3.1-Dha4 (100 µg) such that the final concentration was 1 mg $mL^{-1}$ (57 µM) i.e. total reaction volume was 100 µL. $Ru(bpy)_3Cl_2 \cdot 6H_2O$ (1 µL of a 5.7 mM stock prepared fresh in degassed $H_2O$, 1 equiv), mono-btSOOF-based radical precursor (2 µL of a 5.7 mM stock prepared fresh in degassed DMSO, 1 equiv) and $FeSO_4 \cdot 7H_2O$ (2.5 µL of a 228 mM stock prepared fresh in degassed $H_2O$, 100 equiv) were added sequentially and the vial was sealed with a lid before transferring out of the glovebox and irradiating with blue LED light (50 W) for 30 min. A sample of the crude mixture was then analysed by LC-MS to determine reaction conversion.

ESI-MS analysis of human histone eH3.1:

Calculated Lys(F)-modified mass: 17763 Da

Observed mass: 17763 Da (+16 Da is methionine oxidation, +75 Da is double addition)

Calculated Dha mass: 17686 Da

Observed mass: 17686 Da

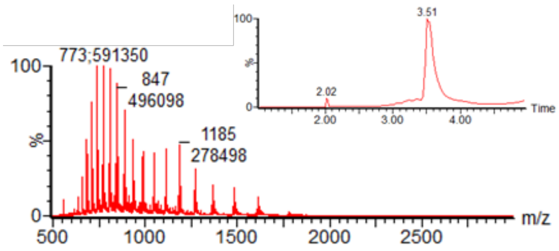

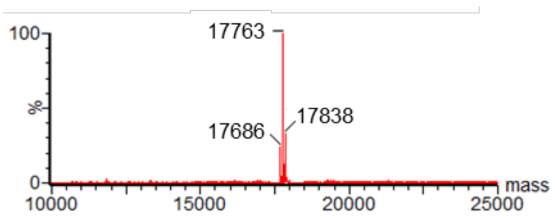

### Formation of human histone eH3.1-Lys(F)4

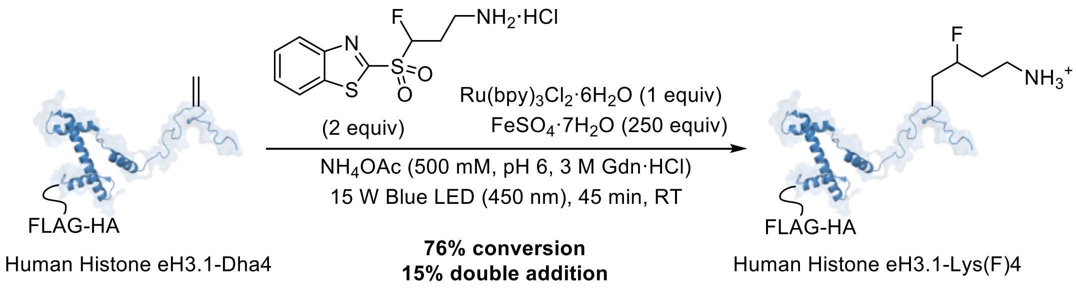

In the glovebox, degassed $NH_4OAc$ buffer (500 mM, pH 6, 3 M Gdn·HCl) was added to a glass HPLC vial containing human histone eH3.1-Dha4 (100 μg) such that the final concentration was 1 mg $mL^{-1}$ (57 μM) i.e. total reaction volume was 100 μL. $Ru(bpy)_3Cl_2·6H_2O$ (1 μL of a 5.7 mM stock prepared fresh in degassed $H_2O$, 1 equiv), mono-btSOOF-based radical precursor (2 μL of a 5.7 mM stock prepared fresh in degassed DMSO, 2 equiv) and $FeSO_4·7H_2O$ (7 μL of a 200 mM stock prepared fresh in degassed $H_2O$, 250 equiv) were added sequentially and the vial was sealed with a lid before transferring out of the glovebox and irradiating with blue LED light (15 W) for 45 min. A sample of the crude mixture was then analysed by LC-MS to determine reaction conversion.

ESI-MS analysis of human histone eH3.1:

Calculated Lys(F)-modified mass: 17763 Da

Observed mass: 17763 Da (+16 Da is methionine oxidation, +75 Da is double addition)

Calculated Dha mass: 17686 Da

Observed mass: 17686 Da

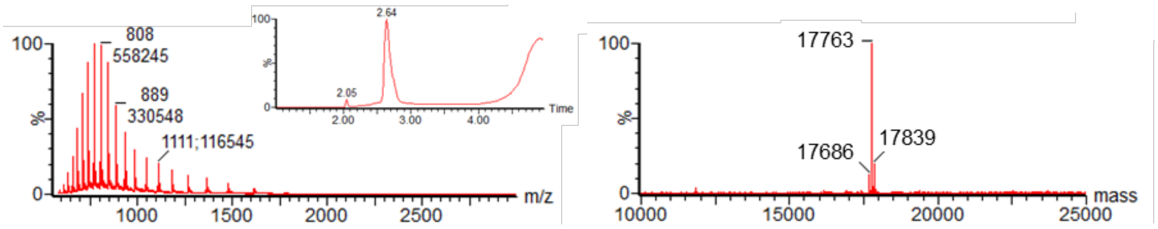

### *Small-molecule radiochemistry*

#### Manual radiosynthesis of [$^{18}$F]btSOOCH$_2$F using an Advion Nanotek® radiosynthesizer

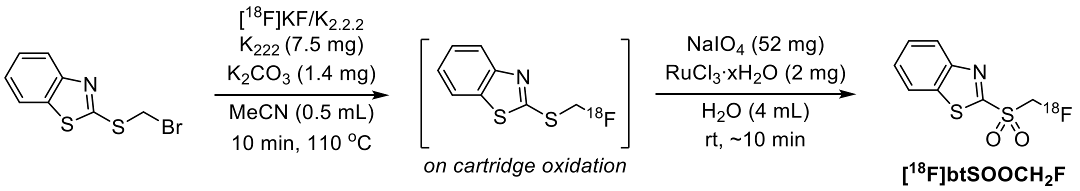

Following protocol established in previous work,[7] a solution of the brominated precursor (0.04 mmol in 0.5 mL dry MeCN) was added to a V-vial containing a stir bar and dry [$^{18}$F]KF/K$_{2.2.2}$. and the solution was left to stir at 110 °C for 10 min. The crude reaction containing the $^{18}$F-labelled compound was then allowed to cool prior to dilution with 4 mL of $H_2O$. The mixture was then filtered through a C18 plus cartridge (pre-conditioned with EtOH (10 mL) and $H_2O$ (10 mL)). A solution containing $NaIO_4$ (52 mg, 0.24 mmol) and $RuCl_3 \cdot xH_2O$ (2 mg, 10 mol%) in $H_2O$ (4 mL) was passed through the C18 plus cartridge, passing 1 mL of oxidation solution every 1 min. After complete addition, the oxidation was left for 5 min at room temperature. Crude [$^{18}$F]btSOOCH$_2$F was then eluted from the cartridge with 1.2 mL of MeCN prior to semi-prep HPLC purification (flow rate = 4 mL min$^{-1}$, Phenomenex Synergi Hydro-RP 4 µm 80 Å LC column (250 x 10 mm)) with 53% MeCN in 25 mM ammonium formate buffer (isocratic). The radio-HPLC peak corresponding to [$^{18}$F]btSOOCH$_2$F was collected in a vial containing 20 mL of water. This solution was then passed over a C18 plus cartridge (pre-conditioned with EtOH (10 mL) and $H_2O$ (10 mL)). The reagent was then eluted from the cartridge into a V-vial with $Et_2O$ (~1.2 mL total volume). This solution was then concentrated to dryness under a flow of $N_2$ at rt. Protein solution containing the photocatalyst and Fe(II) in degassed, buffered conditions was subsequently added under $N_2$.

### Manual radiosynthesis of [$^{18}$F]btSOOLys using an Advion Nanotek® radiosynthesizer

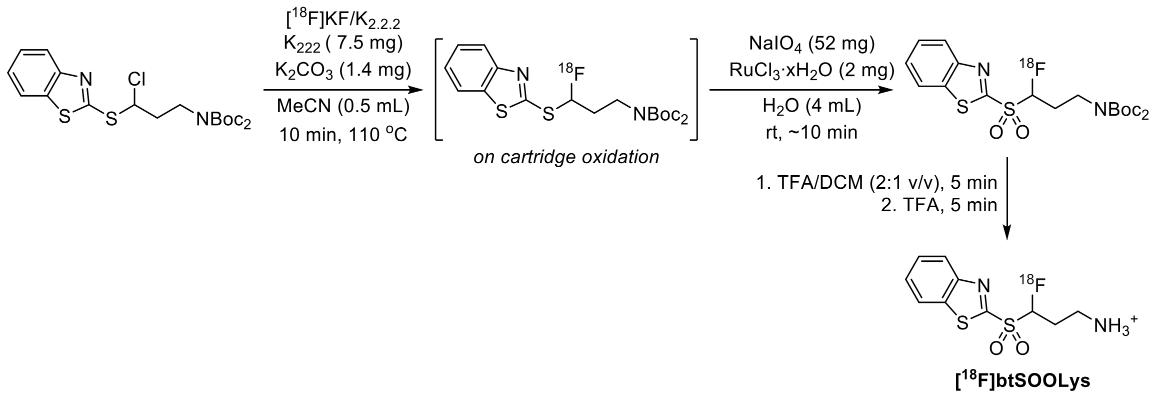

A solution of the chlorinated precursor (0.04 mmol in 0.5 mL MeCN) was added to a V-vial containing a stir bar and dry [$^{18}$F]KF/$K_{2.2.2}$ and the solution was left to stir at 110 °C for 10 min. The crude reaction containing the $^{18}$F-labelled compound was then allowed to cool prior to dilution with 4 mL of $H_2O$. The mixture was then filtered through a C18 plus cartridge (pre-conditioned with EtOH (10 mL) and $H_2O$ (10 mL)). A solution containing $NaIO_4$ (52 mg, 0.24 mmol) and $RuCl_3{\cdot}xH_2O$ (2 mg, 10 mol%) in $H_2O$ (4 mL) was passed through the C18 plus cartridge, passing 1 mL of oxidation solution every 1 min. After complete addition, the oxidation was left for 5 min at room temperature. The crude labelled sulfone reagent was then eluted from the cartridge with 1.2 mL of MeCN prior to semi-prep HPLC purification (flow rate = 4 mL min$^{-1}$, Phenomenex Synergi Hydro-RP 4 µm 80 Å LC column (250 x 10 mm)) under isocratic conditions with 75% MeCN in 25 mM ammonium formate buffer. The peak corresponding to the benzothiazole sulfone reagent was collected in a vial containing 20 mL of water. This solution was then passed over a C18 plus cartridge (pre-conditioned with EtOH (10 mL) and $H_2O$ (10 mL)). The reagent was then eluted from the cartridge into a V-vial with $Et_2O$ (~2 mL total volume). This solution was then concentrated to dryness under a flow of $N_2$ at rt. To the dried diboc protected sulfone reagent was added TFA and DCM (2:1 v/v, 0.4 mL) and the reaction mixture was shaken for 5 min at rt. Under this condition, 30% of the sulfone reagent undergo complete deprotection while the remaining 70% still contained mono-boc protecting group. The addition of another 0.4 mL of pure TFA followed by a 5 min vortex ensured complete deprotection. The solvents were removed under a flow of $N_2$ at 50 °C. Protein solution containing the photocatalyst and Fe(II) in degassed, buffered conditions was subsequently added under $N_2$.

[$^{18}$F]btSOOCH(F)–$(CH_2)_2NBoc_2$ radio-HPLC trace (Method A):

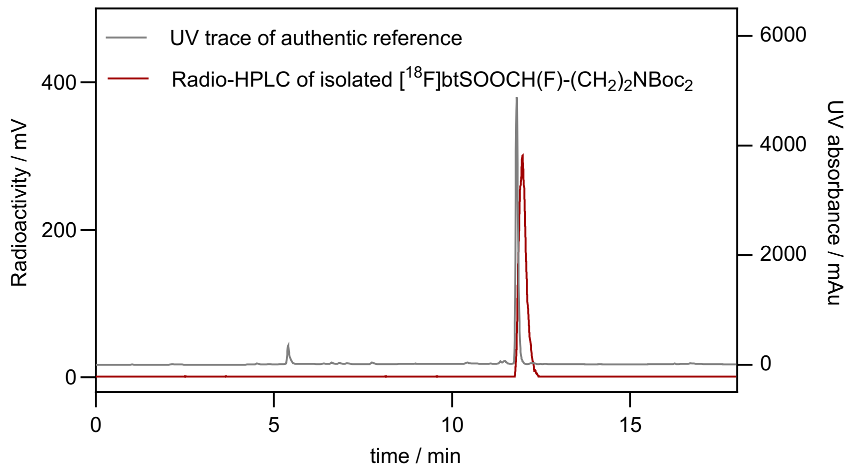

[$^{18}$F]btSOOCH(F)–$(CH_2)_2NH_2$·HCl radio-HPLC trace (Method B):

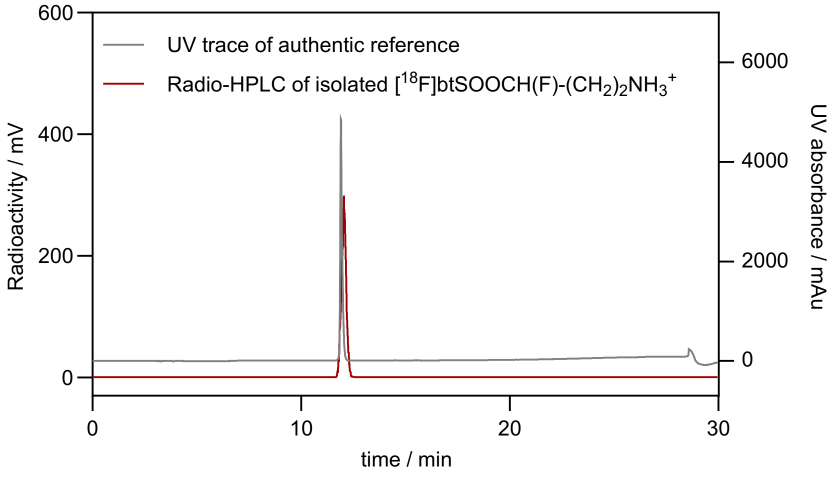

### Automatic radiosynthesis of [$^{18}$F]btSOOCH$_2$F using a Trasis AllinOne radiosynthesiser

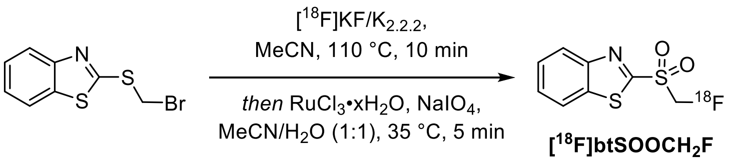

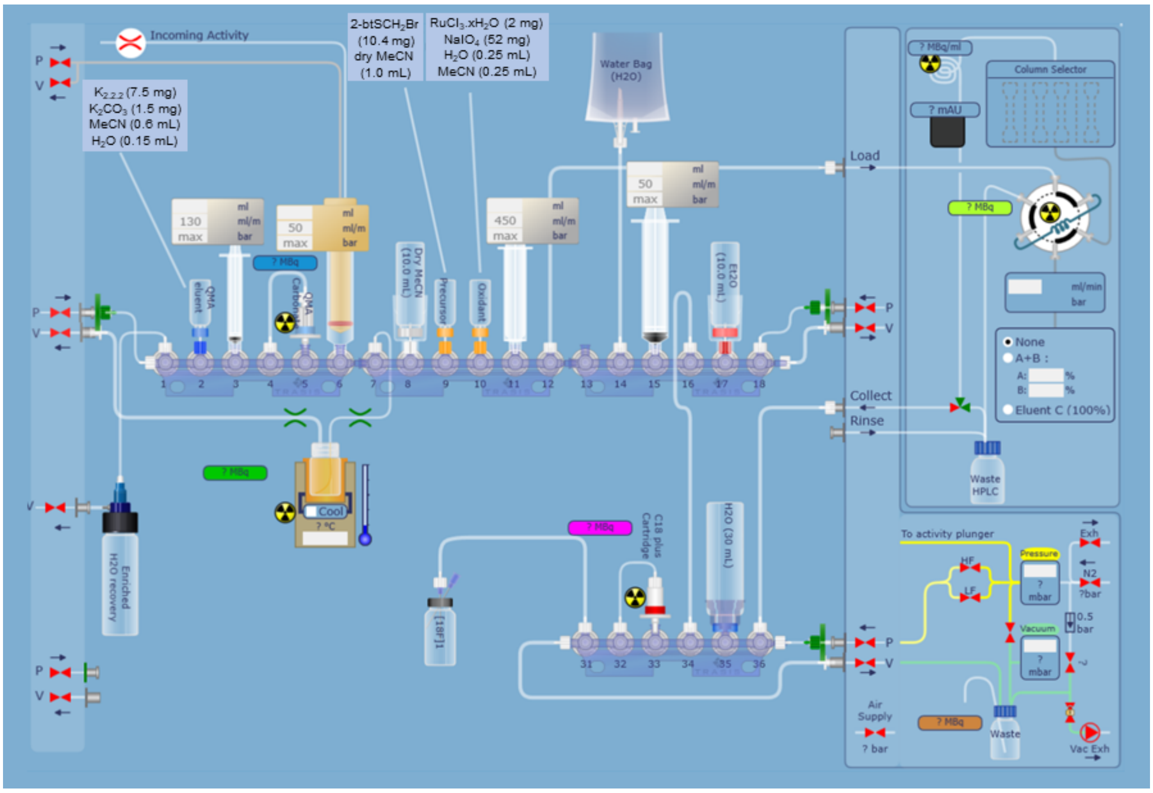

**Figure 1:** Cassette layout for the automated synthesis of [$^{18}$F]btSOOCH$_2$F on the Trasis AllInOne platform.

Similar to previous work,[7] the automated radiosynthesis of [$^{18}$F]btSOOCH$_2$F was carried out with a Trasis AllInOne synthesizer / synthesis unit using a cassette set-up as shown in Figure 1. Briefly, the vial in position 2 charged with $K_2CO_3$ (1.5 mg), $K_{2.2.2}$ (7.5 mg), MeCN (0.6 mL) and $H_2O$ (0.15 mL). The vial at position 9 was charged with the brominated precursor (0.04 mmol) dissolved in dry MeCN (1.0 mL). The vial in position 10 was charged with $RuCl_3{\cdot}xH_2O$ (2 mg), $NaIO_4$ (52 mg), $H_2O$ (0.25 mL) and MeCN (0.25 mL). The vials in position 8 and 17 were filled with dry MeCN and dry $Et_2O$ (approximately 10 mL each) respectively. A water bag was placed in position 14. The HPLC collection vial was filled with $H_2O$ (30 mL) and placed in position 35. A Waters Sep-Pak AccellPlus QMA Carbonate Plus Light cartridge was pre-conditioned with $H_2O$ prior to use and placed in position 5. A Waters Sep-Pak C18 Plus Short cartridge was activated with EtOH (10 mL) and then $H_2O$ (10 mL) prior to use and was placed in position 33.

[$^{18}$F]Fluoride in [$^{18}$O]water was received directly from the cyclotron into the Trasis AllInOne radiosynthesis unit. The [$^{18}$F]fluoride was separated from the water by trapping on the QMA cartridge, followed by elution with the solution from the vial in position 2 into the reactor. The [$^{18}$F]fluoride was then dried, and once this was complete, the $^{18}$F-fluorination of the brominated substrate (2-btSCH$_2$Br) proceeded in the same reactor at 110 °C for 10 min. The reactor was then cooled and the contents of the vial at position 10 were added to the same reactor. The oxidation reaction proceeded at 35 °C for 5 minutes. The crude reaction mixture was diluted with $H_2O$ and MeCN and transferred to the HPLC injection loop for semi-preparative reverse phase HPLC purification (MeCN:$H_2O$ = 55:45, flow rate = 4 mL min$^{-1}$) with a Phenomenex Synergi Hydro-RP 4 µm 80 Å LC column (250 x 10 mm). The desired purified product ($t_R$ = 10.5-11.0 min) was collected in the vial at position 35. This solution was diluted further with $H_2O$ for reformulation using a C18 Plus cartridge. [$^{18}$F]btSOOCH$_2$F was then eluted with $Et_2O$ (2.0 mL) into a V-vial and a syringe pump was used to remove residual $H_2O$ (~0.45 mL). $Et_2O$ was then removed under vacuum and a flow of $N_2$ to yield dried [$^{18}$F]btSOOCH$_2$F (total drying time ~5 min).

### Automatic radiosynthesis of [$^{18}$F]btSOOLys using a Trasis AllinOne radiosynthesiser

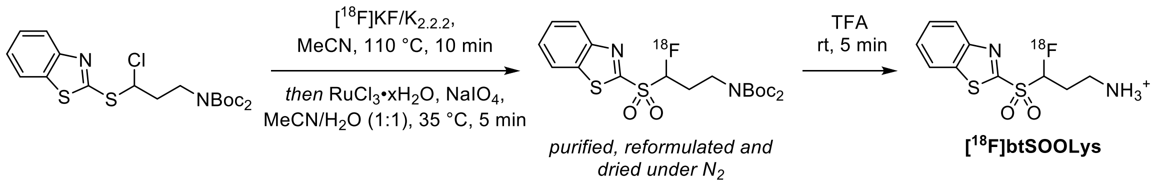

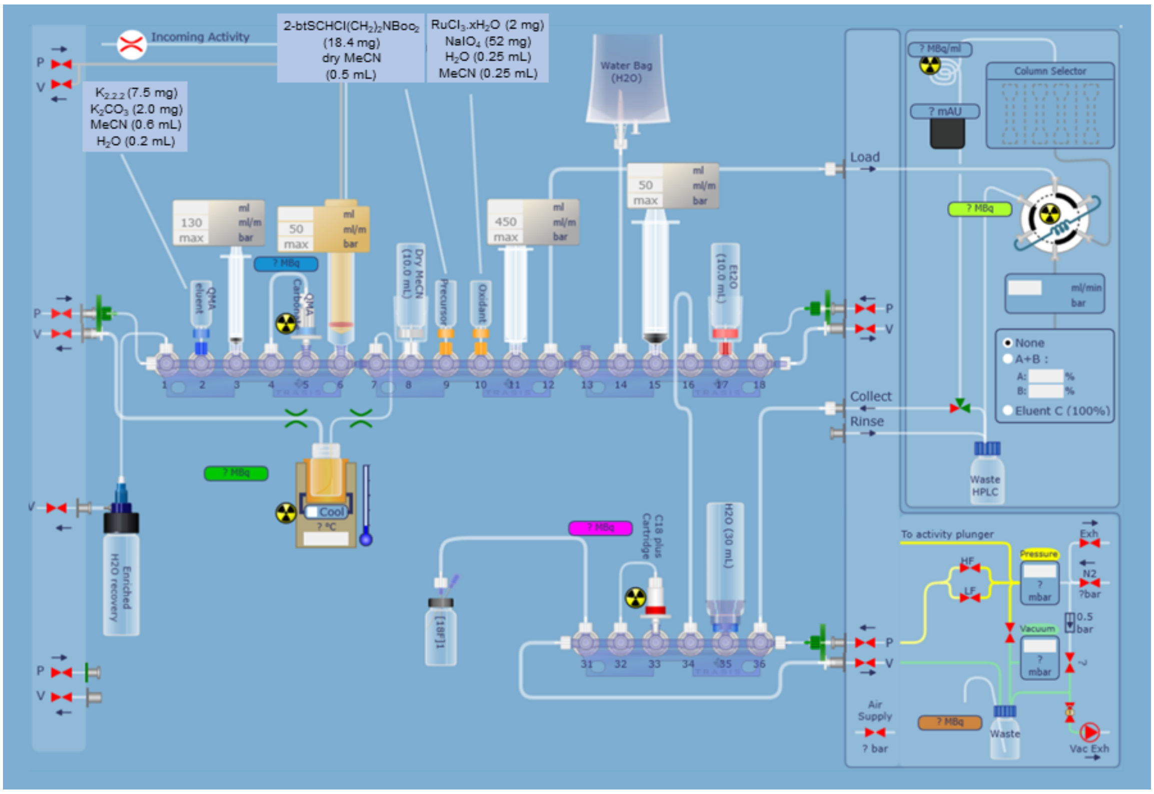

**Figure 2:** Cassette layout for the automated synthesis of [$^{18}$F]btSOOLys on the Trasis AllInOne platform.

The automated radiosynthesis of [$^{18}$F]btSOOLys was carried out in part with a Trasis AllInOne synthesizer / synthesis unit using a cassette set-up as shown in Figure 2 to yield the diboc-protected $^{18}$F-labelled sulfone product. Deprotection was performed through the manual addition of TFA. The vial in position 2 charged with $K_2CO_3$ (2.0 mg), $K_{2.2.2}$ (7.5 mg), MeCN (0.6 mL) and $H_2O$ (0.2 mL). The vial at position 9 was charged with the chlorinated precursor (0.04 mmol) dissolved in dry MeCN (0.5 mL). The vial in position 10 was charged with $RuCl_3{\cdot}xH_2O$ (2 mg), $NaIO_4$ (52 mg), $H_2O$ (0.25 mL) and MeCN (0.25 mL). The vials in position 8 and 17 were filled with dry MeCN and dry $Et_2O$ (approximately 10 mL each) respectively. A water bag was placed in position 14. The HPLC collection vial was filled with $H_2O$ (30 mL) and placed in position 35. A Waters Sep-Pak AccellPlus QMA Carbonate Plus Light Cartridge was pre-conditioned with $H_2O$ (10 mL) prior to use and placed in position 5. A Waters Sep-Pak C18 Plus Short cartridge was activated with EtOH (10 mL) and then $H_2O$ (10 mL) prior to use and was placed in position 33.

[$^{18}$F]Fluoride in [$^{18}$O]water was received directly from the cyclotron into the Trasis AllInOne radiosynthesis unit. The [$^{18}$F]fluoride was separated from the water by trapping on to a QMA cartridge (Waters Sep-Pak AccellPlus QMA Carbonate Plus Light Cartridge preconditioned with 10 mL of $H_2O$) followed by elution with the solution from the vial in position 2 into the reactor. The [$^{18}$F]fluoride was then dried, and once this was complete, the $^{18}$F-fluorination of the chlorinated substrate (2-btSCHCl($CH_2$)$_2$NBoc$_2$) proceeded in the same reactor at 110 °C for 10 min. The reactor was then cooled and the contents of the vial at position 10 were added to the same reactor. The oxidation reaction proceeded at 35 °C for 5 minutes. The crude reaction mixture was diluted with $H_2O$ and MeCN and transferred to the HPLC injection loop for semi-preparative reverse phase HPLC purification (MeCN:$H_2O$ = 3:1, flow rate = 4 mL min$^{-1}$) with a Phenomenex Synergi Hydro-RP 4 µm 80 Å LC column (250 x 10 mm). The desired purified product ($t_R$ = 15.0-15.5 min) was collected in the vial at position 35. This solution was diluted further with $H_2O$ for reformulation using a C18 Plus cartridge. The diboc-protected, $^{18}$F-labelled product was then eluted with $Et_2O$ (2.0 mL) into a V-vial and a syringe pump was used to remove residual $H_2O$ (~0.45 mL). $Et_2O$ was then removed under vacuum and a flow of $N_2$ to yield dried, diboc-protected 2-[$^{18}$F]btSCHF($CH_2$)$_2$NBoc$_2$ (total drying time ~5 min). Neat TFA (0.4 mL) was added using a syringe and the V-vial was shaken for ~5 s. Deprotection was left to proceed at rt for 5 min before the solvent was removed under vacuum and a flow of $N_2$ to yield dried [$^{18}$F]btSOOLys. Addition of $Et_2O$ (0.5 mL) and heating (50 °C for 2-3 min) were required to facilitate efficient removal of TFA (total drying time ~10-15 min).

[$^{18}$F]btSOOCH(F)–$(CH_2)_2NH_2$·HCl radio-HPLC trace (Method B):

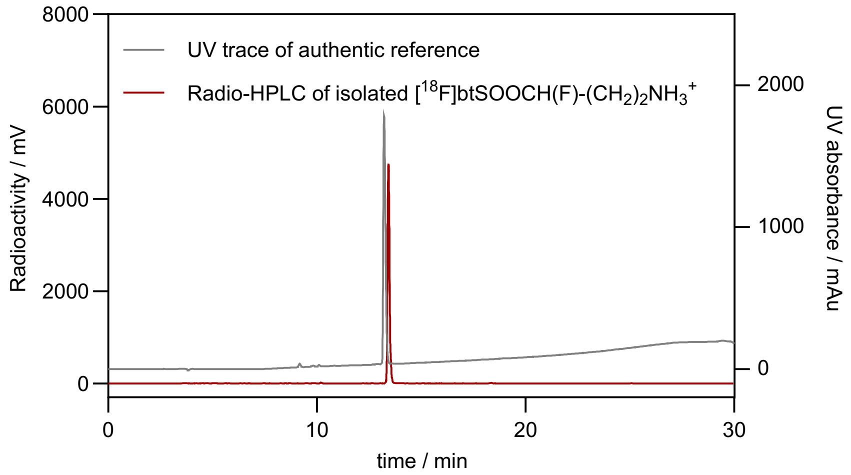

### *Protein radiochemistry*

#### General protein reaction protocol for $^{18}F$-modification

Protein solutions and solvents were degassed either in a glovebox (<10 ppm $O_2$) for at least 8 hours or by purging with nitrogen for at least 15 min. For maintaining anaerobic conditions with a glovebox, the photocatalyst and iron additive were weighed and then transferred into the glovebox where stock solutions were prepared with degassed solvents. Protein reactions (≤500 μL) were prepared in 3.0 mL V-vials (Wheaton clear glass V-vials) by diluting the Dha-containing protein solution in the buffer of choice to reach the desired protein concentration, followed by sequential addition of photocatalyst, iron additives and DMSO (<10% of total reaction volume) in the glovebox. The reactions were then mixed thoroughly with a pipette, capped with 18 mm PTFE/silicone septa inserted in the screw caps and moved out of the glovebox. The 3.0 mL V-vial (sealed with a PTFE/silicone septum) containing the dried and purified sulfone reagent was purged with nitrogen for at least 1 min. A nitrogen balloon was inserted into the vial with the protein solution. Under nitrogen atmosphere, the protein solution was transferred into the V-vial containing the sulfone reagent and the vial was gently shaken before irradiation. For cases where a glovebox is not readily accessible, anaerobic conditions were achieving by purging the protein reactions (≤500 μL) in 7.0 mL V-vials with a nitrogen or argon balloon after mixing with other reaction reagents (pre-dissolved in degassed $H_2O$ or DMSO). Again, the now degassed protein reactions were transferred into vials containing the sulfone reagent under nitrogen and then the protein reactions were shaken before irradiation. A variable light intensity (10 – 50 W) photobox was used with blue LEDs arranged in series to allow up to 3 reactions (in 3.0 mL Wheaton V-vials) at a time. Cooling fan is attached to the photobox for temperature control especially for reactions requiring >20 min. After irradiation, an aliquot of the crude reaction mixture was obtained and diluted into $H_2O$ for radio-HPLC analysis to confirm $^{18}F$-labelling and calculate RCY. Before any animal studies, the crude mixture was purified following incubation with EDTA and then using PD MiniTrap G-25 (GE Healthcare) desalting columns (protein recovery ~50-60%). Aliquots can also be obtained immediately after irradiation for LC-MS analysis after isotopic decay.

### Formation of human histone eH3.1-[$^{18}$F]hAla4

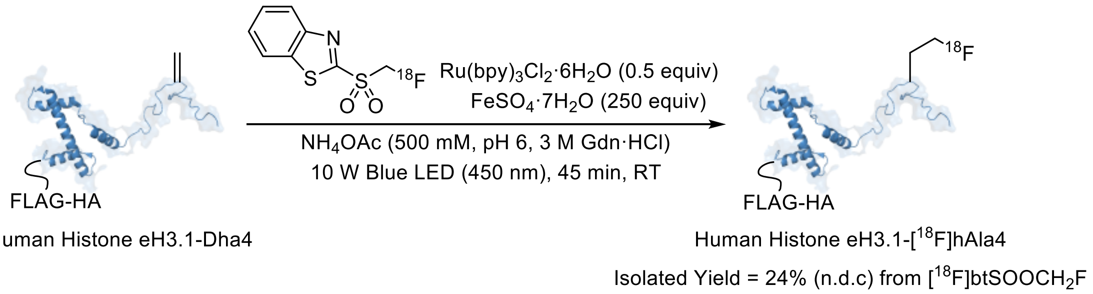

In the glovebox, degassed $NH_4OAc$ buffer (500 mM, pH 6, 3 M Gdn·HCl) was added to a glass V-vial containing human histone eH3.1-Dha4 such that the final concentration was 125 µM in a total reaction volume of 500 µL. $Ru(bpy)_3Cl_2·6H_2O$ (1 µL of a 31 mM stock prepared fresh in degassed $H_2O$, 0.5 equiv), $FeSO_4·7H_2O$ (30 µL of a 520 mM stock prepared fresh in degassed $H_2O$, 250 equiv) and DMSO (<10% v/v) were added sequentially and the V-vial was sealed before transferring out of the glovebox. Under nitrogen, the protein solution was transferred into a nitrogen-filled V-vial containing the purified [$^{18}$F]btSOOCH$_2$F, shaken and then irradiated with blue LED light (10 W) for 45 min. A sample of the crude mixture was then analysed by radio-HPLC to give an RCY value of 67 ± 5% ($n$ = 2, n.d.c.).

At the end of the reaction, 500 equiv. of EDTA (from a 0.5 M stock) was added and the crude reaction mixture was diluted up to a final volume of 500 µL before loading onto a PD-10 MiniTrap desalting column (pre-equilibrated with PBS). The first 200 µL elution volume was discarded. $^{18}$F-labelled histone was collected in the next 400 µL of buffered solution. An aliquot was taken to check the radio- and UV purity. The generation of purified human histone eH3.1-[$^{18}$F]hAla4 yielded an activity value of 85 MBq in 400 µL of the eluting buffer, starting from 360 MBq of [$^{18}$F]btSOOCH$_2$F. Hence, the activity yield of human histone eH3.1-[$^{18}$F]hAla4 was 24% (n.d.c) starting from dried [$^{18}$F]btSOOCH$_2$F. The radio- and UV purity were >99% and >97% respectively.

Crude HPLC analysis (Method B):

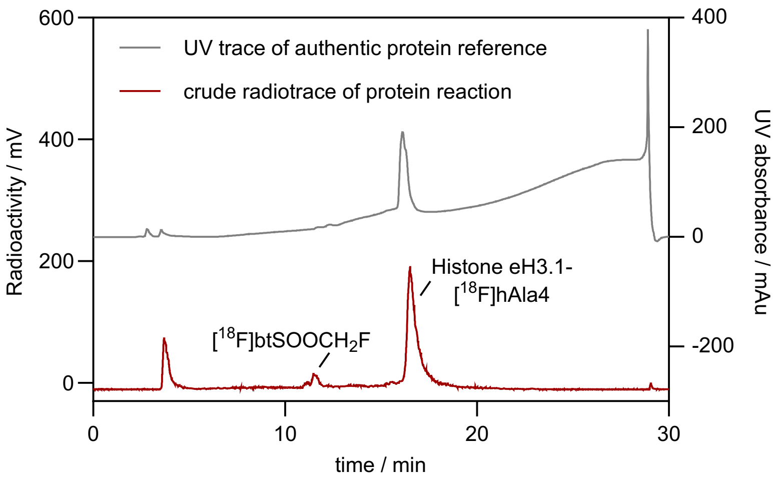

HPLC analysis of purified histone eH3.1-[18F]hAla4 (Method B):

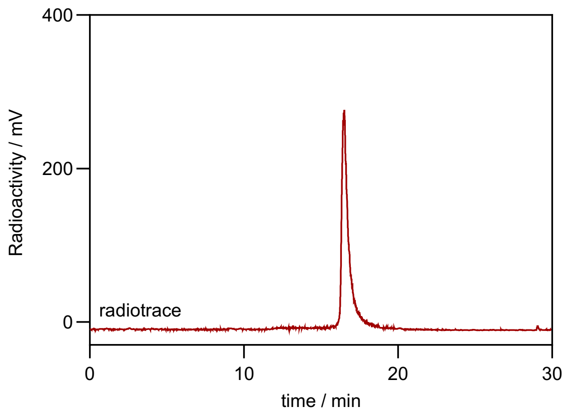

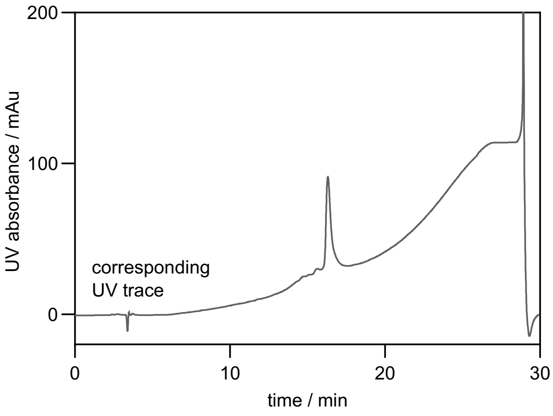

LC-MS analysis post-protein radiolabelling:

ESI-MS analysis of human histone eH3.1:

Calculated Dha mass: 17686 Da

Observed mass: 17686 Da (+16 Da is methionine oxidation)

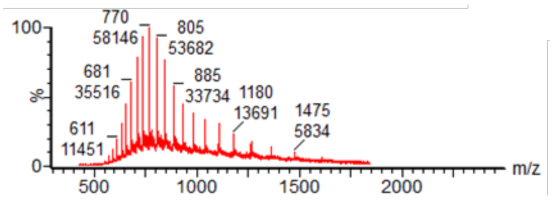

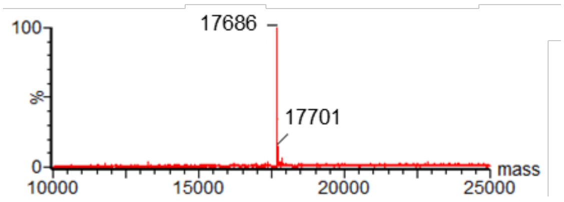

### Formation of human histone eH3.1-[$^{18}$F]hAla56

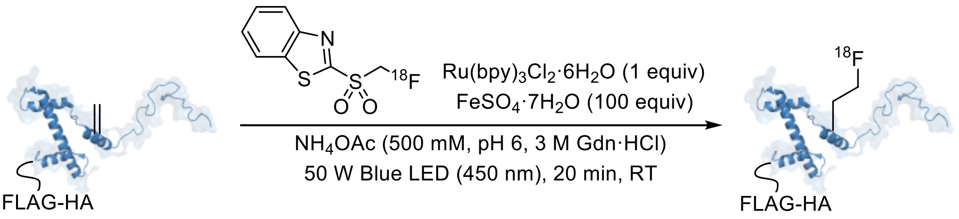

Human Histone eH3.1-Dha56 — Human Histone eH3.1-[$^{18}$F]hAla56

Isolated Yield = 16% (n.d.c) from [$^{18}$F]btSOOC$H_2$F

In the glovebox, degassed $NH_4OAc$ buffer (500 mM, pH 6, 3 M Gdn·HCl) was added to a glass V-vial containing human histone eH3.1-Dha56 such that the final concentration was 125 µM in a total reaction volume of 450 µL. Ru(bpy)$_3$Cl$_2$·6H$_2$O (2.81 µL of a 20 mM stock prepared fresh in degassed $H_2O$, 1 equiv), $FeSO_4$·7H$_2$O (11.25 µL of a 500 mM stock prepared fresh in degassed $H_2O$, 100 equiv) and DMSO (<10% v/v) were added sequentially and the V-vial was sealed before transferring out of the glovebox. Under nitrogen, the protein solution was transferred into a nitrogen-filled V-vial containing the purified [$^{18}$F]btSOOC$H_2$F, shaken and then irradiated with blue LED light (50 W) for 20 min. A sample of the crude mixture was then analysed by radio-HPLC to give an RCY of 49% n.d.c.

At the end of the reaction, 500 equiv. of EDTA (from a 0.5 M stock) was added and the crude reaction mixture was diluted up to a final volume of 500 µL before loading onto a PD-10 MiniTrap desalting column (pre-equilibrated with PBS). The first 200 µL elution volume was discarded. $^{18}$F-labelled histone was collected in the next 400 µL of buffered solution. An aliquot was taken to check the radio- and UV purity. The generation of purified human histone eH3.1-[$^{18}$F]hAla56 yielded an activity value of 56 MBq in 400 µL of the eluting buffer, starting from 353 MBq of [$^{18}$F]btSOOC$H_2$F. Hence, the activity yield of human histone eH3.1-[$^{18}$F]hAla56 was 16% (n.d.c) starting from dried [$^{18}$F]btSOOC$H_2$F. The radio- and UV purity were >99% and >98% respectively.

Crude HPLC analysis (Method B):

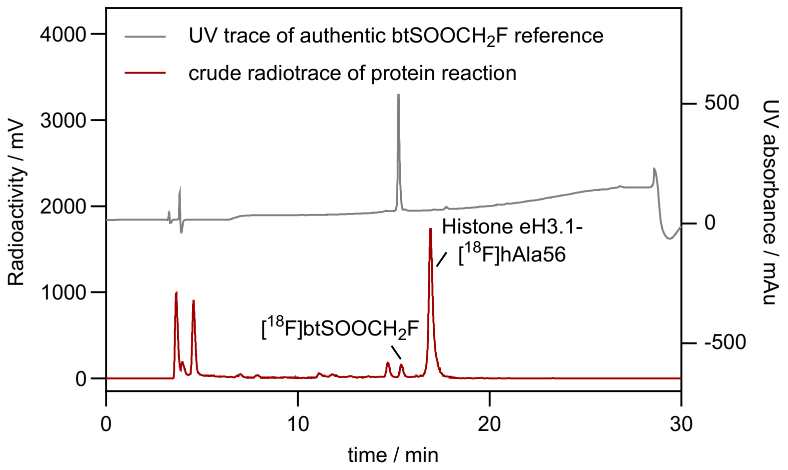

HPLC analysis of purified histone eH3.1-[$^{18}$F]hAla56 (Method B):

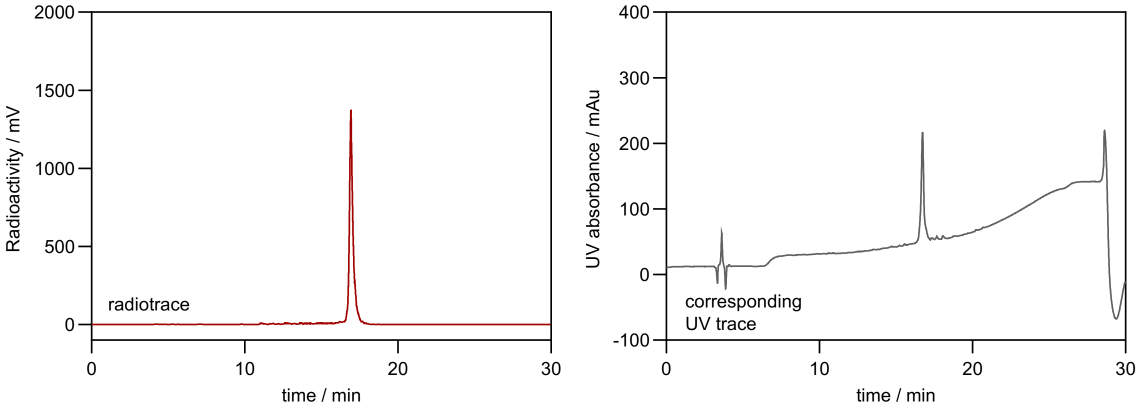

### Formation of human histone eH3.1-[$^{18}$F]Lys27

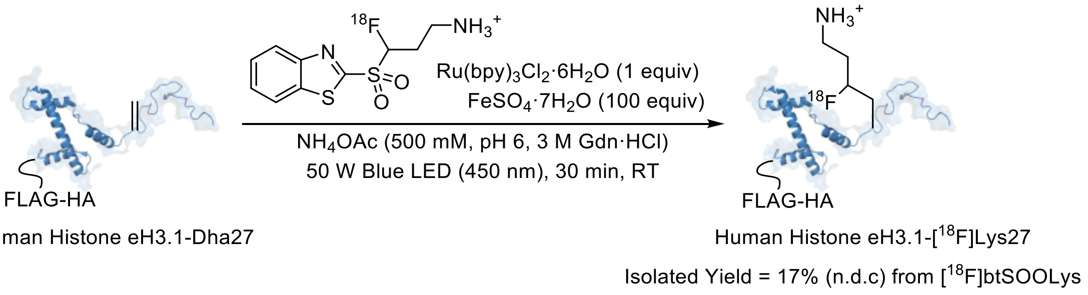

Histone eH3.1-[$^{18}$F]Lys27 was generated in the absence a glovebox whereby anaerobic condition was achieved by purging the protein solution for at least 15 min. $NH_4OAc$ buffer (500 mM, pH 6, 3 M Gdn·HCl) was added to a glass V-vial containing human histone eH3.1-Dha27 such that the final concentration was 200 µM in a total reaction volume of 500 µL. $Ru(bpy)_3Cl_2·6H_2O$ (4 µL of a 25 mM stock prepared fresh in degassed $H_2O$, 1 equiv), $FeSO_4·7H_2O$ (40 µL of a 250 mM stock prepared fresh in degassed $H_2O$, 100 equiv) and degassed DMSO (<10% v/v) were added sequentially and the V-vial was sealed. The protein reaction mixture was purged for a further 15 min. Under nitrogen, the protein solution was transferred into a nitrogen-filled V-vial containing the dried and purified [$^{18}$F]btSOOLys, shaken briefly and then irradiated with blue LED light (50 W) for 30 min.

At the end of the reaction, 500 equiv. of EDTA (from a 0.5 M stock) was added and the crude reaction mixture was loaded onto a PD-10 MiniTrap desalting column (pre-equilibrated with PBS). The first 200 µL elution volume was discarded. $^{18}$F-labelled histone was collected in the next 300 µL of buffered solution. An aliquot was taken to check the radio- and UV purity. The generation of purified human histone eH3.1-[$^{18}$F]Lys27 yielded an activity value of 36 MBq in 300 µL of the eluting buffer, starting from 218 MBq of [$^{18}$F]btSOOLys. Hence, the activity yield of human histone eH3.1-[$^{18}$F]Lys27 was 17% (n.d.c) starting from dried [$^{18}$F]btSOOLys. The radio- and UV purity were >98% and >98% respectively.

HPLC analysis of purified histone eH3.1-[$^{18}$F]Lys27 (Method B):

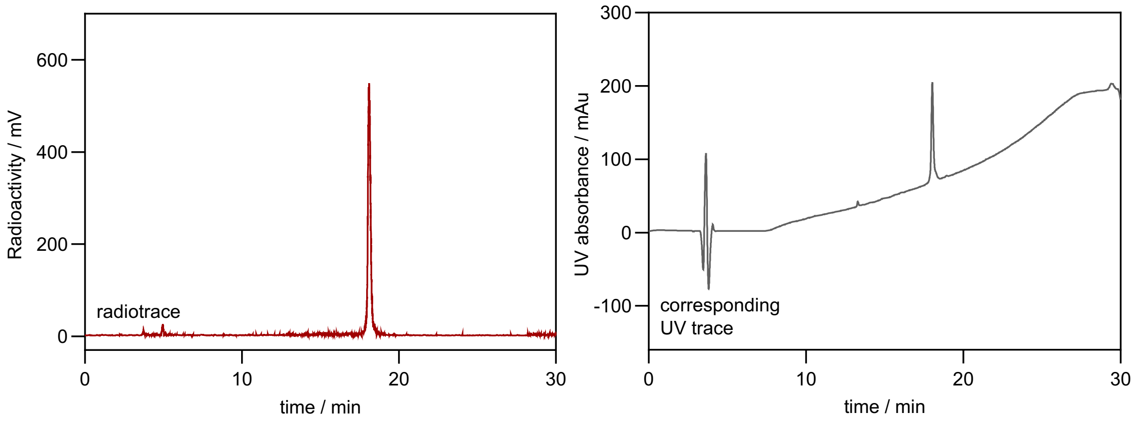

#### Formation of human histone eH3.1-[$^{18}$F]Lys4

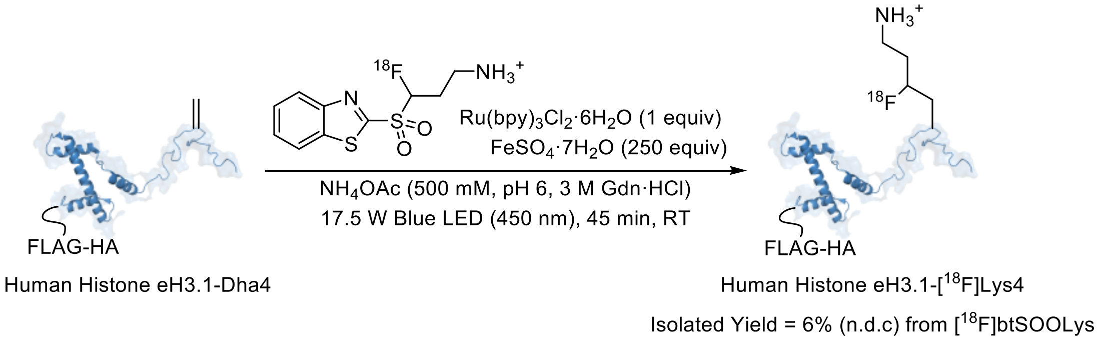

Histone eH3.1-[$^{18}$F]Lys4 was generated in the absence a glovebox whereby anaerobic condition was achieved by purging the protein solution for at least 15 min. $NH_4OAc$ buffer (500 mM, pH 6, 3 M Gdn·HCl) was added to a glass V-vial containing human histone eH3.1-Dha4 such that the final concentration was 200 μM in a total reaction volume of 500 μL. $Ru(bpy)_3Cl_2 \cdot 6H_2O$ (4 μL of a 25 mM stock prepared fresh in degassed $H_2O$, 1 equiv), $FeSO_4 \cdot 7H_2O$ (55.6 μL of a 450 mM stock prepared fresh in degassed $H_2O$, 250 equiv) and degassed DMSO (<10% v/v) were added sequentially and the V-vial was sealed. The protein reaction mixture was purged for a further 15 min. Under nitrogen, the protein solution was transferred into a nitrogen-filled V-vial containing the dried and purified [$^{18}$F]btSOOLys, shaken briefly and then irradiated with blue LED light (17.5 W) for 45 min.

At the end of the reaction, 500 equiv. of EDTA (from a 0.5 M stock) was added and the crude reaction mixture was loaded onto a PD-10 MiniTrap desalting column (pre-equilibrated with PBS). The first 200 μL elution volume was discarded. $^{18}$F-labelled histone was collected in the next 300 μL of buffered solution. An aliquot was taken to check the radio- and UV purity. The generation of purified human histone eH3.1-[$^{18}$F]Lys4 yielded an activity value of 26 MBq in 300 μL of the eluting buffer, starting from 418 MBq of [$^{18}$F]btSOOLys. Hence, the activity yield of human histone eH3.1-[$^{18}$F]Lys4 was 6% (n.d.c) starting from dried [$^{18}$F]btSOOLys. The radio- and UV purity were >99% and >95% respectively.

HPLC analysis of purified histone eH3.1-[$^{18}$F]Lys4 (Method B):

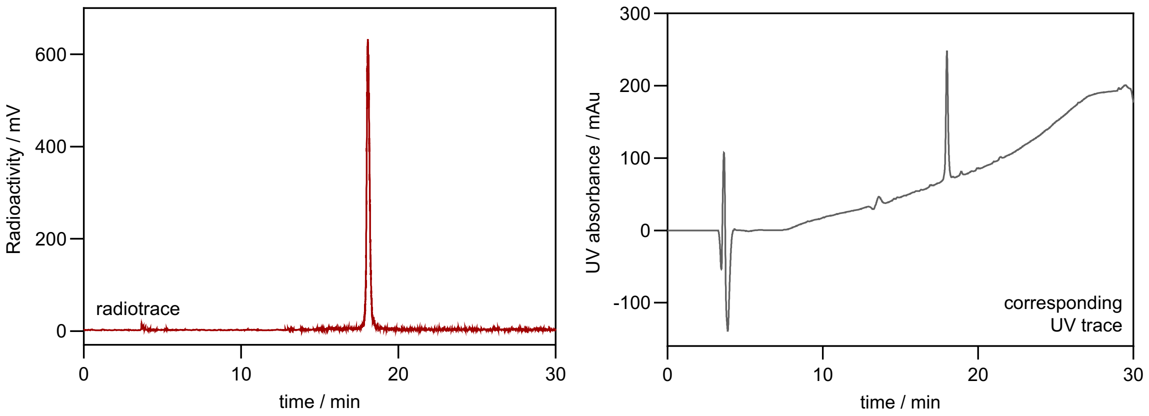

### *PET/CT studies of $^{18}$F-labelled histone H3*

#### Imaging of histones eH3.1-[$^{18}$F]hAla4 and 56

Female CD1 naïve mice (1 for [$^{18}$F]hAla4 and 2 for [$^{18}$F]hAla56) were anesthetized via inhalation of 3% isoflurane gas (0.5 L/min $O_2$) and placed on a warm heat mat in the prone position. The tail of the mouse was warmed with a heat lamp before the injection of [$^{18}$F]hAla-modified histone H3.1 via lateral tail vein ([$^{18}$F]hAla4: 10 MBq; [$^{18}$F]hAla56: 2-3 MBq) . Dynamic PET images (taken every 2.5 min for 2 h) were acquired after $^{18}$F-histone injections using a MILabs VECTor camera, equipped with an ultra-high resolution rat/mouse collimator (1.8mm), followed by a cone-beam CT scan (55 kV, 0.19 mA) for anatomical reference and attenuation correction. Mice were maintained under 2.5% isoflurane gas at 37 °C throughout the duration of image acquisition. PET images were reconstructed using U-SPECT-Rec3.22 software (MILabs, Utrecht, The Netherlands), applying a pixel-based algorithm, ordered subset expectation maximisation (OSEM-3D) with 128 subsets, 4 iterations and 0.8 mm voxel size for fluorine-18 (energy window settings 400–600 keV). Reconstructed PET and CT images were viewed and analysed using PMOD v3.38 (PMOD Technologies, Zurich, Switzerland). Nonlinear regression were performed using GraphPad Prism (GraphPad Software).

The mice were euthanized 2 h after $^{18}$F-histone injections. $^{18}$F-uptake in selected tissues were determined and reported as a percentage of the injected dose per gram of tissue (%ID/g) using a HIDEX automated gamma counter (HIDEX OY, Turku, Finland).

### Imaging of histones eH3.1-[$^{18}$F]Lys4 and 27

For *in vivo* studies with [$^{18}$F]Lys-modified histone H3 proteins, four mice were used for each of [$^{18}$F]Lys4 and [$^{18}$F]Lys27: two mice for *ex vivo* organ counting, tissue homogenisation plus cell fractionation; and then two mice for PET imaging studies plus *ex vivo* organ counting. The mice were imaged (two at a time) with the Molecubes β-cube (PET, MOLECUBES NV, Ghent, Belgium) immediately after being injected with 3.7-4.2 MBq of radiotracer into the tail vein for 120 min in list mode. The subjects were then CT imaged with the Molecubes X-cube immediately after PET imaging. The PET images were reconstructed into time frames 18 x 10s, 17 x 60s, 20 x 300s with OSEM 3D using 30 iterations and 1 subset. The PET/CT images were then analyzed with Carimas software (Turku PET Centre, Turku, Finland)

For both [$^{18}$F]Lys4 and [$^{18}$F]Lys27, the mice were sacrificed at 60 min ($n$ = 2) and 120 min ($n$ = 2) post-injection for *ex vivo* organ analysis. Radioactivity was measured with a Wizard 2480 gamma counter (PerkinElmer) to determine the percentage injected radioactivity dose per gram of tissue (%ID/g) decay-corrected to the time of injection.

To measure changes in %ID/mL over time in the kidney, liver, spleen and lung, central parts of the organs were outlined on at least five CT slices in the coronal plane registered to the dynamic PET series in HERMIA Hybrid Viewer (v6.1, Hermes Medical Solutions AB, Stockholm, Sweden) for each mouse. These volumes were then transferred to the PET image series and mean activity concentrations within each volume were extracted for each time point. The %ID/mL were then calculated based on the injected radioactivity.

### Tissue homogenisation and fractionation

Organs (lung and spleen) harvested at 60 min post-injection were rinsed with cold PBS (pH 7.4, 100 µL) and then centrifuged at rt (3000 rpm, 3 min). Hypotonic Buffer (200 µL for the lung and 100 µL for the spleen of 10 mM Tris, 15 mM NaCl, 1.5 mM $MgCl_2$, pH 7.6, with Protease Inhibitor Cocktail Roche, cat. no. 11873580001) was added. The tissue was then subjected to 20-35 strokes (35 for the lung and 20 for the spleen) with a loose pestle Dounce homogenizer to lyse the intact cells. The mixture was centrifuged at rt (4000 rpm, 20 min) and the supernatant, designated as the cytoplasm fraction, was removed. The nuclei pellet was washed with cold PBS (100 µL), centrifuged at rt (3500 rpm, 10 min) and the supernatant was discarded.

For SDS-PAGE analysis, the cytoplasm or nuclear fraction were diluted into 4x Laemmli buffer with 20% β-mercaptoethanol such that the final mixture contained 2x Laemmli buffer with 10% β-mercaptoethanol. The samples were heated at 95 °C for 1 h before they were loaded onto a 4-12% Bis-Tris gel. The gels were then exposed to a BAS-TR2025 phosphor imaging plate (Fujifilm, Tokyo, Japan) after gel electrophoresis. After an overnight exposure time, the imaging plates were scanned with a BAS-5000 scanner (Fujifilm, Tokyo, Japan), and the autoradiography images were viewed with the Carimas software (Turku PET Centre, Turku, Finland).

### *Investigation into histone proteolysis*

#### In vitro serum incubation with $^{18}$F-labelled human histones eH3.1

Purified $^{18}$F-labelled histones in PBS were incubated in mouse serum at 37 °C (1:4 dilution). Aliquots were taken various time points and immediately frozen until they were ready to be analysed by HPLC (Method B). For control experiments, purified $^{18}$F-labelled histones in PBS were kept at 37 °C.

#### Stability of histones versus nucleosomes in serum and plasma

A solution containing WT human histone eH3.1 in $H_2O$ with protein concentration 56 μM or nucleosome with a total concentration of 5.4 μM was incubated with PBS, rat serum or plasma at a 1:10 dilution at 37 ˚C. WT histone eH3.1 at 56 μM was also incubated with an equimolar amount of histone H4 in $H_2O$ (final concentration is 28 μM for each histone) for at least 15 min before incubating with serum/plasma at a 1:5 dilution at 37 ˚C. For each different histone form in PBS, serum or plasma, aliquots were taken and diluted into 1x Laemmli buffer with 5% β-mercaptoethanol (βME) at 0 min and then at intervals of 15, 30, 60 and 120 min. These samples were immediately frozen at -80 °C until they could be analysed by Western blot using the HA-tag rabbit mAb as the primary antibody and the alkaline phosphatase-coupled anti-rabbit IgG as the secondary antibody.

#### SDS-PAGE and LC-MS analysis on APC-treated histones

Stocks of histones eH3.1 (WT and $^{19}$F-Lys) were prepared at 25 μM protein concentration in $H_2O$ and then diluted with PBS to a final concentration of 1 μM in 1 mL. The diluted histone H3 solution was kept at 37 ºC for at least 15 min with shaking. APC (ThermoFisher Scientific, #RP-43095) was added to achieve a concentration of 10 nM and the subsequent mixture was incubated at 37 ºC with continuous shaking at 600 rpm (Extended Data Figure 4). Aliquots were taken at various time points for LC-MS analysis or gel electrophoresis.

Stocks of histones eH3.1 (WT and K27M mutant) and H4 in $H_2O$ were individually prepared at 50 μM protein concentration and then mixed at equal volumes such that the concentration of each histone is 25 μM. The equimolar mixture of histones eH3.1 and H4 was incubated at 25 ºC for at least 15 min with shaking and then diluted into PBS to reach a final concentration of 1 μM of each histone (1:25 dilution) in 1 mL. This diluted mixture of histones eH3.1 and H4 was kept at 37 ºC for at least 15 min with shaking. APC was added to achieve a specified concentration and the subsequent mixture was incubated at 37 ºC with continuous shaking at 600 rpm. Aliquots were taken at various time points for LC-MS analysis or gel electrophoresis.

#### *Histone administration into rodent brain and immunohistochemistry*

Sprague Dawley rats (Charles River, UK), 8–10 weeks of age, were housed under standard diurnal lighting conditions (12 h) with ad libitum access to food and water. All procedures were carried out in accord with the UK Animals (Scientific Procedures) Act (1986) and licensed protocols approved by local committees (LERP and ACER, University of Oxford) and were carried out under licence number P996B4A4E in aseptic conditions. For surgery, the animals were anesthetized in a 2% isoflurane/oxygen mix (2 L min$^{-1}$) and placed in a stereotactic frame (Stoetling Co., Wood Dale, IL) under maintenance anaesthesia (2.0%) and in accordance with always ARRIVE guidelines. Following isoflurane-induced anaesthesia, the head of the animals was shaved, sterilized, and mounted on a stereotaxic frame. Under an operating microscope, a small midline scalp incision was made after bupivacaine injection and a burr hole was created using a dental drill at 1 mm anterior and 3 mm lateral to the bregma. One microlitre of histone (1 mg mL$^{-1}$) or vehicle in sterile PBS, was then injected slowly over 5 min via a finely drawn glass microcapillary into the left striatum 4 mm from the cortical surface, before suturing the incision. For control animals, 1 µL of sterile PBS was injected. The animals were allowed to recover from the anaesthesia and were closely monitored throughout and were observed to behave normally before they were returned to their home cage after 5 min. At least 3 animals were used per group in this experiment.

Following the injections, the animals were euthanatized with an intraperitoneal injection of sodium pentobarbitone; the brain was removed at either 30 min, 3 h or 6 h and collected either for chromatin extraction or for immunohistochemistry (IHC) after intracardiac perfusion fixation under terminal anaesthesia with 4% paraformaldehyde. The brains for IHC were fixed for a further 4 h and cryoprotected in 30% sucrose overnight at 4˚C before being embedded in Tissue-Tek (Miles Inc, Elkhart, USA) and quickly frozen in liquid nitrogen. Ten-micron-thick serial frozen sections were cut on a microtome and stained with specific antibodies (Serotec, Oxford, UK) to neutrophils (HB199), microglia (Iba-1) and to astrocytes (GFAP). The FLAG-HA tagged histone eH3.1 was detected with an anti-HA antibody. An avidin-biotin-peroxidase method was employed for all immunohistochemistry[8].

### *Histone incorporation into HeLa cells*

#### Cell culture

Adherent WT HeLa cells (Acc No: 93021013, Lot: 17A001, P: +5) were grown as monolayers in Dulbecco's modified Eagle's medium (DMEM) supplemented with 10% fetal bovine serum (FBS), and L-glutamine (2 mM) in a 37 ºC, 5% $CO_2$ incubator.

#### Cell viability assay

WT HeLa cells at approximately 60-80% confluency on a 96-well plate were washed with PBS and then treated with equimolar WT histones eH3.1 and H4 under a range of specified concentrations and incubation times. The cells were then washed three times with PBS and allowed to recover for 2 h in fresh media. Cytotoxicity assay was carried out using a cell counting kit (WST-8 / CCK8 solution, Abcam, ab228554) following the manufacturer’s protocol. Briefly, WST-8 solution was added to each well and the plate was incubated for 1 hours at 37 ºC in the dark. Absorbance was measured at 460 nm (Agilent BioTek Cytation 1 Cell Imaging Multi-Mode Reader).

#### Double thymidine blocking protocol

HeLa WT cells at ~40% confluency were treated with thymidine blocking solution (PBS stock, final concentration 2 mM thymidine per 1 mL media) and the cells were incubated for 16 h. Cells were then washed two times with PBS, fresh media was added, and cells were allowed to grow for 9 h. The thymidine blocking was repeated, and cells were incubated for a further 14 h. Media was removed, and cells were washed twice with PBS.

### RNA-seq

Double thymidine blocked HeLa cells were incubated in PBS containing an equimolar mixture of WT histones eH3.1 and H4 (1 μM each) for 15 min at 37 °C. After the histone treatment, the cells were washed with PBS and incubated with fresh media for 6 h. Media was then removed, and 350 μL of Trizol reagent was added. Total RNA was purified using the Direct-zol Miniprep Plus Kit (Zymo Research) following the manufacturer's instructions. After an equal volume of ethanol was added to the Trizol reagent mixture and placed on the Zymo-Spin IIICG Column, the flow through containing the total protein extract was collected and stored at -20 °C. Protein was later precipitated by adding 4 volumes of ice-cold acetone and incubating for 30 min. The precipitant was pelleted (13,000 rpm, 5 min, 4 °C), washed with 400 μL of 95% ethanol, and air dried for 5 min. The pellet was then dissolved in 0.5 mL Sample Buffer (NuPAGE LDS Sample buffer containing 10% βME) by shaking at 37 °C for 30 min. It was then stored at -20 °C for SDS-PAGE and Western blot analysis.

The mRNA was enriched from the total RNA using NEBNext Poly(A) mRNA Magnetic Isolation Module (NEB #E7490) following manufacturer's instructions. Libraries were prepared with the NEBNext Ultra II Directional RNA Library Prep Kit for Illumina (NEB #E7760). Libraries were then multiplexed, quantified using a High-sensitivity d1000 TapeStation (Agilent), then sequenced using a NextSeq 500 (Illumina).

The raw FASTQ files were processed using the featurecounts workflow (https://github.com/cribbslab/cribbslab).[9] Read quality was evaluated with FASTQC and ReadQC. Reads were aligned to the GRCh38 reference genome using HiSat2 (v2.0.5).[10] Quantification of the mapped reads against the GRCh38 genome annotation was performed using FeatureCounts (v1.5.0).[11] Subsequent analyses were conducted in R version 3.5.1 and RStudio version 1.1.456. Differential expression analysis was performed using the DESeq2 package using wald test.[12]

#### HeLa cell histone incorporation protocol

WT HeLa cells were cultured until they reached 60-80% confluency on 10 cm plates. They were washed with PBS, then incubated with a 1:1 molar ratio of WT histone H4 and the respective histone eH3.1 sample, each at 1 µM, for 15 min in PBS in a 5% $CO_2$ 37 °C incubator. The cells were then washed three times with PBS and allowed to recover for variable times (experiment dependent) in fresh media before being washed twice with PBS and harvested with a cell scraper. The cells were centrifuged in PBS for 5 min 800 g before being placed on ice and proceeding to cell fractionation.

#### HeLa cells fractionation

All fractionation steps were carried out either on ice or at 4 °C in a cold room and all fractions were stored at -80 °C. Approximately 6 – 8 x $10^6$ HeLa cells were resuspended in Hypotonic Buffer (500 µL of 10 mM Tris base, 15 mM sodium chloride, 1.5 mM $MgCl_2$, pH 7.6, with Protease Inhibitor Cocktail Roche), and allowed to lyse for 15 min. Nuclei were pelleted by centrifugation at 400 g for 5 min and the supernatant, designated the cytoplasm fraction, was removed. Nuclei were resuspended again in Hypotonic Buffer (500 µL) and subjected to 10 strokes with a loose pestle Dounce homogenizer to lyse residual intact cells and cytoplasmic organelles before repeating the above centrifugation and removing the supernatant. The nuclei pellet was then resuspended in Hypotonic Buffer containing 1% Triton X-100 (500 µL) and lysed with 10 strokes from a tight pestle Dounce homogenizer. The nuclear lysate fraction was separated from the chromatin fraction by pelleting the latter by centrifugation at 10,000 g for 10 min. The chromatin pellet was washed once with Hypotonic Buffer containing 1% Triton X-100 (500 µL), resuspended in RIPA-SDS Lysis Buffer (500 µL RIPA buffer + 1% SDS) and then sonicated using a Bioruptor Pico (Diageode, 30s on/30s off, 15 min).

#### Chromatin extraction from HeLa cells and rodent tissue

Chromatin was extracted according to the Chromatin Extraction Kit-Flexible Format (Abcam, ab223876). Briefly, HeLa cells were collected by scraping, washed with PBS, crosslinked with 1% formaldehyde (15 min in PBS), and quenched with the kit provided formaldehyde quenching solution (5 min in PBS) before 2x PBS washes. Crosslinked HeLa cells was resuspended in 500 µL of Extraction and Lysis Buffer supplemented with protease inhibitors, then treated with 10 passes of both the loose and tight Dounce homogenizer settings, before being incubated for 30 min at RT with 1000 rpm shaking. Chromatin pellets were spun down (10,000 rpm, 15 min) and washed once with milliQ water. Chromatin was then sheared by sonication with a Bioruptor Pico (30s on/30s off, 15 min), insoluble material was spun down (10,000 g, 10 min) and soluble sheared chromatin was stored at -80 °C.

For SDS-PAGE analysis, the lysate and soluble chromatin samples were diluted with 4x Laemmli buffer supplemented containing 20% βME and then heated at 95 °C for at least 1 h.

#### *Histone incorporation into rodent tissue*

Rodent tissue samples were diced with a scalpel, washed twice with ice-cold PBS, crosslinked in 1% formaldehyde (15 min in PBS) with frequent vortexing, and quenched with the kit provided formaldehyde quenching solution (Abcam, ab223876; 5 min in PBS) before 2x PBS washes. Crosslinked HeLa cells or rodent tissue was resuspended in 500 µL of Extraction and Lysis Buffer supplemented with protease inhibitors, then treated with 10 passes of both the loose and tight Dounce homogenizer settings, before being incubated for 30 min at RT with 1000 rpm shaking. Chromatin pellets were spun down (10,000 rpm, 15 min) and washed once with milliQ water. Chromatin was then sheared by sonication with a Bioruptor Pico (30s on/30s off, 30 min for tissue), insoluble material was spun down (10,000 g, 10 min) and soluble sheared chromatin was stored at -80 °C.

##### Immunoprecipitation for ChIP-seq and proteomics analysis

Immunoprecipitation (IP) Buffer (from Chromatin Extraction Kit, ab223876) was added to the sonicated chromatin (200 µL of sonicated chromatin was added to 2 mL of 1x IP Buffer) followed by 2 µL of A:G Dynabeads (pre-rinsed at least three times with IP Buffer). The mixture was incubated at room temperature for 30 min with mixing at 900 rpm before the supernatant was removed after centrifugation (1 min, 1000 g). To reduce background signal, further addition and incubation with A:G Dynabeads (2 μL) was performed. The mixture was centrifuged and the supernatant collected. Aliquots of the supernatant were taken (2 x 50 μL) for inputs. To 1 mL of cleared chromatin, α-HA (132 ng) or α-H3 (248 ng) antibody was added, and the solution was incubated overnight at 4 °C. A:G Dynabeads (10 μL) were then added and mixed for 5 h at 900 rpm. The supernatant was removed on a magnetic stand. The beads were then resuspended and washed twice with RIPA (radioimmunoprecipitation assay) wash buffer (50 mM HEPES, pH 7.6, 500 mM LiCl, 1 mM EDTA, 1% NP-40, 0.7% Na-Deoxycholate) followed by TE (Tris-EDTA) buffer containing 50 mM NaCl (1 mL). The bead solutions were centrifuged (3 min, 1000 g) to remove the TE buffer. The chromatin IP was eluted from the beads with 210 μL of elution buffer (50 mM Tris-HCl, pH 8.0, 10 mM EDTA, 1% SDS; made fresh) and incubation at 65 °C for 30 min at 900 rpm. The samples were centrifuged (1 min, 16000 g) at room temperature and <200 μL of the desired supernatant was collected. The chromatin supernatant was de-crosslinked overnight at 65 °C. Input samples were diluted with elution buffer (1:4) and similarly, incubated at 65 °C overnight.

#### DNA extraction for ChIP-seq

To the decrosslinked chromatin samples, RNase (1 μL) was added and the mixture was incubated for 1 h at 37 °C (500 rpm). This was then followed by addition of Proteinase K and incubation for 1 h at 45 °C (500 rpm). The DNA was recovered by Qiagen PCR purification kit using standard protocol. Qiagen spin columns were used to bind to the DNA. The DNA was then eluted with 50 – 200 μL of $H_2O$.

#### Adapter ligation, PCR amplification and DNA purification for Illumina sequencing

The DNA was prepared for sequencing using the NEBNext® Ultra™ II DNA Library Prep Kit for Illumina® (New England Biolabs, E7645) following the manufacturer’s instructions. Briefly, ChIP DNA (~10 ng in 50 μL TE buffer; concentration quantified using a High-sensitivity d1000 TapeStation (Agilent)) were end-repaired, A-tailed and ligated using reagents provided in the kit. Cleanup of the ligation reaction was performed without size selection. The ligated DNA fragments were amplified using indexed primers (Illumina) for 6 PCR cycles and purified with 0.9 x volume NEBNext Sample Purification Beads. Size distribution of the DNA libraries were checked and sequenced using NextSeq 500.

The ChIP-seq data was processed using the following steps: Raw FASTQ files were mapped to the rat genome (rn6) using Bowtie2 (v2.4.5). The resulting SAM files were sorted and indexed with Samtools (v1.15.1). The bigwig files were generated using deepTools (v3.5.1). Peak calling was conducted using MACS2 (v2.2.7.1) with an effective genome size of 2.75 billion, a q-value cutoff of 0.01, with the setting either broad or sharp peak calling. Finally, peak annotation was performed with HOMER.

#### Tandem mass spectrometry and data analysis

16 µL of IP eluate and sheared chromatin were loaded onto the 12% Bis-Tris gel. Using pre-stained marker and SafeBlue stained histone bands in sheared chromatin small bands were excised for propionylation. Cut gel bands were washed with water and cut to approx. 1 $mm^3$ cubes and derivatised with propionic anhydride according to a published protocol[13]. After derivatisation, gel pieces were covered with digestion buffer containing 30 ng trypsin (Promega Gold) and 50 mM ammonium bicarbonate in ultra-pure water. Analysis of peptides was carried out using an Ultimate 3000 nano-LC 1000 system coupled to an Orbitrap Ascend (Thermo Fisher Scientific). Supernatant from in-gel digestion was diluted 1:1 in ultra-pure water with 5% formic acid and 5% DMSO. Peptides were initially trapped on a C18 PepMap100 pre-column (300 µm inner diameter x 5 mm, 100 Å) and then separated on an in-house constructed C18 column (Reprosil-Gold, Dr. Maisch, 1.9 µm particle size) column (ID: 50 µm, length: 50 cm) at a flow rate of 100 nL/min. Peptides were separated over 15 min (12-38%B) using mobile phase A (water and 0.1% formic acid) and mobile phase B (acetonitrile and 0.1% formic acid). Separated peptides were directly electrosprayed into an Orbitrap Ascend mass spectrometer (Thermo Fisher Scientific). Mass spectra were acquired in the orbitrap (350-1400 m/z, resolution 60000, AGC target 3 x 106, maximum injection time 50 ms) in a data-dependent mode. The top 40 most abundant peaks in the

survey scan were fragmented using CID (resolution 7500, AGC target 4 x104, maximum injection time 64 ms).

Peptide identification and quantification were performed using MSFragger[14] or Andromeda search engine implemented in MaxQuant (2.3.0.0)[15]. Spectra were searched against a reference database (uniport *Rattus norvegicus*, downloaded 25.08.2022) with additional sequence of recombinant H3-FLAG-HA. Default settings were used apart from the following additional variable modifications: propionylation of lysine and protein termini (Da), acetylation of lysine and protein N-termini (Da), mono- di- and tri-methylation on lysine. No fixed modifications were applied. PDV viewer from within FragPipe was used for spectrum annotation.

### *Small molecule NMR spectra*

$^1H$ and $^{13}C$ NMR Spectra of 2-((bromomethyl)thio)benzothiazole

(all taken in $CDCl_3$ unless where stated otherwise)

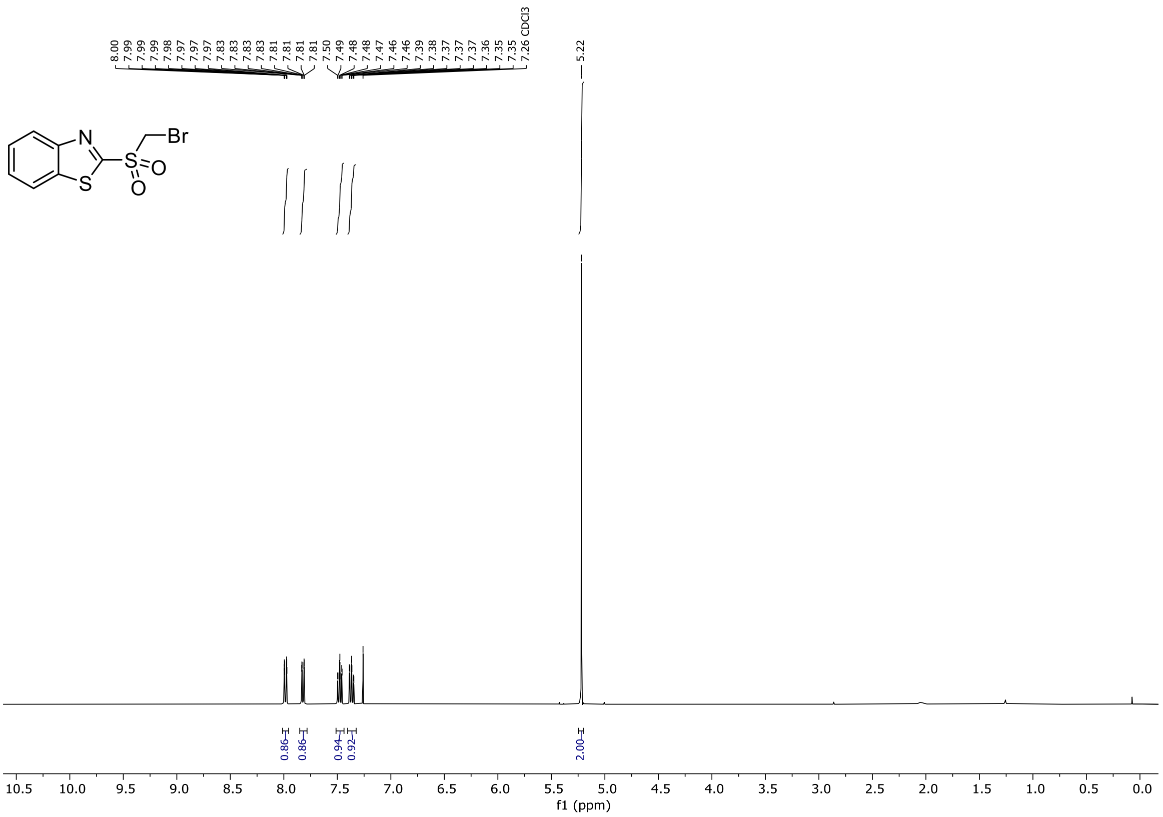

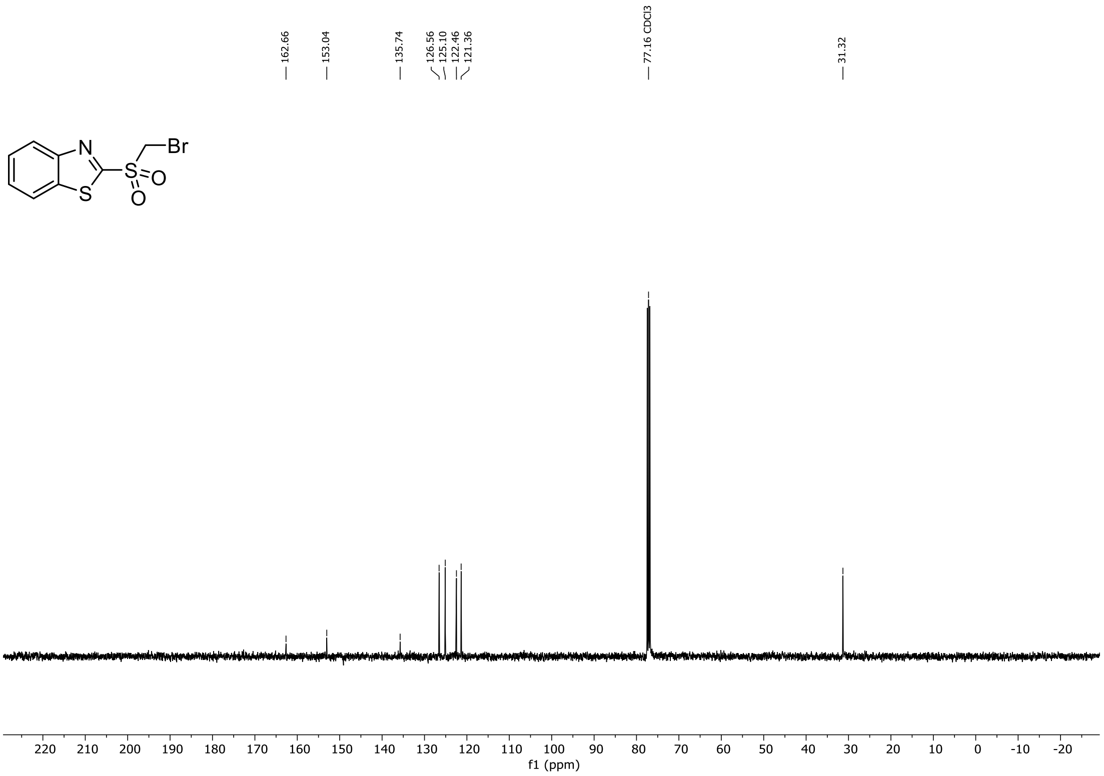

$^1$H and $^{19}$F NMR Spectra of ethyl 2-(benzothiazole-2-ylthio)-2-fluoroacetate

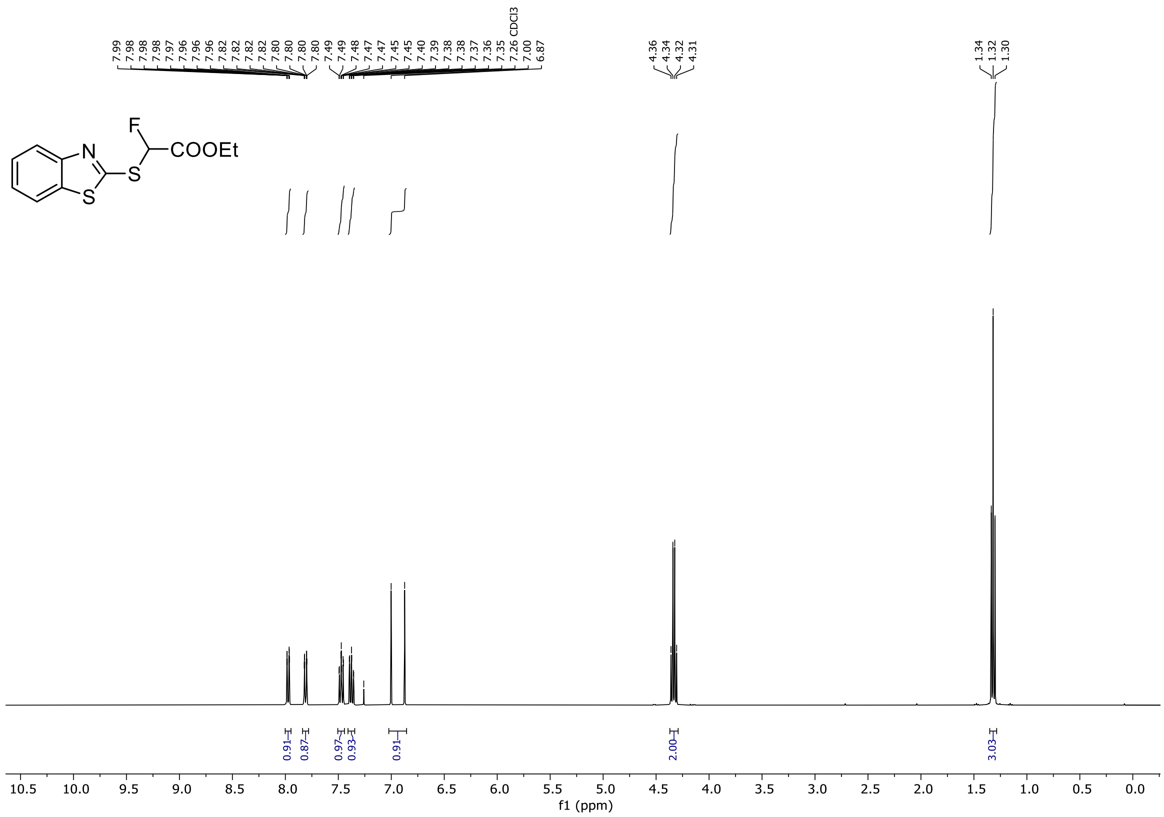

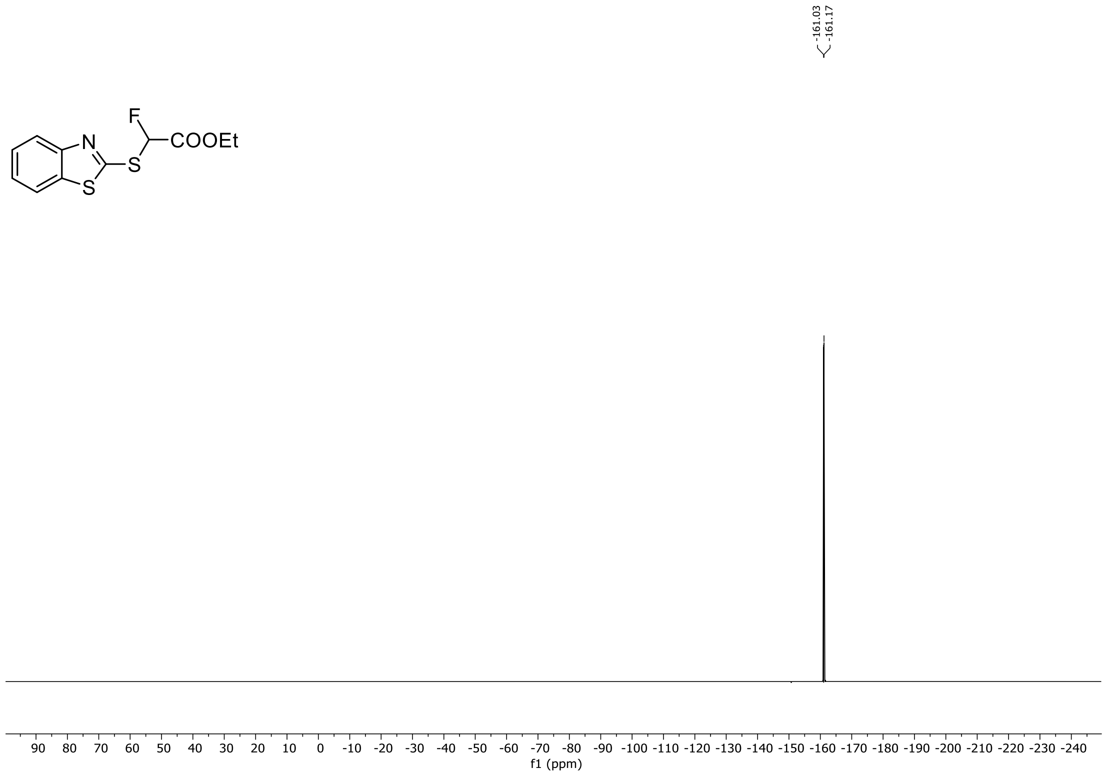

[1]H and [19]F NMR Spectra of ethyl 2-(benzothiazole-2-ylsulfonyl)-2-fluoroacetate

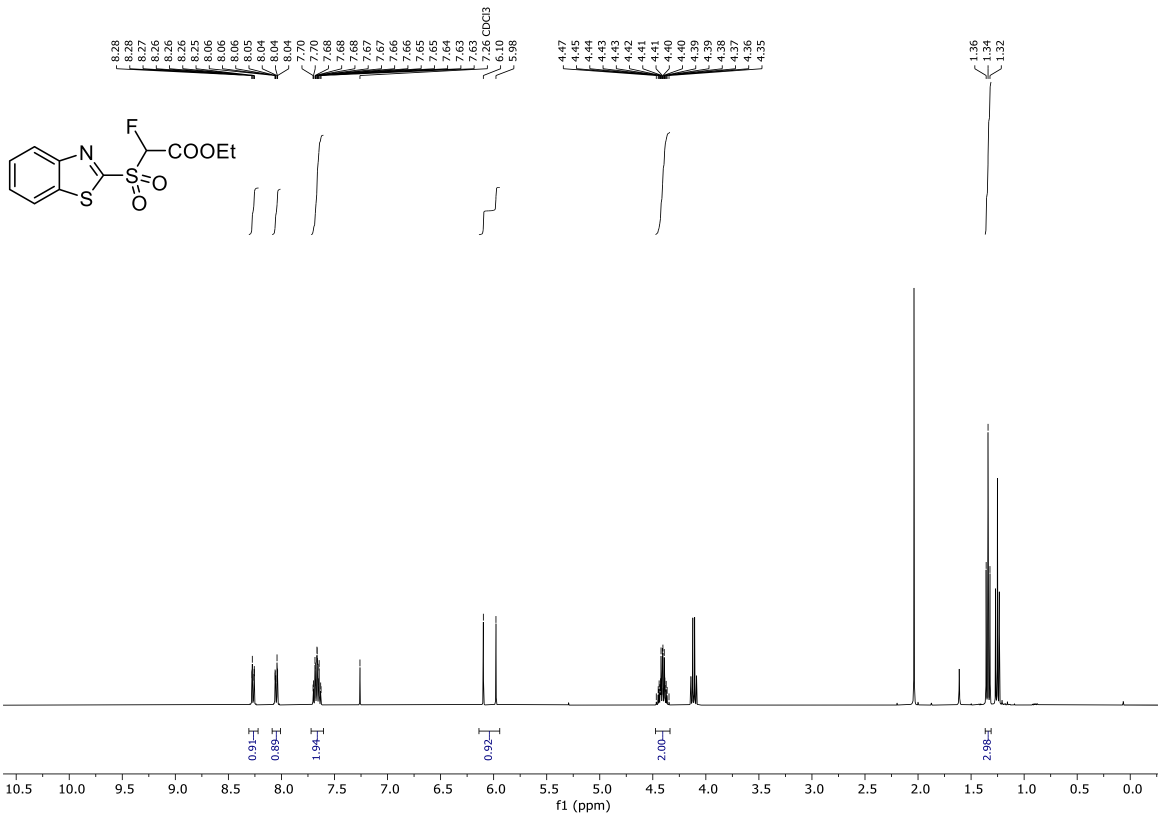

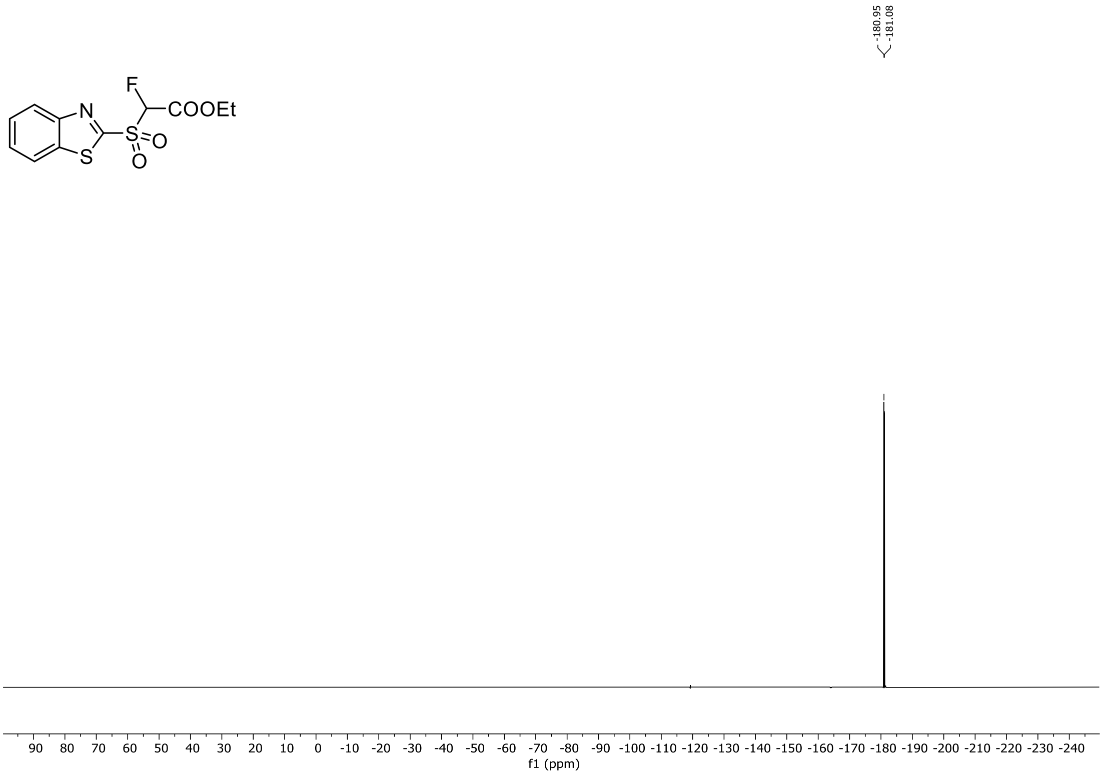

<sup></sup>[1]H and [19]F NMR Spectra of 2-((fluoromethyl)sulfonyl)benzothiazole (all taken in DMSO-$d_6$)

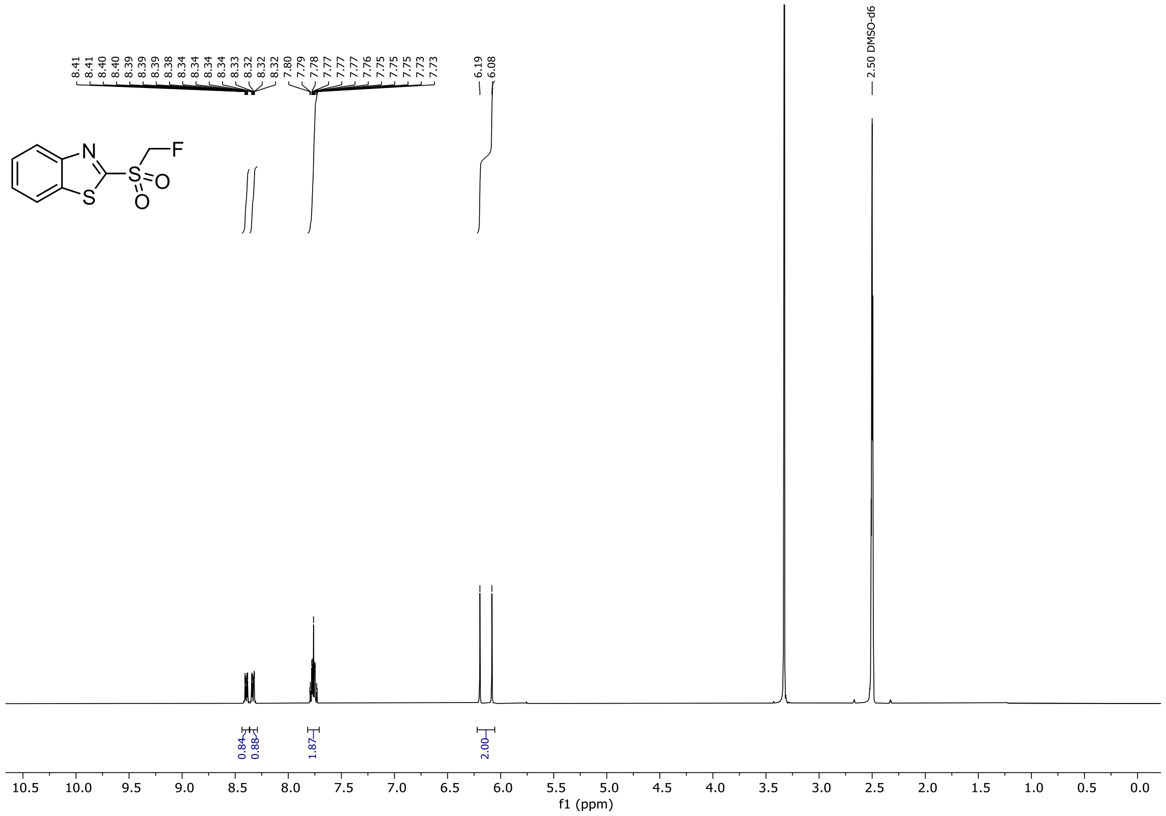

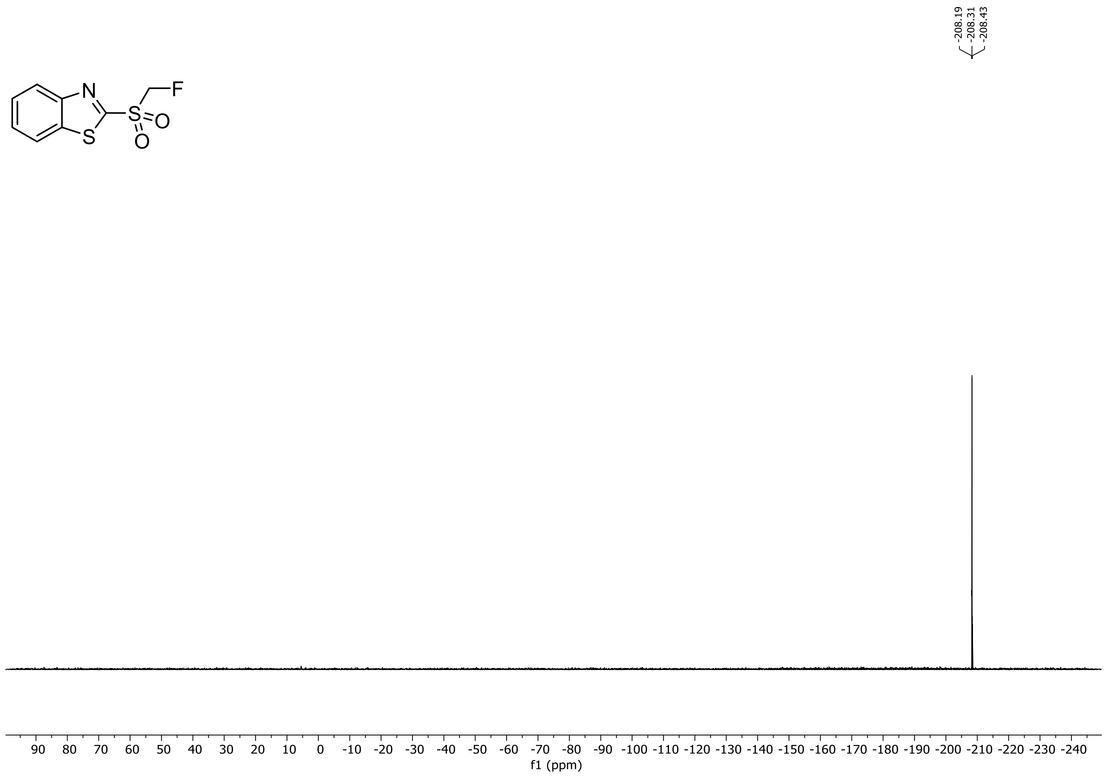

$^{1}H$ and $^{13}C$ NMR Spectra of tert-Butyl (3-(benzothiazol-2-ylthio)propyl)(tert-butoxycarbonyl)carbamate

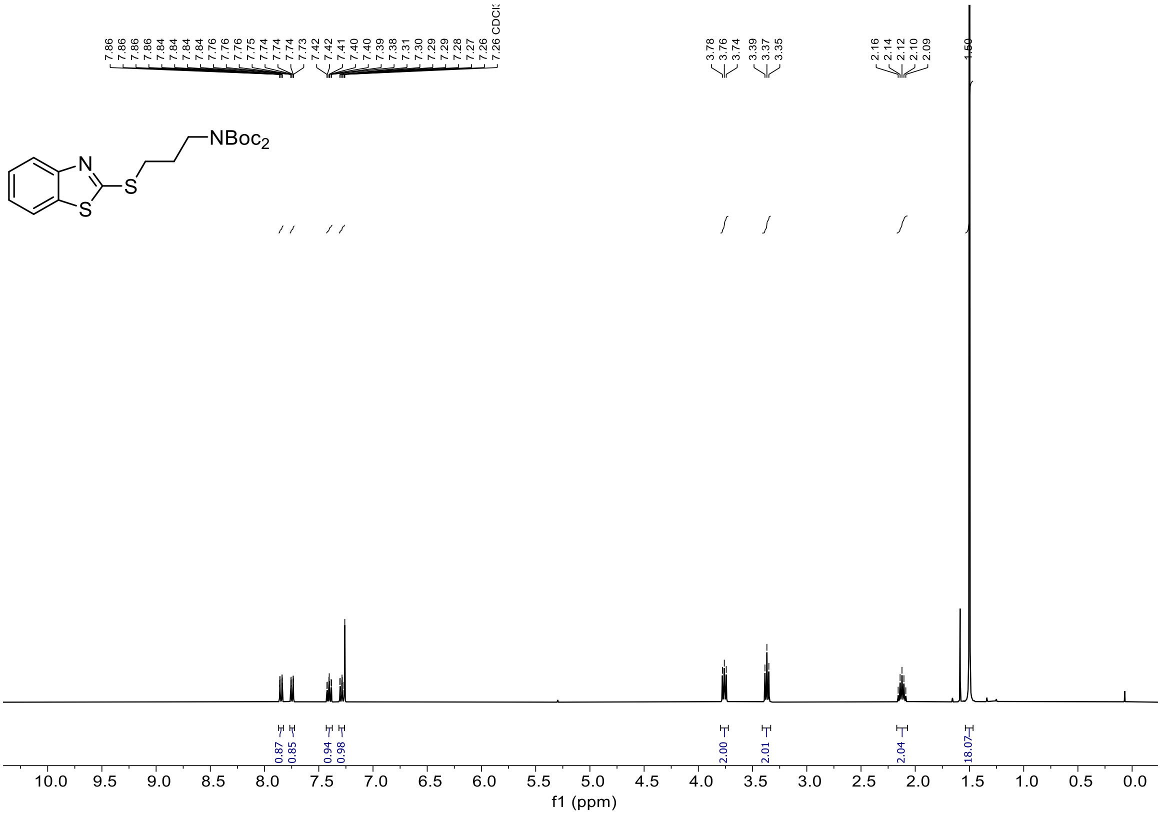

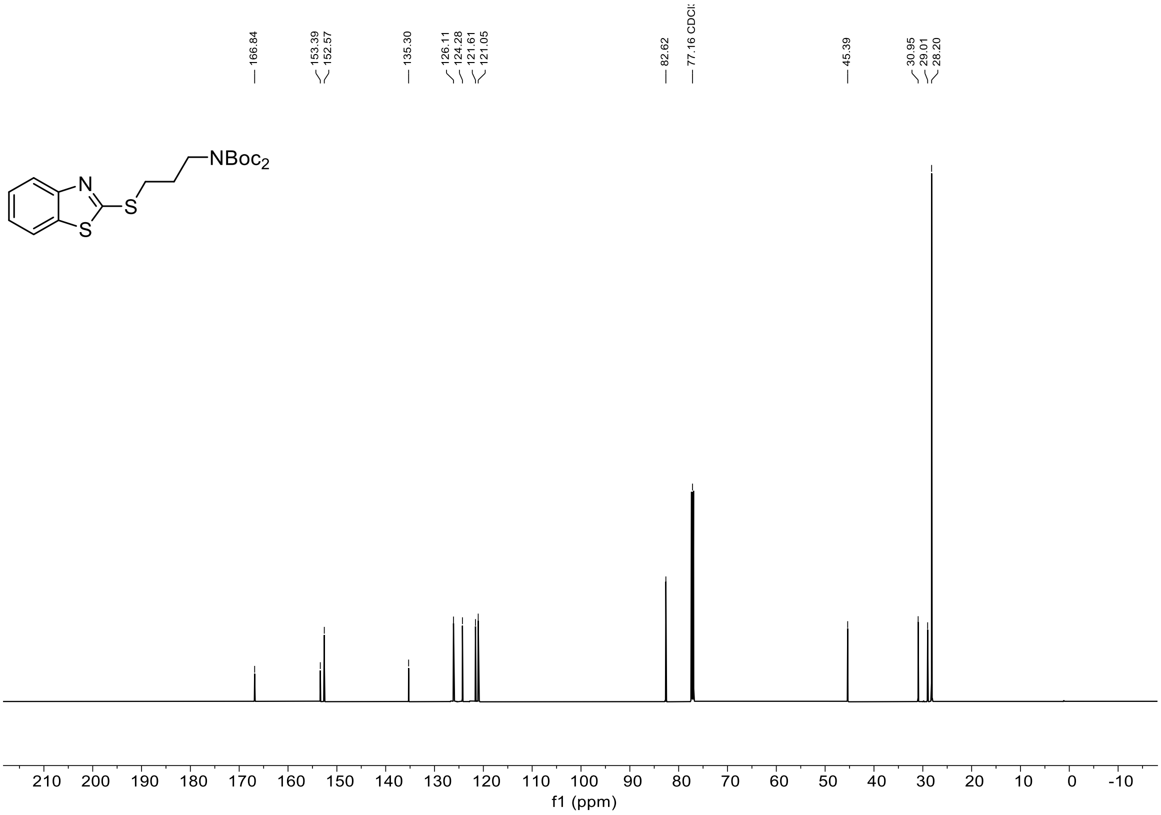

$^1$H and $^{13}$C NMR Spectra of tert-Butyl (3-(benzothiazol-2-ylthio)-3-chloropropyl)(tert-butoxycarbonyl)carbamate

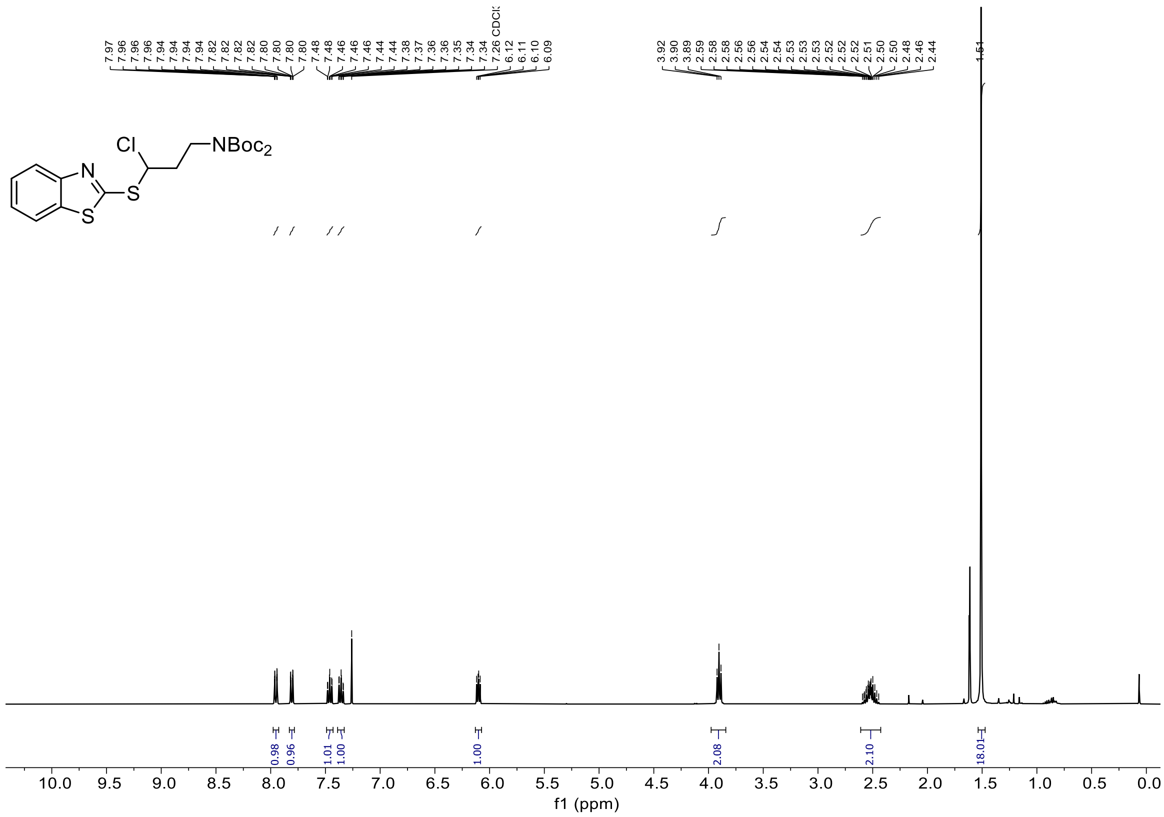

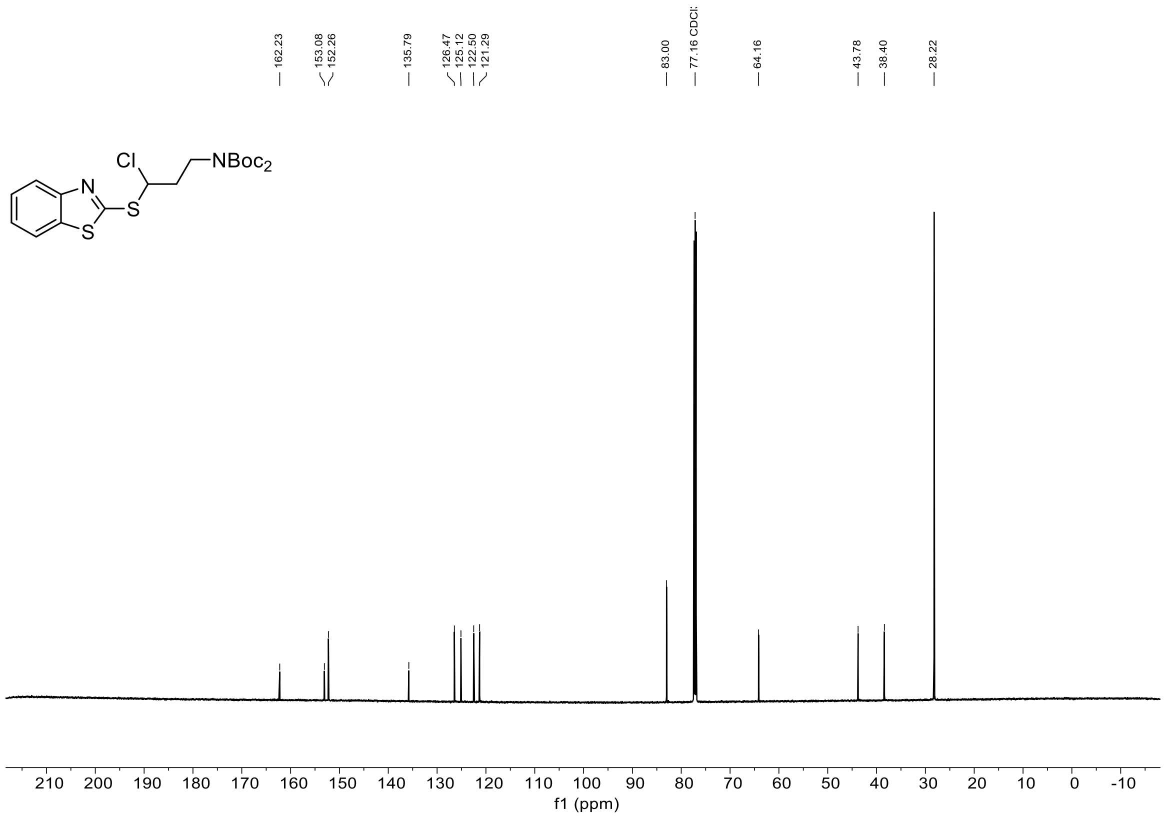

$^{1}H$ and $^{13}C$ NMR Spectra of tert-Butyl (3-(benzothiazol-2-ylsulfonyl)propyl)carbamate

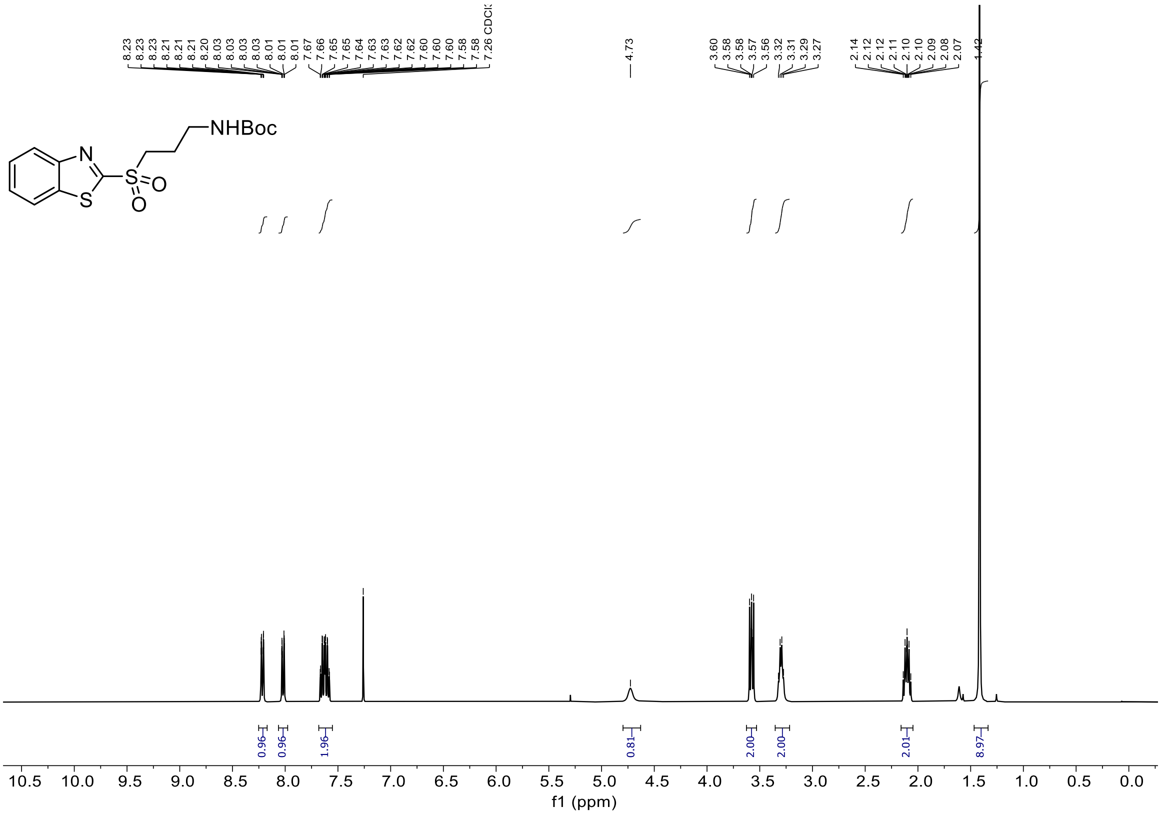

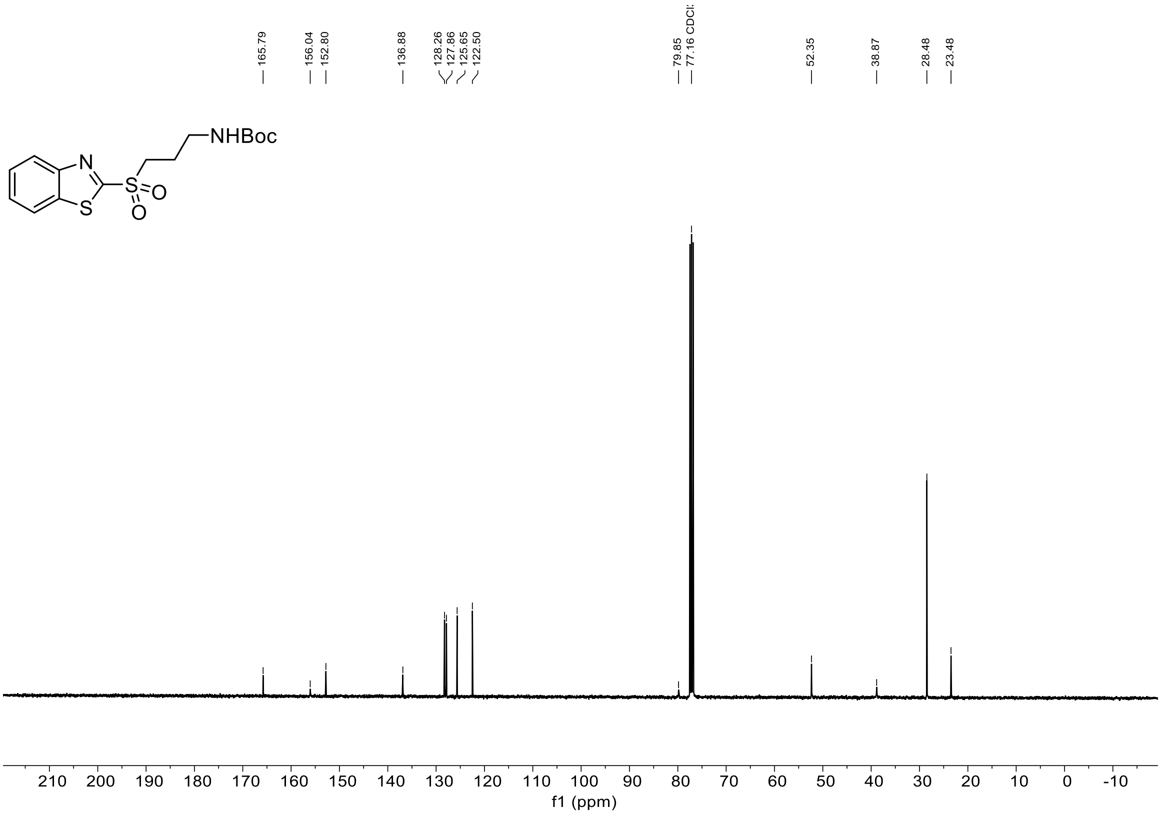

$^{1}H$, $^{19}F\{^{1}H\}$ and $^{13}C$ NMR Spectra of tert-Butyl (3-(benzothiazol-2-ylsulfonyl)-3-fluoropropyl)carbamate

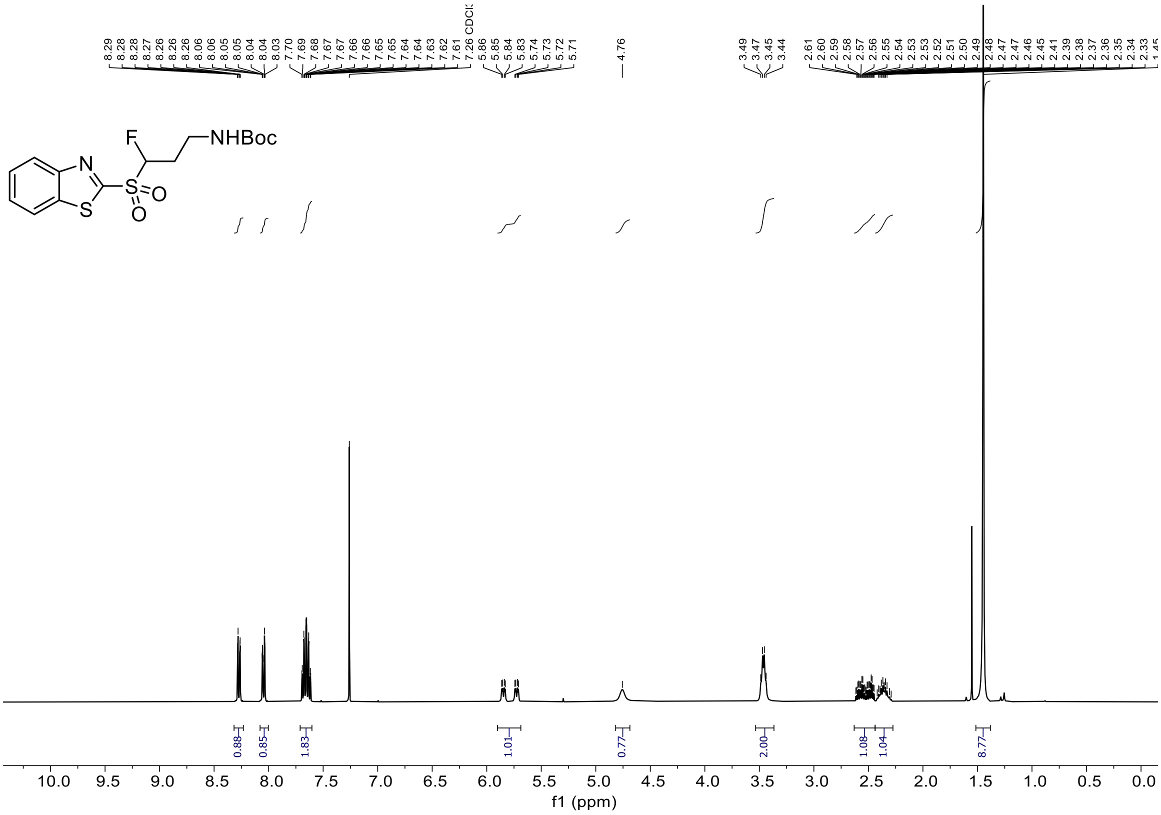

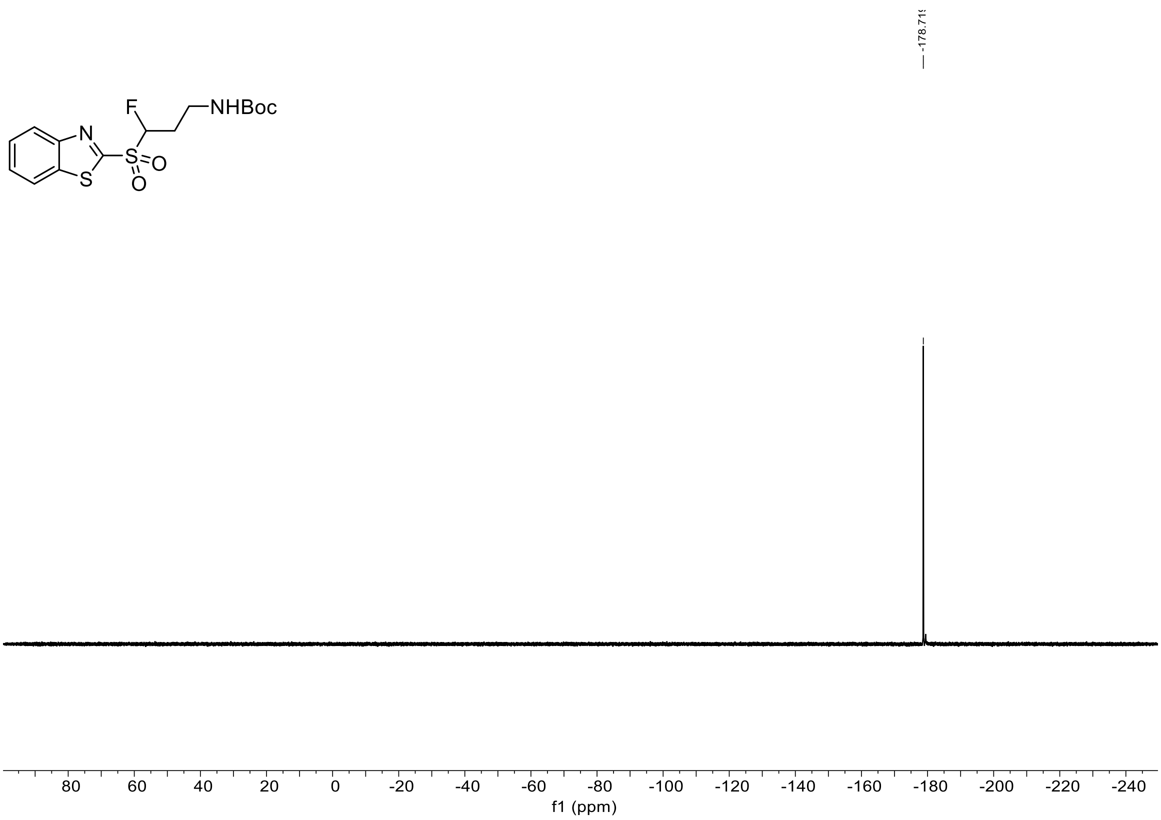

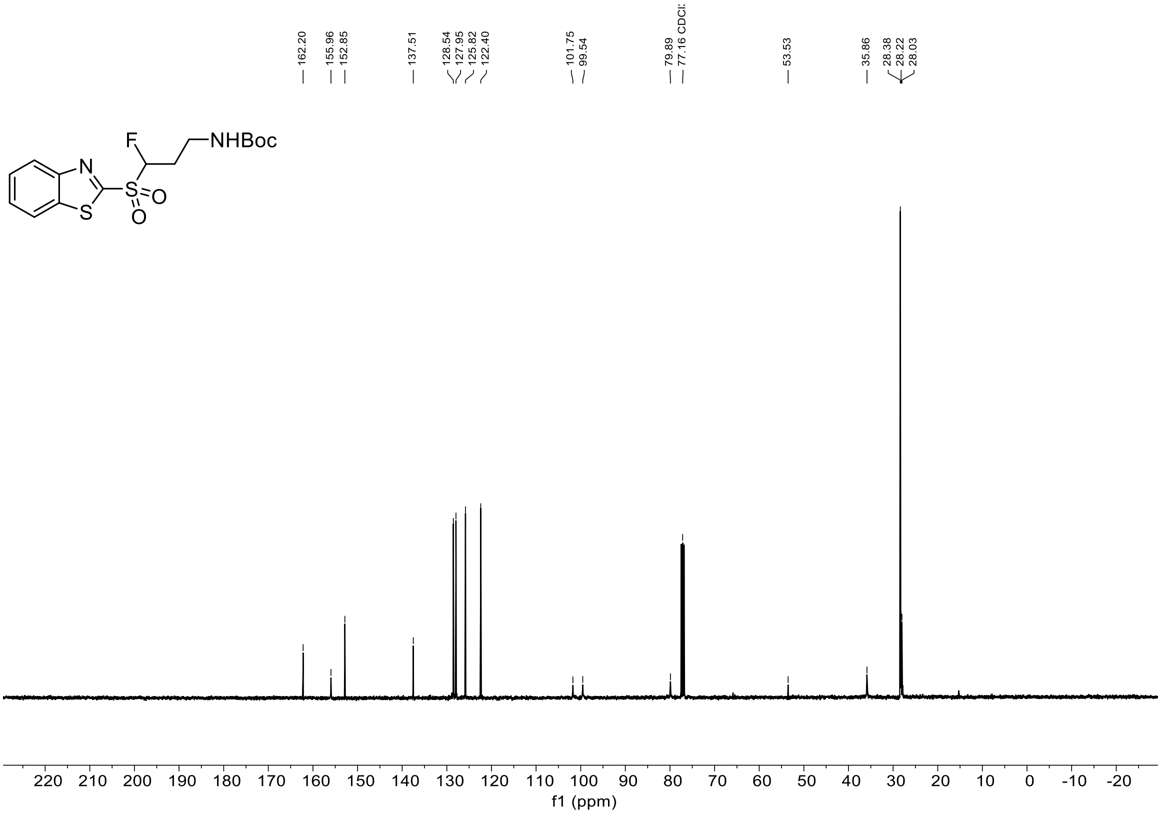

162.20
155.96
152.85
137.51
128.54
127.95
125.82
122.40
101.75
99.54
79.89
77.16 CDCl3
53.53
35.86
28.38
28.22
28.03
F
NHBoc
N
S
S
O
O
220 210 200 190 180 170 160 150 140 130 120 110 100 90 80 70 60 50 40 30 20 10 0 -10 -20
f1 (ppm)

[1]H, [19]F{[1]H} and [13]C NMR Spectra of tert-Butyl 3-(benzothiazol-2-ylsulfonyl)-3-fluoropropan-1-amine hydrochloride (all taken in $CD_3OD$)

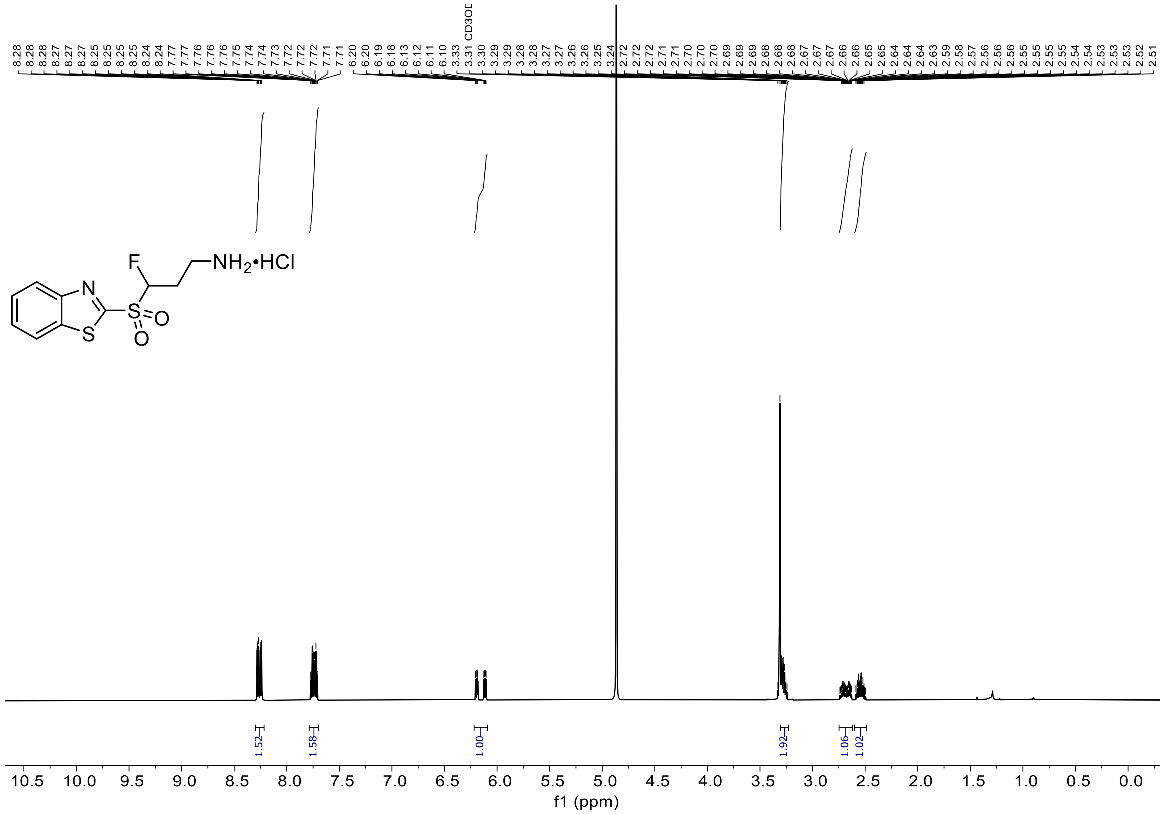

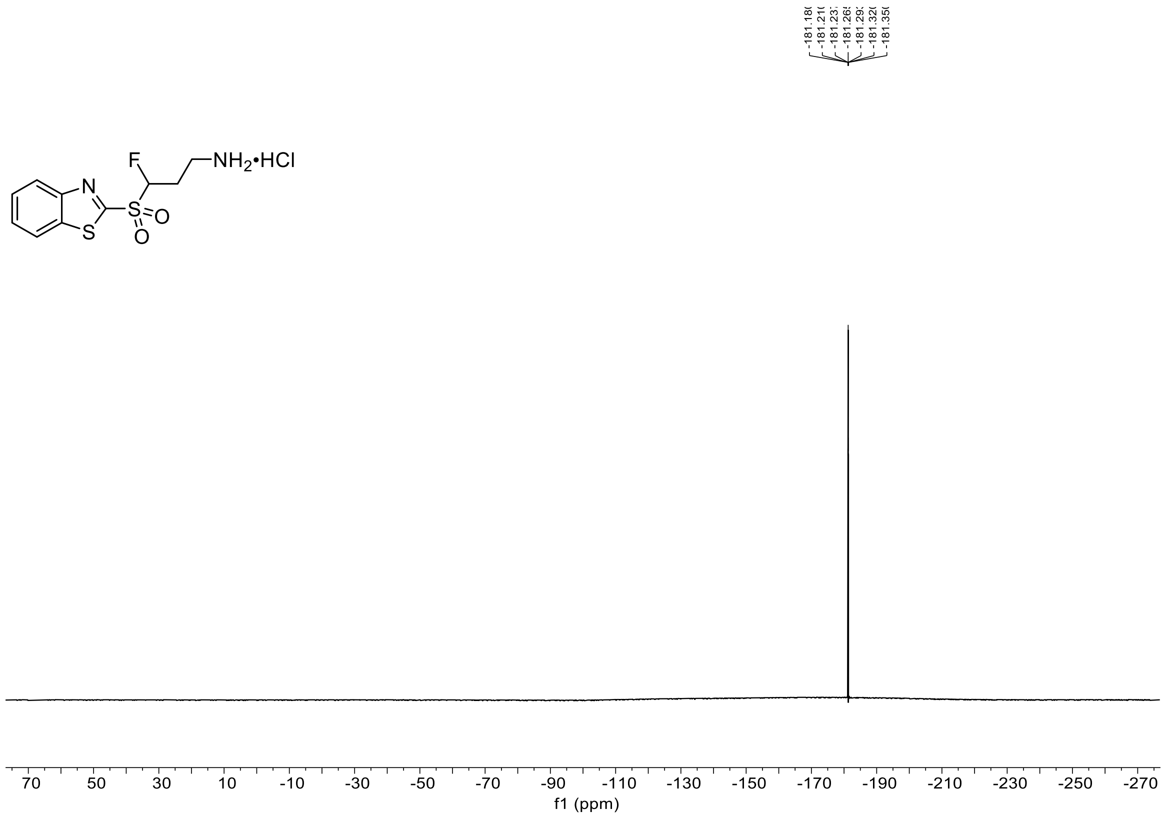

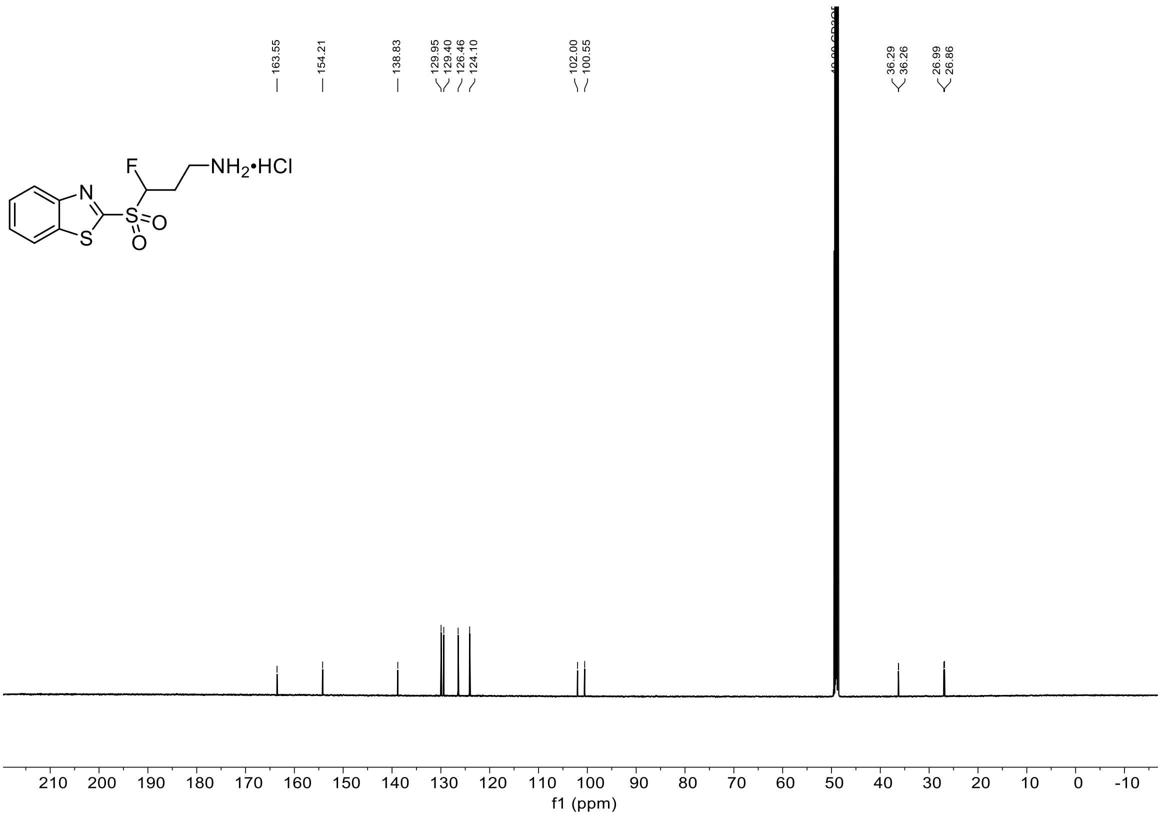
163.55
154.21
138.83
129.95
129.40
126.46
124.10
102.00
100.55
36.29
36.26
26.99
26.86
F
NH2•HCl
N
S
S
O
O
210 200 190 180 170 160 150 140 130 120 110 100 90 80 70 60 50 40 30 20 10 0 -10
f1 (ppm)

$^{1}H$, $^{19}F\{^{1}H\}$ and $^{13}C$ NMR Spectra of tert-Butyl (3-(benzothiazol-2-ylsulfonyl)-3-fluoropropyl)(tert-butoxycarbonyl)carbamate

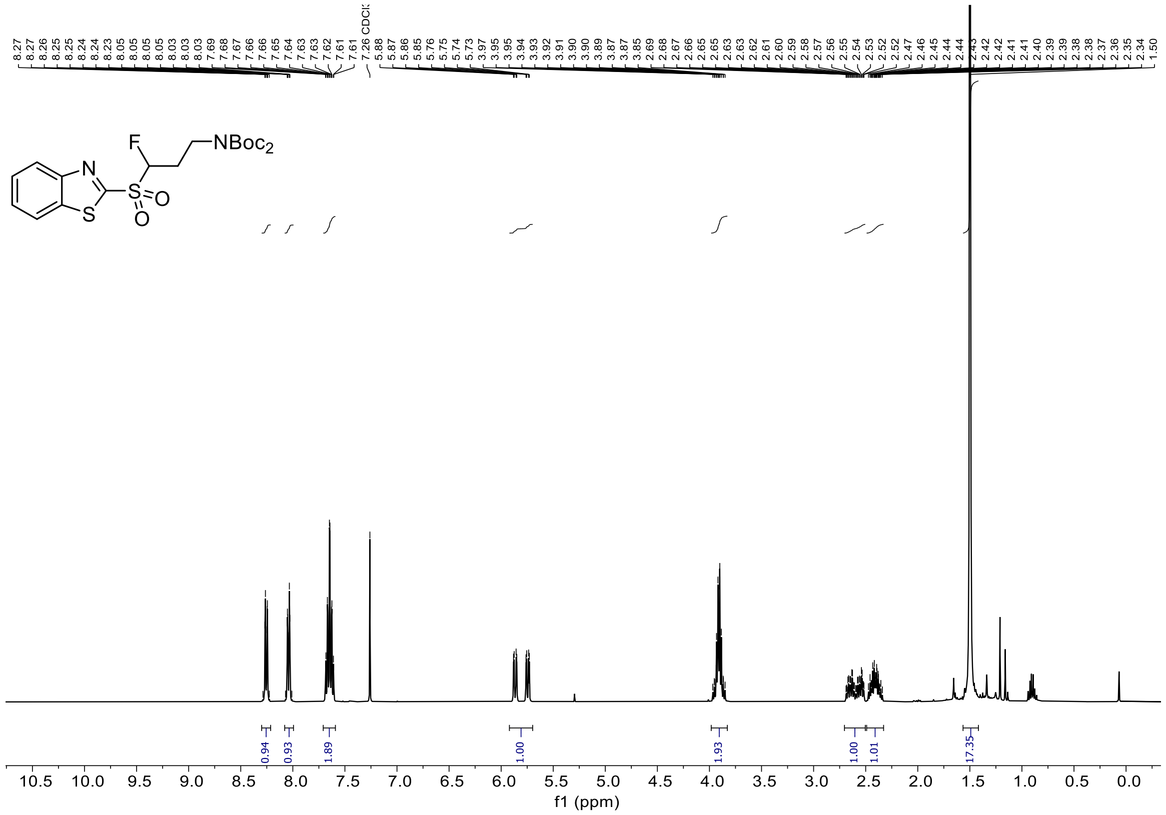

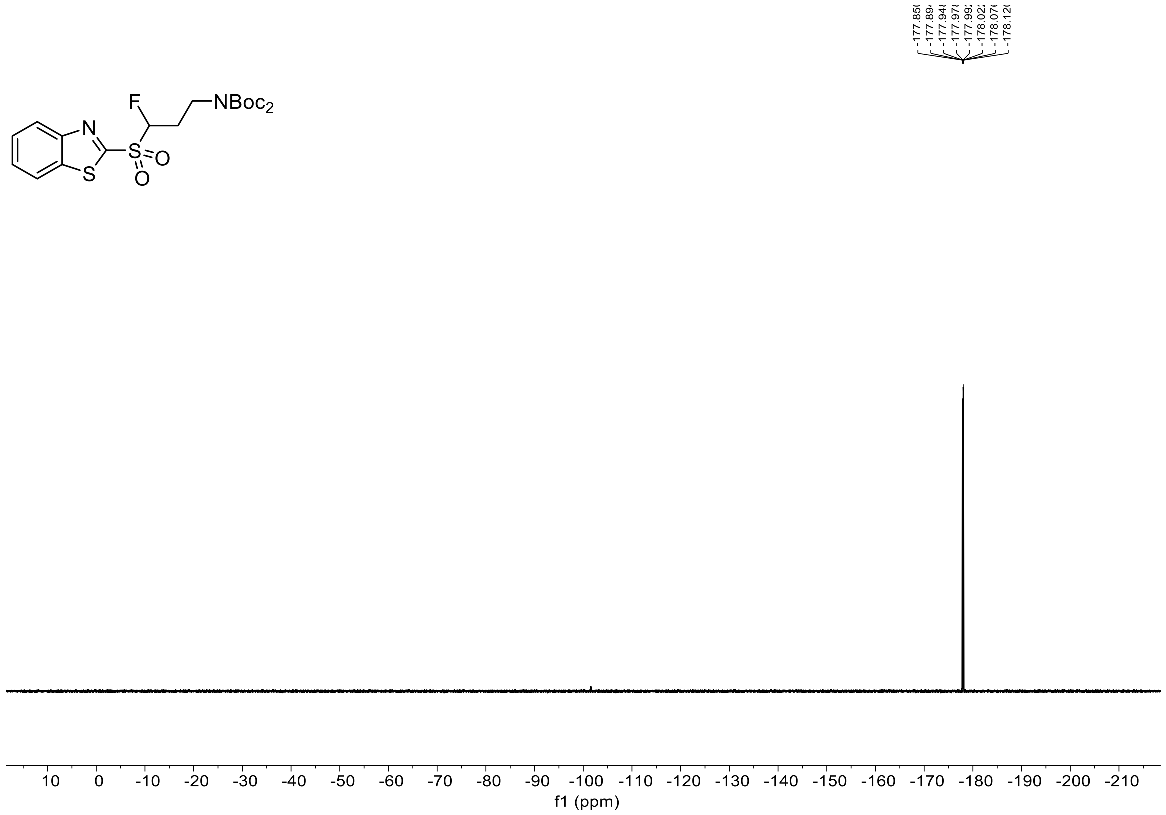

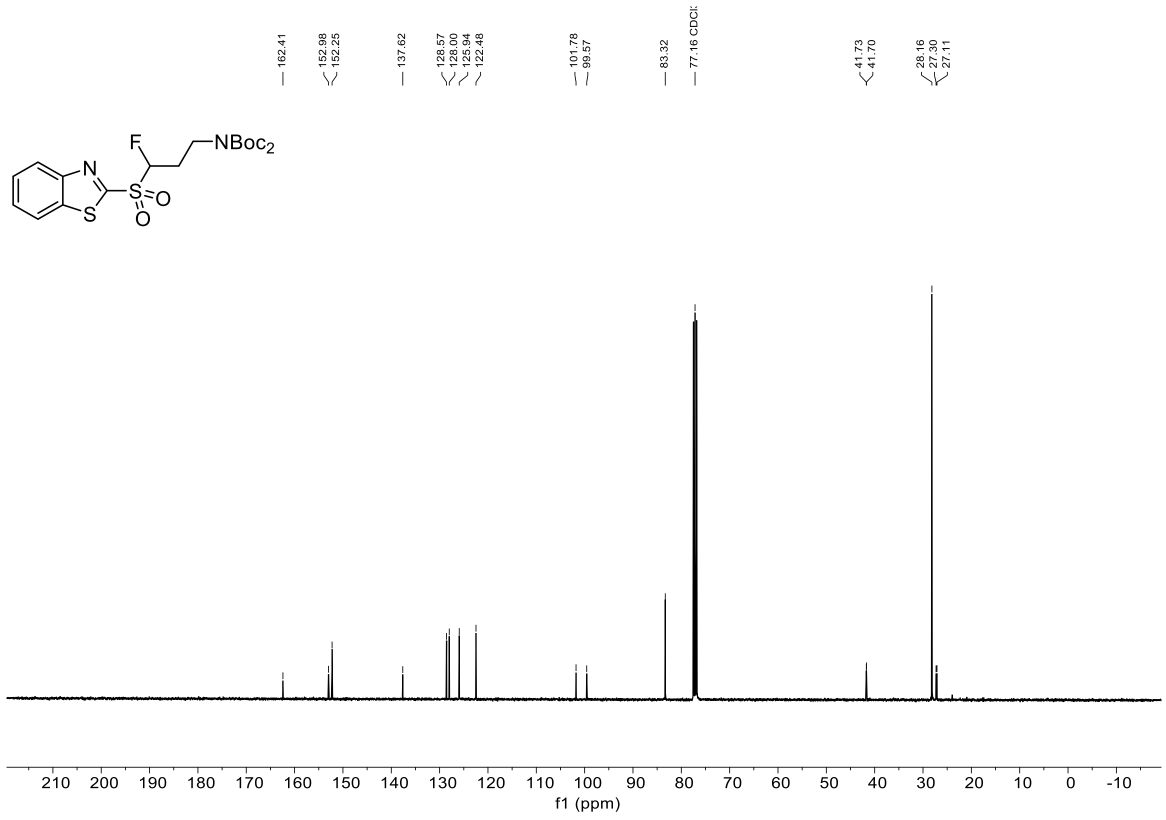
162.41
152.98
152.25
137.62
128.57
128.00
125.94
122.48
101.78
99.57
83.32
77.16 CDCl3
41.73
41.70
28.16
27.30
27.11
F
NBoc2
N
S
S
O
O
210 200 190 180 170 160 150 140 130 120 110 100 90 80 70 60 50 40 30 20 10 0 -10
f1 (ppm)
